# Coupled Cell-Intrinsic and Microenvironmental Heterogeneity Drives Divergent Trajectories in Castration-Resistant Prostate Cancer

**DOI:** 10.64898/2026.07.16.738502

**Authors:** Sharvari Kemkar, Mengdi Tao, Alokendra Ghosh, Adarsh Ramamurthy, Ravi Radhakrishnan

## Abstract

Castration-resistant prostate cancer emerges from coupling between cell-intrinsic heterogeneity and microenvironmental constraints. Mechanistically dissecting this coupling, rather than either factor in isolation, is the central aim of this study. To systematically study the effect of intrinsic and extrinsic spatial axes on disease trajectories, we developed an integrated multiscale framework: a cellular signaling model (MHS) parameterized with TCGA genomic data from control, biochemical recurrence (BR), and treatment-resistant (TR) cohorts, coupled to a spatial agent-based model (ABM). Machine learning surrogates trained on the MHS model identified PTEN, MDM4, and AR as dominant intrinsic drivers, using SHAP-based feature ranking. Clinical validation via Kaplan-Meier and Cox regression across cBioPortal cohorts confirmed these rankings: AR alterations (median OS 20 vs. 86 months), PTEN loss (54 vs. 77 months), and MDM4 amplification (33 vs. 75 months) predicted poor overall survival outcomes independently. At the tissue scale, ABM simulations were run to study the effect of microenvironmental (cell-extrinsic) factors such as physical confinement, androgen uptake kinetics, and adhesion-motility strength, on disease progression. This spatiotemporal analysis revealed that identical genetic alterations produce varied selection outcomes depending on microenvironmental context. Extending this coupling logic to the individual patient level, we parameterized the MHS model using gene expression profiles from 14 patients in the EUREKA1 prospective registry. Patient-specific androgen sensitivity ratios, the ratio of net cell growth under high versus low testosterone, stratified patients by model-predicted androgen dependence without requiring longitudinal PSA observations and showed that PTEN deletion shifts the proliferative response toward androgen independence in a patient-specific magnitude set by the broader expression background. This provides a model-based route from genomic data at diagnosis to personalized prediction of ADT resistance risk. Together, these findings establish that CRPC emergence is an emergent property of the intrinsic-extrinsic coupling, that neither molecular nor spatial analyses in isolation can predict clinical trajectories, and that mechanistic integration of both is required for accurate patient stratification.

## Introduction

Prostate cancer is the second leading cause of cancer-related death among men in the United States, and its clinical management is complicated by profound biological heterogeneity that current risk stratification tools fail to capture.^1^ The disease progresses through multiple stages, from benign prostatic intraepithelial neoplasia to localized prostate cancer and, in a significant fraction of patients, to advanced metastatic disease. ^2^ Although radical prostatectomy (RP) is the standard of care for localized disease, approximately 20 - 30% of patients experience biochemical recurrence (BR) within five years, defined by a detectable post-surgical serum PSA of ≥0.2 ng/ml. ^3^ A subset of these patients progresses further to imaging-confirmed tumor recurrence (TR) and ultimately to castration-resistant prostate cancer (CRPC), an advanced and often lethal stage.^4^ What determines whether BR remains indolent or advances to TR and why patients with apparently similar genomic profiles follow divergent trajectories cannot be predicted from PSA kinetics alone. This clinical unpredictability signals an underlying biological complexity that neither genetics nor clinical markers individually resolve.

Classical clinicopathological variables used for stratification of prostate cancer patients are PSA, Gleason Score, and T Stage. These three inputs form the basis of the NCCN risk classification system, which has organized patients into very-low, low, intermediate, high, and very-high risk categories since its introduction in the 1990s and has been validated extensively in large cohorts. ^5^ The Gleason grading system, updated to the five-tier Grade Group system by the International Society of Urological Pathology (ISUP), quantifies architectural disorder in biopsy tissue and remains one of the most utilized histopathological predictors of outcome. ^6^ Nomogram-based tools represent a meaningful step forward from simple categorical risk grouping by combining multiple clinical variables, including PSA level, Gleason grade, T stage, and biopsy core involvement, into continuous probability estimates that are more individually tailored to each patient. ^7,8^ The further integration of multiparametric MRI into these nomogram frameworks has consistently improved their ability to predict adverse pathological findings at surgery and BR after treatment, demonstrating that imaging-derived spatial information about the tumor adds prognostic value beyond what clinical and biopsy variables alone can provide. ^9^ Despite these advances, all nomogram and imaging-based approaches share the same fundamental limitation: they assess risk at a single point in time and cannot capture how the tumor will evolve under the selective pressure of therapy.

Prostate cancer is a very heterogeneous disease. Prostate tumors are commonly multifocal, where each tumor lesion may possess different genetic signatures; resulting in different growth propensities and survival capabilities. Some common drivers for this genetic difference, extensively studied, are AR and PTEN. Androgen signaling is central to prostate cancer biology at every disease stage. The androgen receptor (AR) pathway governs prostate epithelial proliferation and survival, and approximately 20% of CRPC tumors harbor AR gene alterations in large-scale genomic datasets.^10^ Androgen deprivation therapy (ADT) exploits this dependency: by suppressing androgen-driven proliferation and inducing apoptosis in androgen-dependent cells, ADT initially reduces tumor burden but is rarely curative.^11^ Over time, many tumors develop resistance and progress to CRPC. AR can escape androgen deprivation through several mechanisms including gene amplification, activating ligand-binding domain mutations, and the expression of constitutively active splice variants such as AR-V7, which lacks the ligand-binding domain entirely and therefore remains transcriptionally active regardless of castrate androgen levels, predicting resistance to both enzalutamide and abiraterone in men with metastatic CRPC. ^12^ Another primary molecular driver of this resistance is loss of PTEN, whose deletion constitutively activates the PI3K/AKT pathway and sustains proliferation independent of androgen signaling.^13^ PTEN loss is one of the most frequently occurring genomic alterations in prostate cancer, present in approximately 20-40% of primary tumors and rising to higher frequencies in metastatic and castration-resistant disease.^14^ The relationship between PTEN loss and androgen receptor signaling is not simply additive but reciprocally interlinked: active AR signaling suppresses PI3K pathway activity, meaning that when PTEN is lost this brake is removed and PI3K/AKT signaling becomes self-sustaining even as androgen deprivation therapy suppresses AR, while conversely AR inhibition in turn reactivates AKT signaling, such that inhibiting one pathway rescues the other. ^15^ Downstream of AKT, MDM2 phosphorylation promotes p53 degradation, blunting the apoptotic response to therapeutic stress and compounding the survival advantage of PTEN-deleted cells. ^16^ Critically, PTEN loss is not universal. It is present in a subset of recurrent tumors and absent in others, making it a central candidate for explaining the divergent trajectories observed between BR and TR cohorts.

PTEN and AR signaling alterations are well-established drivers of prostate cancer aggressiveness. However, their phenotypic impact is shaped by the broader genetic background in which they operate, meaning that identical PTEN or AR alterations can produce divergent outcomes across tumors.

Beyond cell-intrinsic factors, spatial heterogeneity in androgen availability within the tumor microenvironment represents a key extrinsic modulator of recurrence dynamics. This androgen landscape is actively shaped by the collective uptake behavior of tumor cells, which can upregulate transporters from the SLCO family ^17^ to increase their rate of androgen acquisition. When cells sequester androgens at high rates, they deplete local concentrations and generate steep intratumoral androgen gradients, creating spatially unequal resource environments across the tumor. Separately, cancer cells can sustain intratumoral androgen levels through de novo steroidogenesis, upregulating enzymes such as CYP17A1, AKR1C3, and SRD5A1 to synthesize testosterone and dihydrotestosterone from cholesterol or circulating adrenal precursors such as DHEA, maintaining AR activity even at castrate systemic levels. ^18,19^ Together, these mechanisms mean that the androgen microenvironment experienced by any given cell depends not only on systemic hormone levels but on the local architecture of competing and synthesizing neighbors.

Epithelial-mesenchymal transition (EMT) is a biologically conserved program in which epithelial cells lose their polarity and cell-cell adhesion, downregulate E-cadherin, upregulate mesenchymal markers such as N-cadherin and vimentin, and acquire migratory and invasive capacity. This process is coordinated by master transcription factors including SNAI1, ZEB1/2, and TWIST, and is activated by signals including TGF-β, WNT, and NOTCH. ^20^ Critically, EMT is not a binary switch but a spectrum of intermediate hybrid states in which cells retain partial epithelial and mesenchymal properties simultaneously. The reverse process, mesenchymal-to-epithelial transition (MET), is equally important for metastatic colonization at distant sites. Cells that have undergone EMT to enter circulation must revert through MET to establish and proliferate within a secondary site, making cellular plasticity across this axis a defining feature of the full metastatic cascade.^21,22^ Variation in EMT state across tumor subpopulations directly shapes the spatial architecture of the tumor, as cells with differing adhesion and motility properties organize into distinct tissue-scale morphologies that determine the physical context in which clonal selection operates.

Thus, cancer progression is inherently a multiscale process, spanning molecular signaling, cellular decision-making, and tissue-level dynamics. Thus, cancer progression is inherently a multiscale process, spanning molecular signaling, cellular decision-making, and tissue-level dynamics. Critically, intrinsic and extrinsic sources of heterogeneity do not act independently; the molecular state of a cell shapes how it remodels its local microenvironment, and that remodeled environment in turn selects for or against particular cell states.

Mechanistic computational models ^23^ are therefore essential tools for disentangling these coupled influences, as they can explicitly represent the bidirectional interactions between cell-intrinsic signaling and extrinsic microenvironmental context across scales. A fundamental challenge is understanding how genetic alterations such as PTEN loss, which act at the molecular scale, propagate upward through the cellular and tissue scales to produce clinically observable recurrence trajectories. Prior computational approaches have addressed parts of this problem in isolation. ODE-based signaling models have elucidated intracellular pathway dynamics ^24,25^, while agent-based and spatial models have captured tissue-level clonal competition and tumor morphology. ^26,27^ However, these frameworks have largely been applied independently: signaling models lack spatial and metabolic context, and spatial models rarely incorporate mechanistic, patient-specific intracellular signaling. As a result, neither approach alone can account for how the same molecular alteration (PTEN loss) produces such different clinical outcomes across patients, nor can they explain why microenvironmental interventions such as androgen withdrawal produce cohort-specific rather than universal tumor responses. The core explanatory gap is the absence of a mechanistic coupling between cell-intrinsic genetic heterogeneity and extrinsic microenvironmental heterogeneity - a gap that, when left unfilled, renders both molecular and spatial analyses incomplete.

To address this gap, we developed an integrated multiscale framework that mechanistically couples cell-intrinsic signaling with extrinsic microenvironmental context and applied it to publicly available prostate cancer cohorts from The Cancer Genome Atlas (TCGA). Specifically, we focused on biochemical recurrence (BR) and treatment-resistant (TR) patient cohorts, selected because of the availability of gene expression data that enabled cohort-specific parameterization of intracellular signaling models. On the intrinsic side, the framework incorporates PTEN status and AR signaling as determinants of androgen sensitivity and apoptotic resistance. The multiscale hybrid systems (MHS) cellular model integrates the four signaling axes most central to prostate cancer fate decisions: AR signaling, PI3K/AKT/PTEN, Ras–MAPK, and p53-mediated DNA damage responses. The MHS model computes probabilistic cell-fate outcomes (growth, quiescence, or apoptosis) and is parameterized using cohort-specific differentially expressed genes from TCGA bridging static genomic profiles to dynamic cellular behaviors. To extend the framework to capture effect of extrinsic heterogeneity, we then embed this cellular model within a spatial ABM that simulates tumor growth in a mechanistically explicit tumor microenvironment (TME). In this TME, testosterone gradients shaped by cellular SLCO-mediated uptake kinetics, physical packing density, and adhesion-motility coupling modulate clonal competition and tumor spatial architecture. The MHS-defined cellular logic, encoding cohort-specific AR, mitogenic, and DNA damage responses, is directly wired to the ABM: intrinsic signaling state sets the proliferation, migration, and adhesion propensities of each agent, while local microenvironmental signals modulate those propensities in real time. This bidirectional architecture is the structural implementation of the coupling hypothesis. It is this coupling, not either factor in isolation, that constitutes the mechanistic basis of CRPC emergence and the source of the clinical unpredictability that motivates this work.

Using this framework, we demonstrate the coupling principle at each scale and across patient cohorts. At the cellular scale, PTEN loss drives earlier and more aggressive PSA-defined recurrence in a dose-dependent manner, but the TR cohort’s broader genomic background amplifies this effect in ways that PTEN status alone cannot predict, revealing a second tier of cell-intrinsic heterogeneity operating at the genome-wide level. Machine learning surrogates of the MHS model, analyzed by SHAP-based feature attribution, converge on PTEN, MDM4, and AR as the dominant intrinsic determinants of cell fate across three independent classifier architectures. Kaplan-Meier analysis of ∼4700 patients from cBioPortal validates that alterations in these genes confer significantly worse overall survival. At the tissue scale, ABM simulations reveal three distinct instances of intrinsic-extrinsic coupling with distinct mechanistic origins: physical confinement suppresses the proliferative advantage of PTEN-deleted clones regardless of genotype; androgen uptake kinetics create cohort-specific competitive landscapes that selectively enrich resistant clones in BR but paradoxically preserve heterogeneity in TR tumors due to the intrinsically lower androgen-dependence threshold of TR-sensitive cells; and PTEN-loss-driven suppression of E-cadherin sets the adhesive state that governs whether resistant cells undergo a threshold phase transition into spatially clustered aggregates. Clinical survival analysis corroborates both microenvironmental axes: high AKR1C3 expression independently predicts worse progression-free survival across primary and metastatic cohorts, high RHOA expression stratifies progression-free survival in primary localized disease, and loss of CDH1 expression associates with worse outcomes in a direction consistent with EMT-driven progression.

Finally, parameterizing the MHS model with individual gene expression profiles from 14 EUREKA1 trial patients ^28^ reveals patient-specific androgen sensitivity ratios that characterize individual ADT response potential, extending the framework to single-patient parameterization. Together, these findings demonstrate that mechanistic coupling of intrinsic molecular heterogeneity and extrinsic microenvironmental constraints is necessary to capture the diversity of CRPC progression trajectories observed clinically.

## Results

### Cell-Intrinsic Heterogeneity Contributes to Cohort-Specific Amplification of PTEN-Driven CRPC Progression

A central clinical challenge in prostate cancer management is understanding why patients with similar PTEN status exhibit markedly different disease trajectories. To evaluate how patient-specific genetic backgrounds modulate the downstream consequences of PTEN loss, we conducted the sensitivity analysis across three simulated patient cohorts: Control (CNT), Biochemical Recurrence (BR), and Tumor Recurrence (TR). Clinical and genomic data from the TCGA portal was used to stratify patients into these cohorts.

To delineate the specific contribution of cell-intrinsic signaling to disease progression, we utilized the PCa MHS cellular model to perform a targeted sensitivity analysis on PTEN. As a critical tumor suppressor, PTEN loss is mechanistically linked to constitutive activation of the PI3K/AKT pathway, a known driver of androgen independence. In addition, the cellular model was also parameterized by mapping cohort-specific differentially expressed genes (DEGs) to corresponding model species concentrations. (Described in Methods)

### Effect of PTEN Deletion Level on PSA Dynamics and CRPC Progression

We systematically varied the extent of PTEN deletion to model a spectrum of cellular resistance, while maintaining a constant growth factor environment (10 nM GF). We tracked Net Cell Growth (NCG) and total serum PSA concentrations under simulated ADT, monitoring for the onset of biochemical recurrence (BR), defined as the time point at which serum PSA exceeds the clinical threshold of 0.2 ng/mL.

Higher PTEN deletion levels consistently produced earlier recurrence and a more aggressive rise in PSA concentrations, resulting in substantially higher final PSA levels [Fig. 1]. These results demonstrate that even within a defined risk group, cell-intrinsic heterogeneity in PI3K/AKT pathway activity, conferred by variable degrees of PTEN loss, is sufficient to drive substantial variance in time to recurrence and rate of clinical progression. PTEN status thus emerges as a primary molecular determinant of intrinsic recurrence potential in this model.

**Figure 1.**
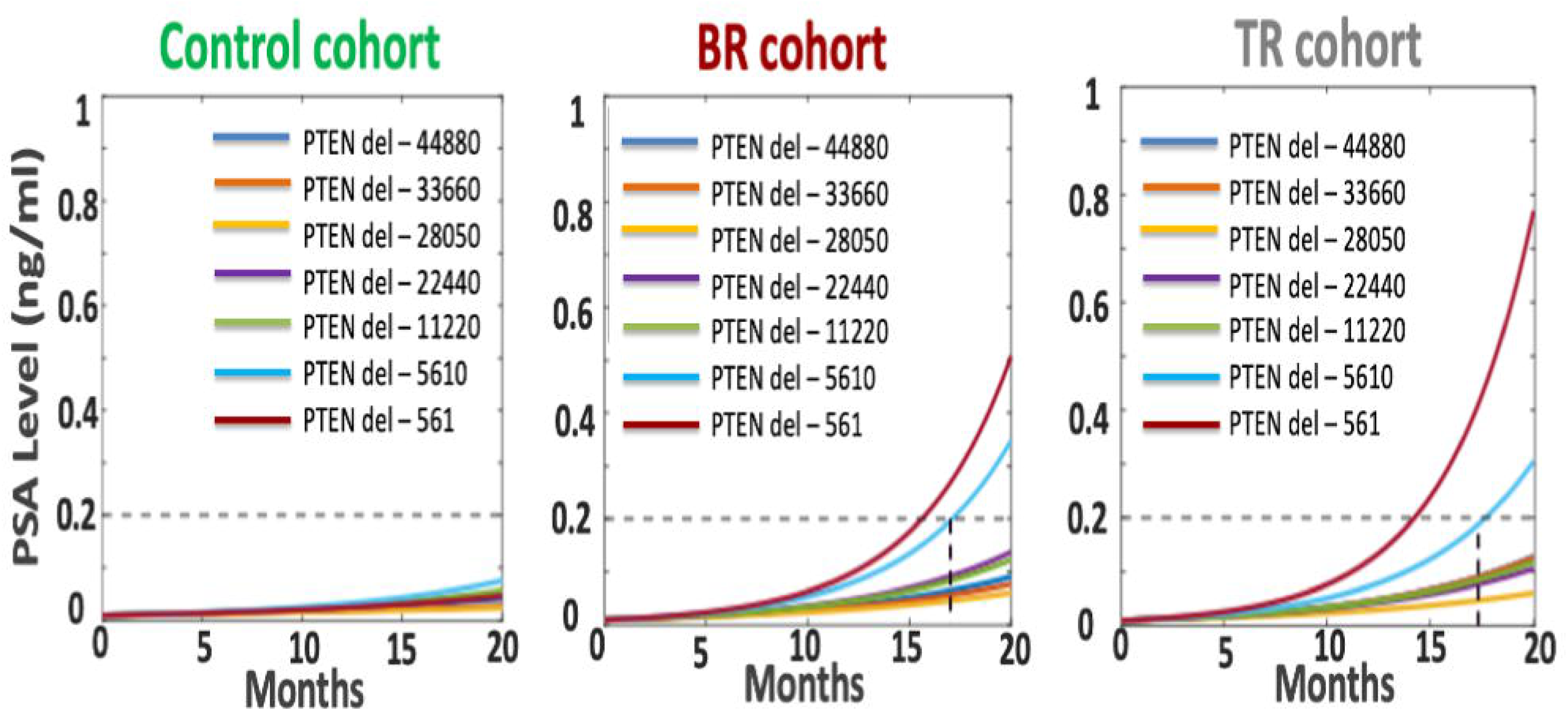
Cell-intrinsic genomic background amplifies PTEN-driven castration-resistant progression in a cohort-specific manner. Simulated serum PSA dynamics under ADT for the Control (CNT), biochemical recurrence (BR), and tumor recurrence (TR) cohorts across seven PTEN deletion levels (increasing severity, blue to dark red). Each cohort was parameterized from TCGA-derived differentially expressed genes mapped to the cellular MHS model; basal GF was fixed at 10 nM. The dashed line at 0.2 ng/mL marks the clinical BCR threshold. CNT remains sub-threshold regardless of PTEN status; BR and TR both show dose-dependent PSA escalation, but at equivalent deletion levels TR rises earlier and more steeply, indicating that PTEN loss provides the proliferative signal for recurrence while the broader TR genomic background amplifies its severity.

With basal GF fixed at 10 nM across all cohorts and PTEN deletion level varied systematically, we simulated NCG and total PSA concentration for each group. The CNT cohort remained stable regardless of PTEN status, confirming that the recurrence-permissive genomic background is a prerequisite for PTEN-driven progression. Both BR and TR cohorts exhibited PSA-defined recurrence, but the intensity of this response diverged sharply between the groups. At the highest PTEN deletion levels, the TR cohort displayed a significantly more aggressive phenotype than the BR cohort, characterized by a steeper and more rapid escalation in serum PSA [Fig. 1]. This result reveals a critical finding: while PTEN loss provides the necessary proliferative signal for recurrence, the broader genomic context of the TR cohort acts as a potent amplifier, translating the same molecular perturbation into a markedly more severe clinical outcome. Cell-intrinsic heterogeneity, therefore, operates at two levels: within individual pathway components (PTEN dosage) and across the genome-wide signaling background of distinct patient cohorts.

### Survival Analysis Validates ML-Identified Molecular Drivers Machine Learning Surrogate and SHAP-Based Feature Ranking

The computational cost of conducting global sensitivity analysis directly on the MHS model, which integrates ODEs and Boolean logic across disparate time scales, prohibits the use of traditional variance-based methods such as Latin Hypercube Sampling combined with Sobol indices. To overcome this bottleneck, we trained a machine learning (ML) surrogate to approximate the full MHS model, enabling computationally efficient exploration of the high-dimensional parameter space. ^29^

MHS model simulations across three cohorts were used to generate training data, with net cell growth (NCG) probability as output. To determine the optimal threshold for binary classification of cell fate as proliferative (NCG ≥ 0.15) or quiescent (NCG < 0.15), we trained supervised ML classifiers across thresholds ranging from 0 to 0.4 in increments of 0.05 and selected the threshold yielding the highest balanced accuracy on the test set. [Fig S5.2.1] Three classifier architectures - Neural Network (NN), Random Forest (RF), and Support Vector Machine (SVM) were trained on the resulting binary outputs, with input features z-score normalized, class imbalance corrected using SMOTE, and hyperparameters optimized via 5-fold cross-validation.

Shapley Additive Explanations (SHAP) were then applied to each trained surrogate to rank molecular species by their marginal contribution to cell fate predictions. The NN model ranked MDM4, PTEN, AR, and CASP9 as its top features; the RF model ranked PTEN, AR, MDM4, MDM2, phosphorylated AKT, and phosphorylated ERK; and the SVM model ranked PTEN, MDM4, AR, BCL2, RAF, and MDM2. Critically, PTEN, MDM4, and AR appeared as top features across all three model architectures [Fig. 2], providing a consensus identification of the dominant intrinsic determinants of cell fate that is robust to classifier choice.

**Figure 2.**
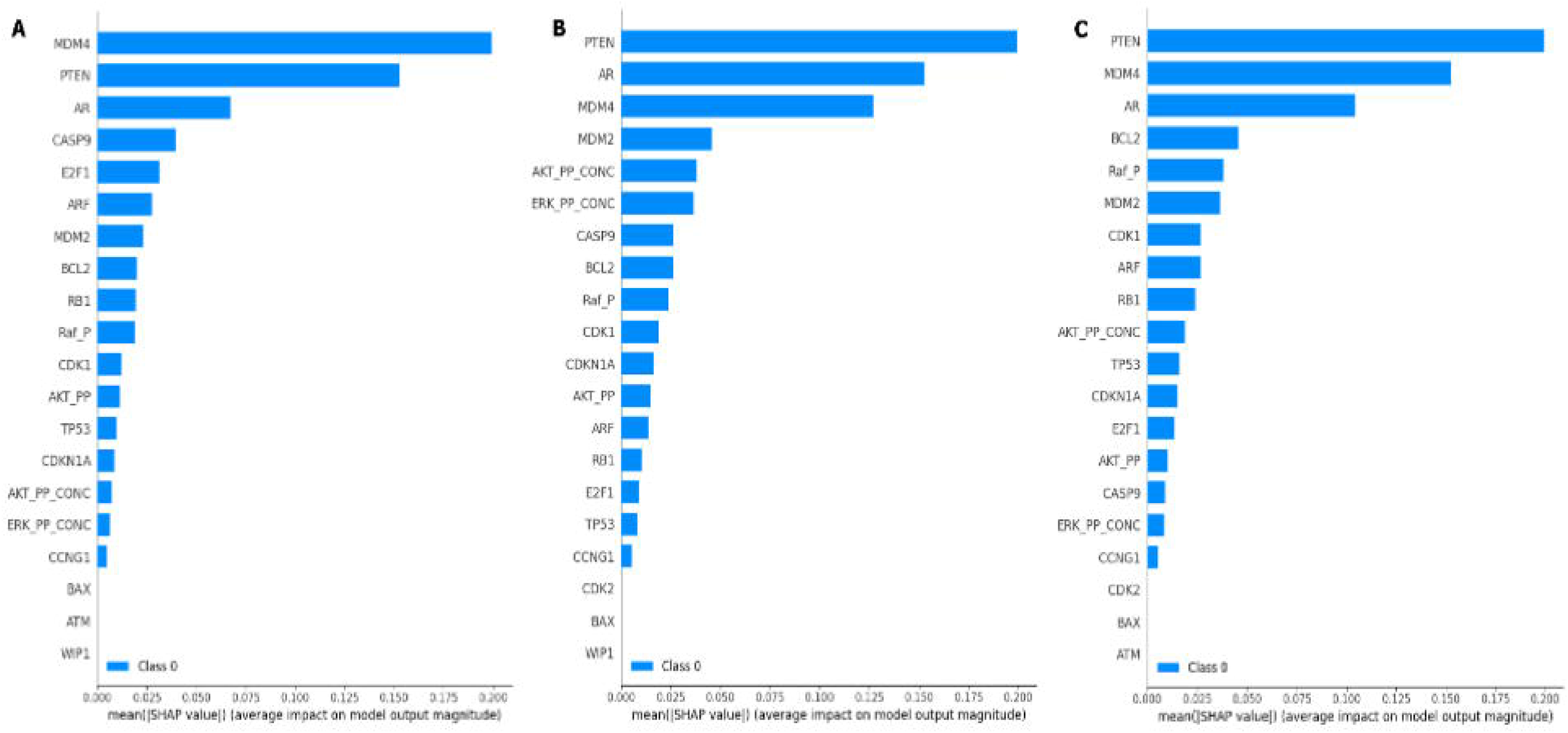
SHAP-based feature ranking across three ML surrogate models. A supervised ML surrogate was trained to approximate the cellular MHS model, enabling efficient global sensitivity analysis. SHAP summary plots rank input features by mean marginal contribution to cell-fate prediction for each classifier: (A) neural network, (B) random forest, (C) support vector machine. PTEN, MDM4, and AR emerge as consensus drivers across all three architectures, giving a classifier-independent identification of the dominant determinants of castration-resistant cell fate.

To evaluate the clinical relevance of the intrinsic drivers identified by SHAP analysis, we performed Kaplan-Meier (KM) survival analysis on prostate cancer patient data from cBioPortal. Survival curves were generated for patients stratified by the presence or absence of alterations in the top ML-ranked genes: PTEN, AR, and MDM4. For each gene, we compared overall survival probability between patients with and without alterations, with statistical significance assessed by log-rank test (significance threshold p < 0.05). Multivariate hazard ratios (HR) were also calculated with cohort and patient age as the covariates. Among the consensus top-ranked genes, AR alterations conferred the most severe survival disadvantage, with a median OS of 20 versus 86 months in wildtype patients (log-rank p < 0.001; multivariate HR = 4.07 [95% CI: 3.62–4.58]). PTEN loss independently predicted poor outcomes (median OS 54 vs. 77months; p < 0.001; HR = 1.51 [1.36–1.69]), consistent with its role as the primary PI3K/AKT dysregulation node identified by SHAP. MDM4 amplification was associated with significantly shortened survival (median OS 33 vs. 75 months; p < 0.001; HR = 2.25 [1.66–3.05]). (Fig 3A-C) When patients harboring any alteration in PTEN, AR, or MDM4 were combined into a single OR-classifier, the survival separation remained highly significant, spanning 29% of the cohort (median OS 41 vs. 95 months; p < 0.001; HR = 2.38 [2.16–2.63]. (Fig 3D) This direct correspondence between model-ranked features and clinical survival outcomes provides validation that the MHS model is capturing biologically and clinically meaningful pathway dynamics.

**Figure 3.**
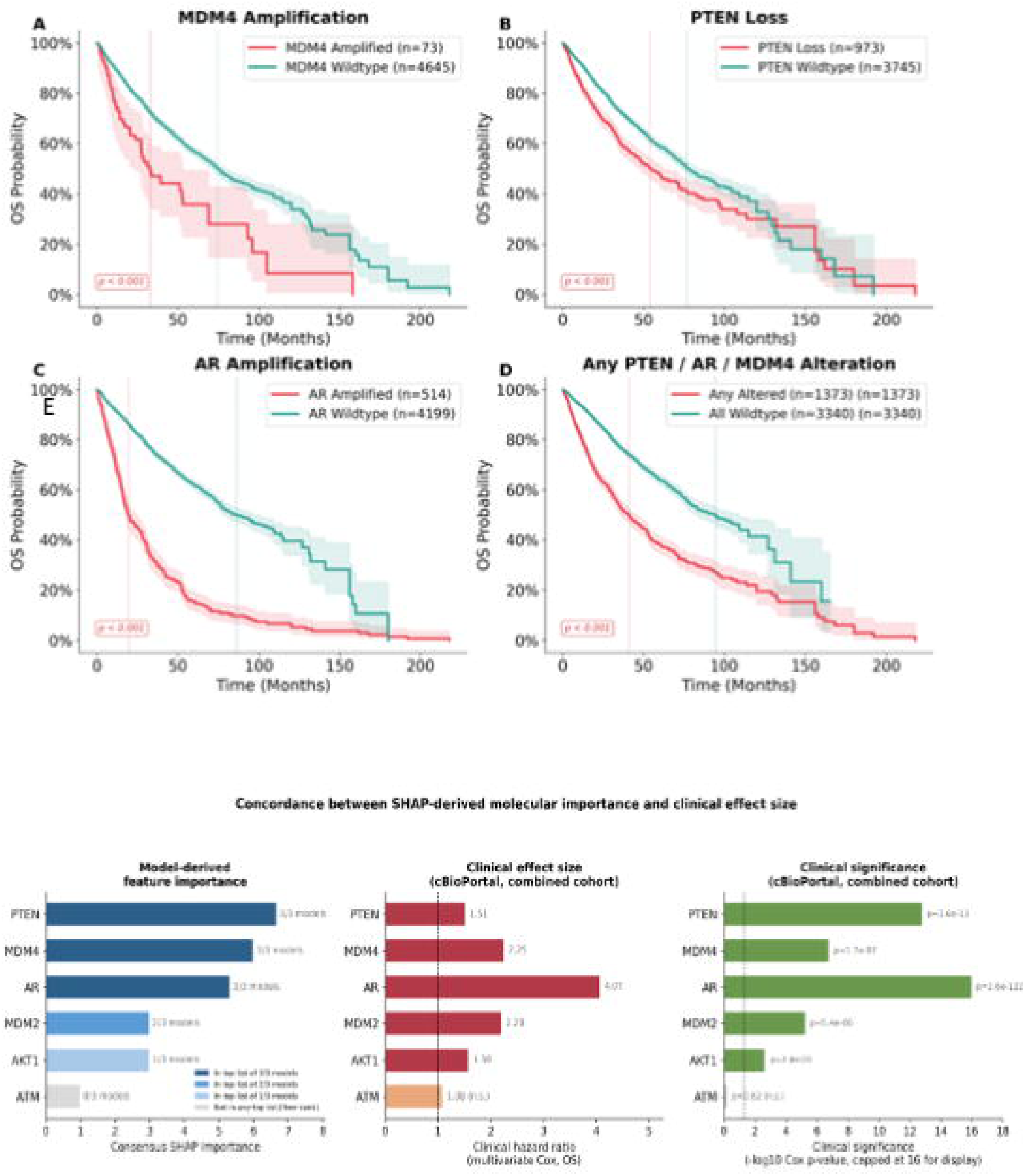
Kaplan-Meier survival analysis validates SHAP-identified molecular drivers of castration-resistant progression. Overall survival probability was plotted for patient cohorts with (red) and without (blue) genomic alterations in SHAP-consensus features identified by the MHS surrogate model, across seven cBioPortal prostate cancer cohorts (N= 4718 patients). (A) MDM4 amplification. (B) PTEN loss. (C) AR mutation/amplification. (D) Combined alteration burden: any alteration in PTEN, AR, or MDM4 (==27%== of cohort). All p < 0.001. (E) Comparing the consensus SHAP rank across the three classifier architectures against two independent measures of clinical importance derived from the combined cBioPortal cohort: hazard ratio magnitude and statistical significance. Consensus SHAP importance (x-axis, left panel) is calculated as the maximum mean rank across models plus one, minus each gene’s own mean rank position across the surrogate models in which it appeared, so that longer bars indicate higher consensus importance; this value reflects relative rank order rather than continuous SHAP magnitude.

KM analyses were additionally performed for other MHS model species including MDM2, AKT1, ATM, CDKN1A, BAX, CASP9, BCL2, and RAF1. MDM2 and AKT1 alteration reached statistical significance (MDM2: median OS 30 vs. 74 months, HR = 2.20 [1.57–3.09]; AKT1: median OS 52 vs. 75 months, HR = 1.58 [1.18–2.11]). ATM, CDKN1A, CASP9, BAX, BCL2, and RAF1 did not meet the significance threshold, primarily due to low genomic alteration frequencies across the available cohorts (altered arm n < 20 in most datasets). This analysis is elaborated in Supplementary S6.1.

To assess whether the relative ranking of SHAP-identified molecular drivers corresponds to relative clinical importance, we compared the consensus SHAP rank across the three classifier architectures against two independent measures of clinical importance derived from the combined cBioPortal cohort: hazard ratio magnitude and statistical significance. (Fig 3E) Genes with fewer than 20 altered patients in the combined cohort (CDKN1A, RAF1, BCL2, CASP9, BAX) could not be assigned a hazard ratio and were excluded from this comparison rather than treated as unimportant, since rarity of alteration does not imply lack of biological relevance. Among the six remaining genes, concordance with hazard ratio magnitude was directionally positive but modest while concordance with statistical significance was stronger. The discrepancy was driven largely by PTEN, whose high alteration frequency in this cohort (N 973) yields a very small p-value (1.58e-13) but a comparatively modest hazard ratio (1.51) relative to the rarer AR and MDM4 amplifications. Despite this difference, both measures of clinical importance converged on the same three consensus genes identified by SHAP across all three model architectures, PTEN, AR, and MDM4, as the top-ranked features by either criterion. This convergence indicates that the model’s intrinsic importance ranking corresponds to clinically meaningful molecular drivers, while the differing strength of concordance across metrics reflects the distinct biological roles of common, moderate-effect alterations such as PTEN loss versus rarer, higher-penetrance events such as AR and MDM4 amplification.

The KM results thus bridge computational feature importance and patient prognosis, establishing PTEN, AR, and MDM4 as the clinically validated molecular determinants of risk most consistently identified by the MHS surrogate model. We note that while the SHAP analysis describes the behavior of the surrogate model rather than establishing causal biological mechanisms, the convergence of model predictions and survival data impresses upon the biological importance of these pathway nodes as drivers of disease progression.

### Tissue-Scale Modeling Reveals Microenvironmental Constraints as Critical Regulators of Cohort-Specific Clonal Selection and Tumor Progression

While the MHS model successfully elucidated the role of cell-intrinsic genetic heterogeneity, specifically the impact of PTEN loss and cohort-specific genomic backgrounds on recurrence potential, tumor progression is rarely governed by intracellular signaling alone. The microenvironment exerts continuous physical and metabolic pressures on cancer cells, and the central question motivating this work is how intrinsic clonal fitness and extrinsic microenvironmental constraints are coupled to produce the emergent tissue-level behaviors observed across patient cohorts. The multiscale framework is described in detail under Supplementary Section 7.

To systematically investigate how tissue-level outcomes emerge from the coupling of cell-intrinsic states and extrinsic environmental pressures, we simulated the joint dynamics of PTEN-resistant (R) and PTEN-sensitive (S) clones across a range of microenvironmental conditions. Importantly, these simulations do not model *de novo* genetic mutations; the R and S phenotypes possess fixed, inherent growth potentials defined by the MHS systems biology model. The observed changes in tumor composition and morphology are therefore the result of differential clonal fitness - the effectiveness with which each fixed phenotype proliferates or survives when subjected to local variations in testosterone concentration, spatial packing, and cell-cell adhesion within the tumor microenvironment.

### Crowding and Physical Confinement Modulates the Proliferative Advantage of PTEN-Deleted Clones

PTEN-deleted prostate cancer cells possess an intrinsic resistance to ADT-induced apoptosis. If such cells persist following prostatectomy, their genotype can drive recurrence. However, whether this intrinsic advantage translates into accelerated tumor growth depends critically on the physical context of the residual tissue. In solid cancers, dense cellular packing activates contact inhibition, a critical brake on tumor growth. As tumors become crowded, cells sense compression through mechanosensitive ion channels at the membrane, amplifying signals via PIP2 and phosphoinositide lipids. The Hippo pathway then relays these signals to sequester YAP and TAZ in the cytoplasm, preventing growth gene expression. This converges on the cell cycle, where crowding raises p21 and p27 inhibitors while lowering Cyclin D1, halting division. Dense packing also creates hypoxia that reinforces arrest, and a coordinated cytoskeletal program allows cells to withstand mechanical load. This system means tissue architecture, not genotype alone, sets the ceiling on cancer aggressiveness.

We modeled two distinct spatial scenarios: a ‘Low Crowding Density’ (small confinement) setting, representing a spatially unconstrained environment where cells have ample space to proliferate and migrate, and a ‘High Crowding Density’ (dense confinement) setting, representing a spatially constrained environment where cells are immediately subject to significant contact inhibition and crowding. While the intrinsic MHS cellular model predicts that higher initial R/S ratios should yield higher net growth, ABM simulations reveal that this potential is modulated by physical confinement. In the loose packing scenario, the environment acted as a permissive substrate, significantly amplifying the aggressive growth trajectory of high-R/S tumors. Conversely, dense tissue packing imposed contact inhibition that substantially dampened this intrinsic advantage [Fig. 4]. This behavior was consistent across both BR and TR cohorts, demonstrating that the aggressiveness of a tumor is not solely an intrinsic property but an emergent one that depends on the physical permissiveness of the surrounding tissue architecture. Intrinsic genotype sets the ceiling of aggressive potential; the microenvironment determines how much of that potential is realized.

**Figure 4:**
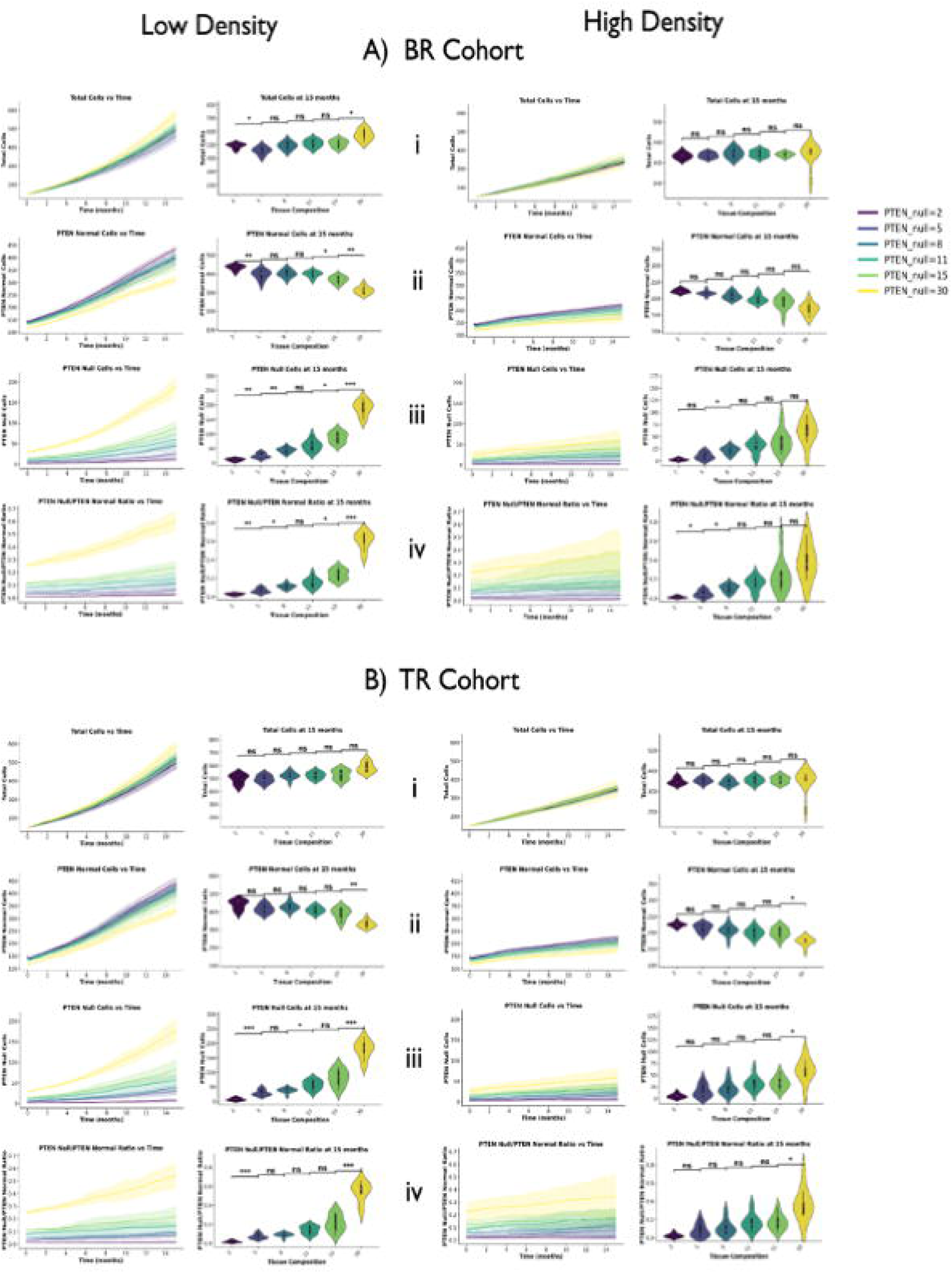
Physical confinement gates the proliferative advantage of PTEN-deleted clones in BR and TR cohorts. ABM simulations of prostate cancer tumor tissue under two spatial regimes: low packing density (loose confinement), representing a permissive environment where cells have ample space to proliferate and migrate, and high packing density (dense confinement), representing a physically constrained environment where contact inhibition is immediately enforced. Results shown for BR (A) and TR (B) cohorts; Each condition was simulated with 10 stochastic replicates; differences in means and variance between conditions were assessed using the Mann-Whitney U test and Levene’s test, respectively, with Bonferroni correction for multiple comparisons at a significance threshold of p = 0.05. Significance levels are denoted as * p < 0.05, ** p < 0.01, and *** p < 0.001. Color gradient: dark purple (lowest PTEN deletion) → yellow (highest). Shaded bands = ±1 SD. The x-axis represents initial tumor genetic composition as the ratio of PTEN-deleted (Resistant) to PTEN-normal (Sensitive) cells. The y-axis captures temporal profiles of total tumor load (i), PTEN-normal cell count (ii), PTEN-deleted cell count (iii), and R/S ratio (iv), and distributions of these same quantities at 15 months are shown as violin plots. Under loose confinement, spatial permissiveness amplifies the intrinsic growth advantage of high R/S tumors. In contrast, dense confinement substantially dampens this potential through contact inhibition, revealing that realized tumor aggressiveness is an emergent property of the coupling between intrinsic genotype and physical tissue architecture.

### Testosterone Uptake Kinetics Drive Cohort-Specific Clonal Selection

Under ADT, prostate tumors must survive in a systemic environment of profound androgen scarcity. One key mechanism of resistance is the upregulation of SLCO membrane transporters (SLCO2B1, SLCO1B3), which facilitate active uptake of testosterone and androgen precursors including DHEAS, maintaining intracellular androgen levels that sustain proliferation despite low systemic concentrations. ^30^ In CRPC, SLCO transporters are known to be further upregulated, supporting continued cancer growth even when systemic androgens are severely depleted. ^17^ By focusing on this uptake mechanism, we can investigate how individual-cell resource sequestration alters the shared androgen landscape of the tumor, and how that altered landscape feeds back onto the proliferative dynamics of competing clones - the core coupling question at the tissue scale.

A cellular uptake rate parameter governs substrate depletion, modulating the diffusion–reaction of testosterone. This parameter serves as a proxy for SLCO transporter expression levels. Starting from a mixed tumor with an R/S ratio of 1, we conducted sensitivity analysis on the cellular androgen uptake rate. Our simulations revealed that high uptake rates create localized zones of testosterone depletion, reducing interstitial androgen concentration available to neighboring cells. However, the proliferative consequences of this self-induced depletion depend on each cohort’s intrinsic androgen sensitivity, a property defined by the MHS model and parameterized from TCGA data, producing strikingly divergent outcomes between cohorts [Fig. 5A].

**Figure 5:**
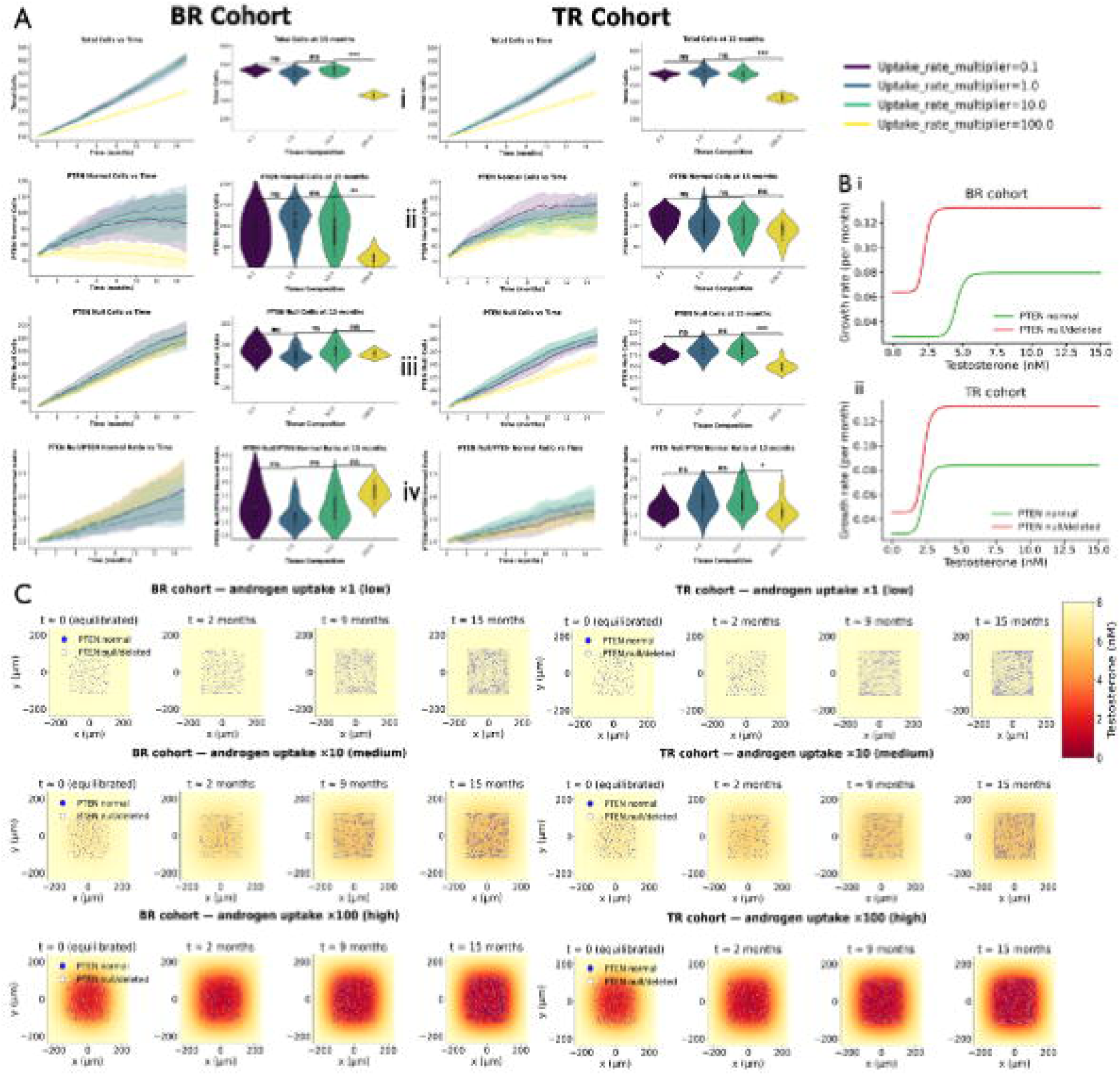
Androgen uptake rate governs cohort-specific clonal selection through resource-mediated competitive displacement. ABM sensitivity analysis on cellular testosterone uptake rate, initialized from a mixed tumor (R/S ratio = 1) for BR and TR cohorts. The x-axis represents uptake rate magnitude relative to nominal (X0) across all panels. (A) Temporal profiles of total tumor load (i), PTEN-normal cell count (ii), PTEN-deleted cell count (iii), and R/S ratio (iv), with violin plots showing distributions of each quantity at 15 months. (B) MHS model-derived proliferative response of PTEN-resistant (R) and PTEN-sensitive (S) cells as a function of local testosterone concentration for the BR cohort (i) and TR cohort (ii), defining the intrinsic androgen sensitivity that governs each cohort’s competitive response to uptake-driven depletion. (C) PhysiCell simulation snapshots of spatial testosterone distribution at representative time points under low (i) and high (ii) uptake conditions for BR and TR cohorts, illustrating the localized androgen depletion zones that emerge at high uptake rates and whose competitive consequences are determined by the cohort-specific sensitivity curves in (B). Each condition was simulated with 10 stochastic replicates; differences in means and variance between conditions were assessed using the Mann-Whitney U test and Levene’s test, respectively, with Bonferroni correction for multiple comparisons at a significance threshold of p = 0.05. Significance levels are denoted as * p < 0.05, ** p < 0.01, and *** p < 0.001.

In the BR cohort, sensitive cells (SBR) exhibited strong nonlinear dependence on androgen availability. As resource sequestration reduced local testosterone from physiological levels (∼10 ng/mL) to intermediate levels (∼2.5 ng/mL), the proliferative capacity of SBR cells declined sharply, falling well below that of resistant cells (RBR), which maintained robust proliferation at intermediate androgen concentrations. [Fig. 5B,C]. This differential response created a broad competitive window that enabled PTEN-resistant clones to outcompete their sensitive counterparts, progressively increasing the R/S ratio. Moderate-to-high uptake rates thus act as a selective pressure that enriches the resistant population in BR tumors, converting a heterogeneous mixed tumor into a predominantly resistant one.

In stark contrast, the TR cohort exhibited resistance to this selection pressure due to a significantly lower threshold of androgen dependency. Unlike the BR phenotype, TR cohort sensitive cells (STR) cells maintained robust proliferation well below the 2.5 ng/mL mark, with their proliferative threshold shifted to extremely low concentrations (<1 ng/mL). [Fig. 5B,C]. Consequently, even under conditions of high uptake-driven depletion, STR cells continued to proliferate alongside resistant clones. This physiological robustness prevented resistant clones from achieving competitive dominance, allowing the TR cohort to maintain a heterogeneous tumor composition even under significant resource stress. This result is a direct demonstration of intrinsic - extrinsic coupling: the same microenvironmental pressure (androgen gradient steepness) produces opposite clonal selection outcomes in BR and TR tumors because the intrinsic androgen sensitivity of sensitive cells, defined by cohort-specific genomic profiles, determines the competitive response function.

### Adhesion–Motility Coupling Drives Spatial Clustering as a Threshold Phase Transition

While resource gradients represent one axis of microenvironmental coupling, tumor spatial architecture is also governed by the physical mechanics of cell–cell interaction. The Epithelial–Mesenchymal Transition (EMT) is a critical program in which epithelial cells shed adhesive constraints, typically through downregulation of E-cadherin and N-cadherin, and acquire a motile, invasive mesenchymal phenotype. In prostate cancer, reduced E-cadherin expression is a hallmark of EMT and is associated with poor prognosis.^31^ Importantly, PI3K/AKT pathway activation via PTEN loss suppresses E-cadherin expression through transcriptional repressors such as Snail and through post-translational mechanisms ^32^, directly linking the intrinsic genetic alterations captured in the MHS model to the mechanical phenotype of the cell in the ABM. Conversely, the re-acquisition of adhesive traits is often necessary for the collective migration and colonization of circulating tumor cells at distant sites, such as the bone. Here, upregulation of E-Cadherin and EpCAM leads to enhanced cell–cell adhesion, interaction with osteoblasts, and better metastatic colony formation.^33^ This linkage is a second structural instance of intrinsic–extrinsic coupling: PTEN status sets the adhesive state of the cell, which then determines how that cell interacts with the spatial architecture of the tumor.

To dissect how variations in adhesive strength drive emergent spatial organization, we performed sensitivity analysis on cell–cell adhesion and motility parameters within the ABM, tuning the adhesion strength parameter. We assumed a reciprocal relationship between adhesion and motility as a proxy for the EMT spectrum: high-adhesion agents (epithelial-like) were assigned proportionally lower motility vectors, while low-adhesion agents (mesenchymal-like) were assigned high motility. Simulations were initialized with a randomly mixed R/S population (ratio = 1) under normal androgen conditions (8 ng/mL), with adhesion and motility of resistant agents varied by an order of magnitude (0.1×, 1×, 10× nominal) while sensitive cells were held at nominal values.

Our simulations revealed that variations in adhesion and motility had negligible impact on overall tumor load but drove significant divergences in spatial morphology and tissue composition (R/S ratio). [Figs. 6,7]. At low-to-nominal adhesion levels, motile forces dominated, producing a ‘fluid’ tissue architecture in which resistant cells dispersed freely among the sensitive population. Beyond a critical adhesion threshold, the system underwent a rapid phase transition: adhesive forces dominated, driving the self-organization of resistant cells into dense, segregated clusters that excluded sensitive neighbors. To quantify this morphological transition, we defined a normalized clustering index C(t) based on the fraction of homotypic neighbors for each agent/cell, normalized to its value at initialization, such that C(t) ≈ 1 indicates maintained spatial mixing and C(t) > 1 indicates progressive self-segregation of resistant clones relative to the initial random distribution [Fig. 7]. This sigmoidal relationship between adhesion strength and clustering index demonstrates that spatial reorganization in the tumor is a threshold-dependent emergent phenomenon - a morphology transition governed by the coupling of intrinsic adhesive state (set by PTEN-driven PI3K/AKT activity) and the extrinsic physical laws of contact mechanics.

**Figure 6:**
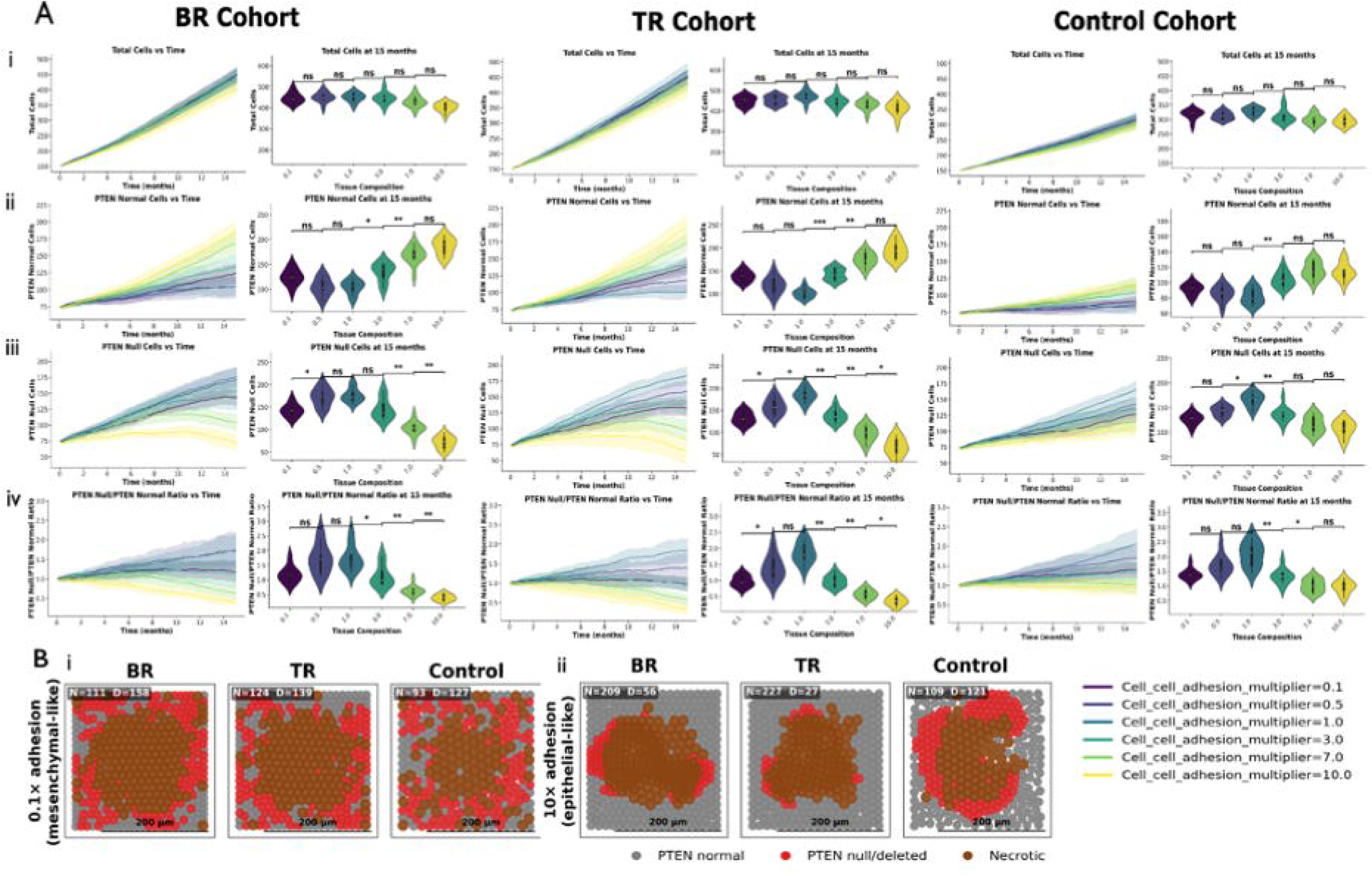
Adhesion and motility govern spatial morphology without altering overall tumor load. ABM sensitivity analysis on cell-cell adhesion and motility parameters of PTEN-resistant agents, initialized from a randomly mixed R/S population (ratio = 1) under physiological androgen conditions (8 ng/mL) and 1x androgen uptake multiplier. Adhesion and motility of resistant agents were varied reciprocally across an order of magnitude (0.1×, 1×, 10× nominal) to proxy the EMT spectrum, from mesenchymal-like (low adhesion, high motility) to epithelial-like (high adhesion, low motility), while sensitive agent parameters were held at nominal values. (A) Temporal profiles of total tumor load (i), PTEN-normal cell count (ii), PTEN-deleted cell count (iii), and R/S ratio (iv) and corresponding violin plots at t=15months shown across BR, TR, and Control cohorts; illustrating that despite order-of-magnitude variation in adhesion strength, net tumor composition remains statistically equivalent across all conditions. (B) PhysiCell simulation snapshots illustrating emergent spatial morphology under low (i) and high (ii) adhesion conditions for BR, TR, and Control cohorts. Each condition was simulated with 10 stochastic replicates; differences in means and variance between conditions were assessed using the Mann-Whitney U test and Levene’s test, respectively, with Bonferroni correction for multiple comparisons at a significance threshold of p = 0.05. Significance levels are denoted as * p < 0.05, ** p < 0.01, and *** p < 0.001.

**Figure 7:**
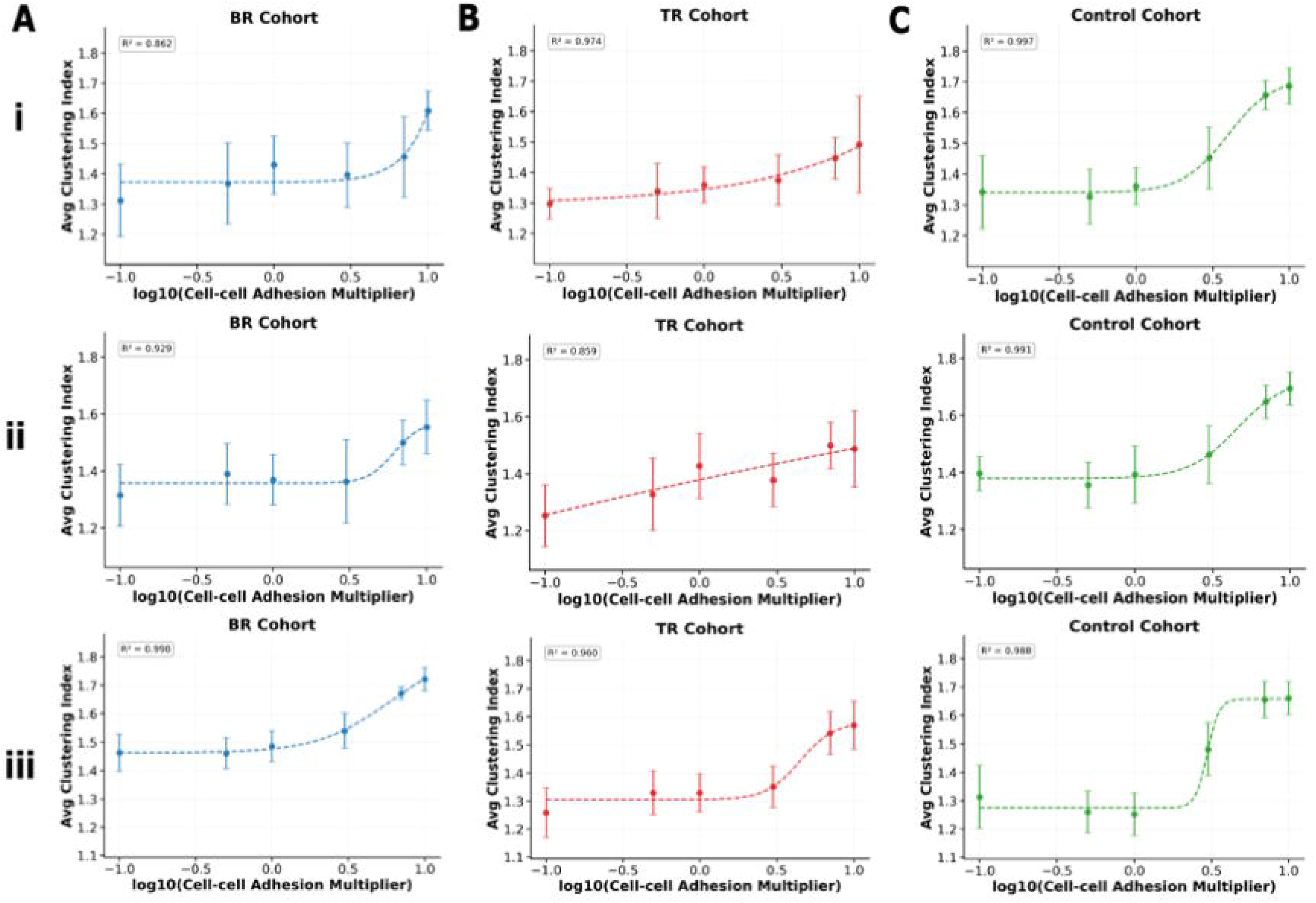
Adhesion strength drives a threshold phase transition in spatial segregation of PTEN-resistant clones across cohorts and androgen uptake conditions. Normalized clustering index C of PTEN-deleted cells at t = 15 months as a function of log-transformed cell-cell adhesion strength relative to nominal (log (X/X)), shown for BR (A), TR (B), and Control cohorts (C) (columns) across three androgen uptake rates (rows: ×0.1 (i), ×1 (ii), ×100 (iii)). C is defined as the fraction of a cell’s physical (contact-based) neighbors that share the same type, averaged across all PTEN-deleted cells. At t = 0, the population is initialized as a randomly mixed 1:1 R/S ratio, yielding a baseline of C ≈ 0.5, where resistant cells are equally likely to neighbor sensitive or resistant cells by chance. C → 1 reflects progressive homotypic segregation of resistant clones into spatially coherent clusters. A sigmoidal function was independently fit to each cohort–uptake condition (dashed curve); goodness of fit is reported as R². Data points represent means ± SD across 10 stochastic replicates per adhesion condition. The transition threshold and steepness vary across cohorts and androgen uptake conditions, indicating that the androgen microenvironment and treatment context modulate both the adhesion level required to trigger spatial segregation and the sharpness of that transition.

Extending this analysis across cohorts revealed several key insights. First, total tumor load remained statistically unchanged across the full range of adhesion strengths, confirming that adhesion-motility variation does not drive net tumor growth. However, the R/S ratio declined with increasing adhesion, with PTEN-deleted cells becoming progressively less dominant relative to sensitive cells at high adhesion levels. This indicates that while high adhesion drives spatial reorganization of resistant clones into dense homotypic clusters, it simultaneously constrains their competitive advantage by reducing motility-driven dispersal, shifting the balance toward sensitive cells without altering overall tumor burden. Second, the adhesion threshold required to trigger clustering and the sharpness of that transition both varied across cohorts. Control cells showed the lowest threshold and sharpest transition, meaning a relatively modest increase in adhesion was sufficient to drive strong spatial segregation. BR cells required higher adhesion levels to trigger clustering, and this threshold rose further under low androgen conditions. TR cells showed the most inconsistent behavior, with no detectable transition at nominal androgen levels and only a partial recovery at high androgen. Third, the maximum clustering magnitude reached at high adhesion followed the order Control > BR > TR. TR resistant cells not only transitioned later and less sharply but also clustered less strongly overall, pointing to a more motility-dominant phenotype even under high adhesion conditions.

Finally, the effect of androgen uptake on clustering was cohort specific. Control clustering sharpened considerably with increasing androgen availability, suggesting that androgen-rich conditions reinforce adhesion-driven spatial organization in untreated cells. BR clustering was largely insensitive to uptake rate in transition shape but reached its highest clustering magnitude at high uptake. TR clustering was minimally affected by uptake rate across all adhesion levels, consistent with a partial decoupling of AR signaling from adhesive mechanics in castration-resistant cells.

Taken together, the three microenvironmental axes (physical confinement, androgen resource gradients, and adhesion-motility) each operate through a distinct biophysical mechanism yet converge on a common principle: intrinsic genotype defines the boundaries of clonal potential, while the microenvironment determines which fraction of that potential is realized and in what spatial form. The cohort-specific divergences observed across all three axes are not coincidental; they are the direct consequence of the MHS-encoded genomic differences between BR, TR, and Control cohorts interacting with shared physical and metabolic constraints. This establishes microenvironmental gating as a generalizable organizing principle of cohort-level tumor heterogeneity in prostate cancer.

### Clinical Survival Analysis Corroborates Microenvironmental Drivers found to be sensitive in the Tissue-Scale ABM

To evaluate whether the microenvironmental axes hypothesized by the ABM are reflected in clinical outcomes, we performed KM survival analysis on prostate cancer patient data from cBioPortal across four mRNA-eligible cohorts, with expression-based stratification applied to genes mapping onto three functional axes: crowding/mechanotransduction, androgen uptake/synthesis, and adhesion/motility/cytoskeleton.

### Crowding/Mechanotransduction Axis

Our ABM shows that physical crowding suppresses the aggressiveness of PTEN-deleted cells: dense packing dampens their proliferative advantage (Fig. 4). We tested whether transcript levels of the genes that execute this brake track recurrence in TCGA-PRAD (n = 494), screening 27 genes across six contact-inhibition pathways on two endpoints (DFS, PFS), with each gene additionally tested for a gene-by-PTEN interaction. (Since PTEN loss supports aggressive growth, we also asked whether each gene’s prognostic effect differs between PTEN-deleted and PTEN-normal tumors.) Full results are in Supplementary Table S6.4.1.

We tested genes from six pathways that normally enforce crowding-induced growth arrest --Hippo/YAP-TAZ contact inhibition^34^, mechanosensitive ion channels^35^, phosphoinositide kinases^36^, cell-cycle inhibitors^37^, hypoxia mediators ^38^, and a PIP2-trafficking/cytoskeletal signature^39^.

Fourteen genes reached nominal significance (p < 0.05, log-rank on Kaplan Meier curves or Cox HR) on at least one endpoint: three in the Hippo pathway (YAP1, WWTR1, LATS1), one phosphoinositide kinase (PIK3CA), two cell-cycle arrest genes (CCND1, MKI67), two hypoxia mediators (HIF1A, VEGFA), and six PIP2-trafficking/cytoskeletal genes (RAE1, PLS1, EZR, LAMP2, HMOX1, FLNC) (full results in Supplementary Table S6.4.1). The crowding axis screen evaluated 27 genes across two recurrence endpoints and associations have been reported at nominal significance (p < 0.05). It is important to note that, all results are limited by the few recurrence events in this primary cohort (30 DFS, 93 PFS), and warrant confirmation in a higher-event metastatic cohort. Combined with the low recurrence event counts in this primary cohort, which limit statistical power, these associations should be read as hypothesis-generating rather than confirmatory: they nominate candidate contact-inhibition genes for validation in higher-event, expression-annotated cohorts.

The Hippo pathway is known to be the cell’s main system for sensing crowding and shutting down growth in response. ^40,41^ YAP1 (KM log rank p = 0.037), WWTR1, and LATS1 were each individually significant, with high expression of all three associated with better recurrence-free survival (YAP1 HR 0.62, p=0.011; WWTR1 HR 0.47, p=0.004; LATS1 HR 0.58, p=0.056, DFS; Table S6.4.1). For LATS1, the arrest-enforcing kinase of the pathway, this direction matches the screen’s expectation, since higher LATS1 should reinforce growth arrest and delay recurrence. For the growth-promoting effectors YAP1 and WWTR1, the direction is opposite to expectation. This is interpretable rather than contradictory: YAP/TAZ activity is set post-transcriptionally through phosphorylation and nuclear localization rather than transcript abundance, so bulk mRNA is a weak proxy for pathway output, and high transcript may instead mark an intact, feedback-competent Hippo circuit that still restrains growth. (Kaplan Meier curves in Fig 8A) PIK3CA (phosphoinositide kinase) is associated with lipid signaling, that feeds into the PI3K/AKT pathway. It showed the same counterintuitive direction (HR 0.62, p=0.043, DFS; KM log rank p = 0.022). (Fig 8B) This follows the same caveat: in prostate cancer, PI3K pathway activity is driven chiefly by PTEN loss and activating PIK3CA mutation rather than by PIK3CA transcript level, so expression might be an unreliable proxy for pathway activation.

**Figure 8.**
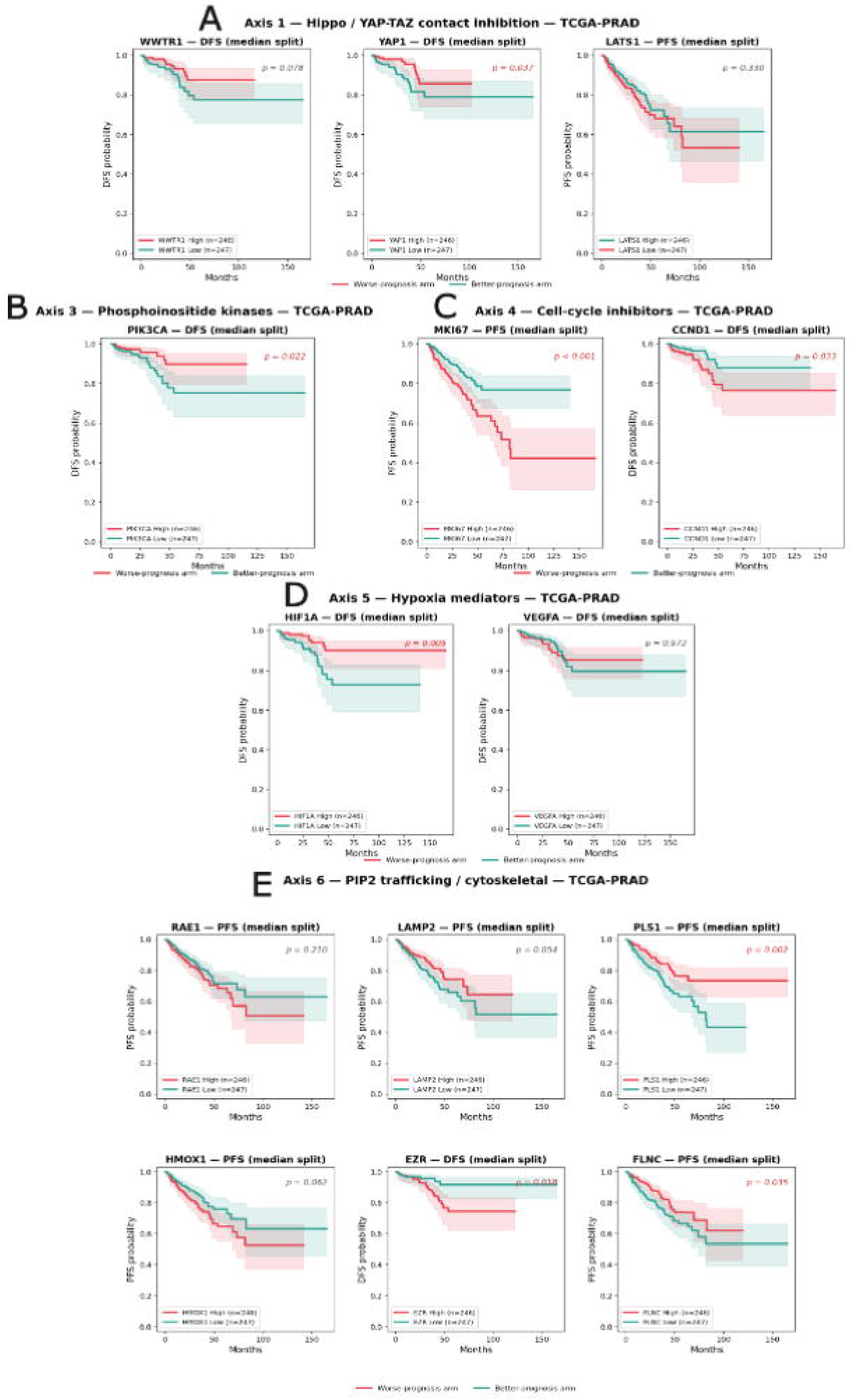
Clinical survival analysis of crowding/mechanotransduction genes mapped to the tissue-scale ABM contact-inhibition axis Kaplan-Meier survival stratification across five mechanobiology axes in TCGA-PRAD. Patients were dichotomized at the median expression (median split) of representative genes from each crowding-response axis (n = 246 high vs. 247 low), and disease-free survival (DFS) or progression-free survival (PFS) was compared between arms; red curves denote the worse-prognosis arm and teal the better-prognosis arm, with shaded 95% confidence bands and log-rank p-values shown per panel (significant values in red). (A) Axis 1 - Hippo/YAP–TAZ contact inhibition: WWTR1 (DFS, p = 0.078), YAP1 (DFS, p = 0.037), and LATS1 (PFS, p = 0.330). (C) Axis 3 - Phosphoinositide kinases: PIK3CA (DFS, p = 0.022), where, contrary to the prior, high expression marked the better-prognosis arm. (C) Axis 4 - Cell-cycle effectors/inhibitors: MKI67 (PFS, p < 0.001) and CCND1 (DFS, p = 0.033), both with high expression in the worse-prognosis arm. (D) Axis 5 - Hypoxia mediators: HIF1A (DFS, p = 0.005) and VEGFA (DFS, p = 0.972); high HIF1A was associated with better outcome, opposite to the expected direction. (E) Axis 6 - PIP2 interactome / cytoskeletal mechanosensing: RAE1 (PFS, p = 0.210), LAMP2 (PFS, p = 0.054), PLS1 (PFS, p = 0.002), HMOX1 (PFS, p = 0.062), EZR (DFS, p = 0.018), and FLNC (PFS, p = 0.035); note that high PLS1 marked the better-prognosis arm, opposite to the proteomic prior. Axis 2 (mechanosensitive channels; PIEZO1, TRPV4) is omitted, as no constituent gene reached nominal significance (p < 0.05) on DFS or PFS. Only genes reaching nominal significance are shown for each axis except where noted. All analyses use the TCGA-PRAD cohort. Log-rank p annotated per panel; HRs are univariate Cox per +1 SD z-score.

In cell-cycle arrest, CCND1 and MKI67 stand out. Cyclin D1 (CCND1), the gene whose product drives cell-cycle entry and thus directly opposes crowding arrest showed a strongly PTEN-dependent effect: high CCND1 trended toward worse recurrence (log rank p = 0.033, Fig 8C), but the effect was far larger in PTEN-Resistant tumors (per-standard-deviation hazard ratio 4.6 vs. 1.1 on DFS). This difference in effect size, captured as a gene × PTEN interaction, was the strongest result in the screen (interaction HR 5.40 on DFS, p = 1.6 × 10; HR 2.78 on PFS, p = 0.015): PTEN loss amplifies CCND1’s hazard, rather than simply adding to it. It matches prior work: Ju et al. (2014) ^42^ reported a cyclin D1 gene signature predicting worse recurrence-free survival; our screen extends this by showing the effect is specific to PTEN-deleted tumors, a PTEN-dependence not previously reported. Second, the proliferation marker Ki-67 (MKI67) marked worse PFS when high (log-rank p < 0.001 Fig 8C), confirming that the recurring tumors are the actively dividing ones. Thus, Cyclin D1, the direct effector of cell-cycle progression, becomes a markedly stronger marker of recurrence specifically when PTEN is lost, supported by the concordant Ki-67 proliferation signal. This links the two halves of the study: in the model, crowding physically holds back PTEN-deleted cells; in patients, once PTEN is lost, expression of the cell-cycle effector that overrides that brake most strongly marks the tumors that recur.

Higher HIF1A (hypoxia signature) was associated with better recurrence-free survival in the localized prostate cancer cohort (HR=0.046[0.27-0.82], p =0.008), opposite to literature reports showing high HIF1A predicts worst outcome in prostate cancer. ^43^ VEGFA expression showed no significant association with DFS in median-split Kaplan-Meier analysis (p = 0.972) but continuous z-score Cox regression showed significant results. (HR 1.39 [1.03–1.87], p = 0.030). (Fig 8D)

The PIP2 axis represents the PIP2 interactome, which is a set of proteins that physically associate with membrane lipid and couple the cell’s mechanical state to its cytoskeleton, membrane trafficking, and transport machinery. We include it because a prior proteomics study in prostate cancer cell lines^39^ linked several of these proteins to more aggressive phenotype. To extend on that observation, we studied their signatures at the mRNA expression level. Within the PIP2-trafficking/cytoskeletal signature, six individual genes reached nominal significance: RAE1, PLS1, EZR, LAMP2, HMOX1, and FLNC under per-standard deviation hazard ratio analysis or Kaplan Meier median split stratification: PLS1 (protective; DFS KM p = 0.025, Cox p = 0.055; PFS KM p = 0.002, Cox p = 0.018), RAE1 (PFS Cox p < 0.001, HR 1.36), FLNC (PFS KM p = 0.035, Cox p = 0.021), LAMP2 (PFS Cox p = 0.002, HR 0.70), HMOX1 (PFS Cox p = 0.006, HR 1.18), and EZR (DFS KM p = 0.018). (Table S6.4.1; Fig 8E). Out of these, PLS1, LAMP2, FLNC ran opposite to the screen’s expectation.

Beyond these, eight other gene × PTEN interaction terms reached nominal significance (p < 0.05): ANXA1, YAP1, CDKN1A, EPAS1, WWTR1, LATS2, and PIK3CA on PFS, and PLS1 on DFS (Table S6.4.2). (Fig S6.4.1) These run in the opposite direction from CCND1: their interaction HRs are all below 1 (range 0.24–0.55), meaning PTEN deletion attenuates rather than amplifies these genes’ prognostic effect. CCND1 is therefore the only gene in the screen where PTEN loss sharpens a pathway’s link to recurrence; for the rest, PTEN loss weakens it. Given the small PTEN-deleted arm (n = 85 PFS / 55 DFS), these eight interactions are reported as exploratory.

Taken together, the crowding axis is supported at the level of its terminal proliferative effectors, CCND1 and MKI67, whose mRNA reports proliferative output and which tracked recurrence as predicted, while its upstream mechanotransduction regulators might be better assessed by activity- or localization-based readouts than by bulk expression.

### Androgen Uptake and Synthesis Axis

In the ABM, the androgen resource sequestration axis is modeled through cellular testosterone uptake kinetics as a proxy for transporter-mediated androgen uptake. The candidate species evaluated clinically spanned both active uptake (SLCO2B1, SLCO1B3) and intratumoral *de novo* androgen synthesis (AKR1C3).

Among all the cohorts with sufficient mRNA expression data along with clinical survival data, only one cohort results in a significant hit. High AKR1C3 expression in TCGA-PRAD (primary localized disease) cohort was associated with poor progression-free survival (PFS) (log-rank p 0.038). (Fig 9A - i) In multivariate Cox regression adjusted for age, high AKR1C3 expression carried a 2.19-fold increased hazard of progression (HR = 2.19 [1.01–4.75]; p=0.046), approaching statistical significance for PFS. Furthermore, continuous z-score Cox regression in the TCGA-PRAD cohort for DFS yielded HR = 1.51 [1.25–1.83] (p < 0.001); in MSKCC (mixed prostate cancer cohort with DFS endpoint), the z-score threshold split yielded HR = 6.37 [2.17–18.73] (p < 0.001); in SU2C (metastatic CRPC with OS endpoint), the z-score split yielded HR = 5.89 [1.25–27.72] (p = 0.025); and in MCTP (metastatic prostate cancer), HR = 6.66 [1.65–26.97] (p = 0.008). (Supplement S6.2) This is consistent with the literature: AKR1C3 expression is significantly stronger in CRPC tissues than in matched hormone-naive specimens from the same patients, and high expression is an independent predictor of poor PSA progression-free survival after radical prostatectomy. AKR1C3 has also been identified as a driver of EMT through ERK signaling, and its upregulation is observed in enzalutamide-resistant cells where it reprograms AR/AR-V7 signaling. This pattern across biologically distinct cohorts (primary disease, CRPC, and metastatic) supports the clinical relevance of intratumoral androgen synthesis as a driver of disease progression.^44^

**Figure 9:**
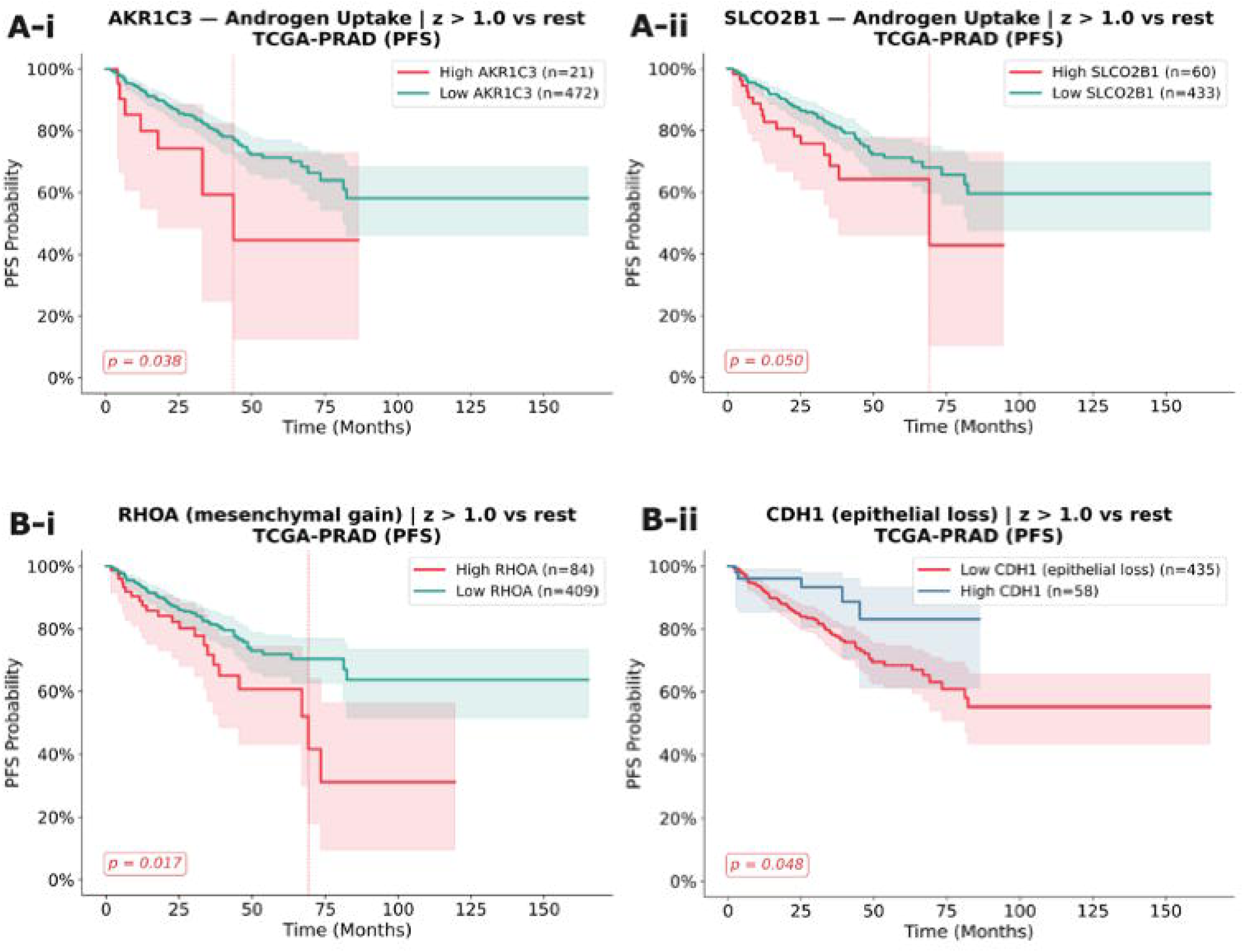
Clinical survival analysis of androgen uptake and synthesis genes mapped to the tissue-scale ABM testosterone-uptake axis. Kaplan-Meier survival analysis of mRNA expression in TCGA-PRAD (progression-free survival, PFS). Patients were stratified using a z-score threshold (z > 1 vs. rest). Dashed vertical lines indicate median survival where estimable. (A) Androgen uptake and synthesis axis: (i) AKR1C3 high expression was associated with significantly worse PFS (log-rank p = 0.038; N high = 21, N low = 472); (ii) SLCO2B1 high expression showed a borderline association with worse PFS (log-rank p = 0.050; N high = 60, N low = 433). (B) Adhesion-motility axis: (i) RHOA high expression was associated with significantly worse PFS (log-rank p = 0.017; N high = 84, N low = 409); (ii) CDH1 low expression, reflecting epithelial identity loss, was associated with significantly worse PFS (log-rank p = 0.048; N high = 435, N low = 58).

SLCO2B1 and SLCO1B3, the direct molecular proxies for the uptake rate parameter in the ABM, showed limited prognostic signal by mRNA expression. SLCO2B1 reached borderline significance in TCGA-PRAD using the PFS endpoint (z > 1 split; log-rank p = 0.05) (Fig 9A-ii), with a multivariate Cox HR (adjusted for age) of 1.66 [0.97 – 2.85] (p=0.066), approaching statistical significance. SLCO1B3 did not reach significance in any cohort or endpoint where z-score stratification was feasible. (TCGA-PRAD PFS: p=0.4; MSKCC DFS: p=0.991). (Supplement S6.2) Nevertheless, clinical literature supports their relevance: patients carrying both SLCO2B1 and SLCO1B3 genotypes that import androgens more efficiently exhibited a median two-year shorter time to progression on ADT in a cohort of 538 patients, and SLCO1B3 and SLCO2B1 are upregulated 3.6- and 5.5-fold respectively in metastatic CRPC tumors compared to untreated prostate cancer.^17^

### Adhesion-Motility Axis

In the ABM, adhesion-motility coupling governs the phase transition from a dispersed tumor architecture to a spatially clustered morphology, grounded mechanistically in PTEN-loss-driven suppression of E-cadherin via PI3K/AKT/Snail signaling and the corresponding acquisition of migratory phenotypes, characterized by upregulation of N-cadherin (CDH2), fibronectin (FN1), and cytoskeletal regulators including RHOA.

Clinical validation of this axis was performed across four mRNA cohorts spanning primary to metastatic disease, with mRNA z-scores evaluated for two epithelial retention markers (CDH1, EPCAM) and eight mesenchymal markers spanning EMT drivers (VIM, CDH2, SNAI1), ECM remodeling and invasion (MMP9, FN1), cytoskeletal motility regulators (RHOA, ITGB1), and pro-invasive signaling (CXCL8).

Among the ten genes evaluated, the two most directionally informative signals were RHOA, a cytoskeletal regulator of invasion whose high expression marked worse outcomes, and CDH1 (E-cadherin), an epithelial identity marker whose loss was associated with worse prognosis, both consistent with an EMT-driven invasive phenotype.

In TCGA-PRAD (primary localized disease), high RHOA expression was associated with significantly worse PFS under both KM stratification (log-rank p = 0.017) (Fig 9B-i) and multivariate Cox regression (HR = 1.80 [1.11–2.90]; p = 0.016). Low CDH1 expression, reflecting epithelial identity loss, was associated with worse PFS by KM (Fig 9B-ii) (log-rank p = 0.048), though the Cox HR did not reach significance after age adjustment (HR = 2.34 [0.95–5.77]; p = 0.065). In MSKCC (localized/early metastatic disease), FN1 and RHOA showed significant univariate Cox associations with worse DFS (FN1: HR = 3.97 [1.80–8.76], p = 0.001; RHOA: HR = 3.44 [1.32–8.93], p = 0.011); KM stratification was not feasible in MSKCC for these genes due to insufficient high-expression group sizes. In MCTP, MMP9 showed a significant Cox association (HR = 4.81 [1.24–18.71]; p = 0.023), though again without a supporting KM given the small cohort size. The remaining genes did not reach significance in any cohort, which is expected given the low mRNA-eligible patient counts in the metastatic subsets and the low OS event rate in the primary cohorts. The complete results are reported in Supplementary Section 6.3

These findings, when considered along with the established clinical literature on individual adhesion/motility relevant markers, indicate their importance in prostate cancer prognosis. Umbas et al. (1994) ^31^demonstrated that decreased E-cadherin expression correlated with poor prognosis in prostatectomy specimens. Gravdal et al. (2007) ^45^ showed by KM analysis that a switch from E-cadherin to N-cadherin expression in prostate tumor tissue microarrays correlated with recurrence. Wen et al. (2014)^46^ showed that Snail (SNAI1) expression independently predicted biochemical recurrence after radical prostatectomy, providing a mechanistic link between PTEN loss, CDH1 suppression, and clinical recurrence that parallels the coupling encoded in the ABM. In primary prostate cancer patients, reduced E-cadherin expression is significantly associated with shorter biochemical recurrence-free survival, and PI3K/AKT-mediated Snail activity has been functionally implicated as the mediating mechanism.^47^ RHOA has also been studied for its impact on androgen sensitivity in prostate cancer, with studies citing elevated RHOA expression associated with aggressive disease. ^48^

Taken together, these findings support that the pattern of mesenchymal marker gain and epithelial marker loss both associating with worse outcomes in independent cohorts is directionally consistent with the ABM’s predictions, lending clinical support to the adhesion-motility axis as a biologically relevant determinant of disease progression.

### Patient-Specific Gene Expression Profiles Stratify Individualized ADT Response in an Independent EUREKA trial dataset

The cohort-level analyses above demonstrate an axis of coupling between intrinsic genomic background and ADT-induced androgen scarcity determines collective recurrence trajectories across BR and TR patient groups. A key implication of this framework is that the same coupling logic should operate at the level of individual patients: a patient’s specific gene expression profile should produce a distinct proliferative response landscape under varying androgen conditions, and that landscape should predict their individual susceptibility to ADT resistance. To test this, we applied the MHS framework to 14 prostatectomized patients from the EUREKA1 registry ^28^, using per-patient differential gene expression profiles as model inputs.

We first assessed whether the model reproduces individual PSA dynamics. Longitudinal PSA data were available for three patients (two recurrent, one non-recurrent). Simulated PSA envelopes encompassed the observed values for all three patients. PSA trajectories predicted from the model produced clinically plausible longitudinal dynamics for recurrent and non-recurrent patients is presented in Supplementary Section S8.3. Simulated PSA envelopes encompassed the observed longitudinal values for all three patients. (Figure S8.3.1), The two recurrent patients were best described by an initial condition containing a pre-existing resistant clone, and the non-recurrent patient by a sensitive-only initial condition. The two-population MHS model therefore produces clinically plausible PSA dynamics under ADT, and the distinction between pre-existing resistant and sensitive-only initial conditions is the key determinant of recurrence trajectory.

We next applied the framework to the 14 prostatectomized patients with available gene expression data. For each patient, differentially expressed genes were mapped onto the corresponding MHS model species, yielding a patient-specific proliferative response landscape across androgen conditions. From this landscape we derived a single androgen sensitivity ratio per patient, presented in the following section.

### Androgen Sensitivity Ratio Stratifies Individual ADT Response Potential

To derive a single patient-level index of ADT response potential, we computed for each patient the ratio of NCG under high-testosterone conditions to NCG under low-testosterone conditions (separately for S (PTEN-normal) and R (PTEN-deleted) cell populations) (Figure 10). This ratio quantifies the degree to which a patient’s tumor cells depend on androgen availability for net growth: a high ratio indicates strong androgen dependence, consistent with expected sensitivity to ADT-induced androgen withdrawal; a low ratio, or a ratio near unity, indicates relative androgen independence, consistent with expected intrinsic resistance.

**Figure 10.**
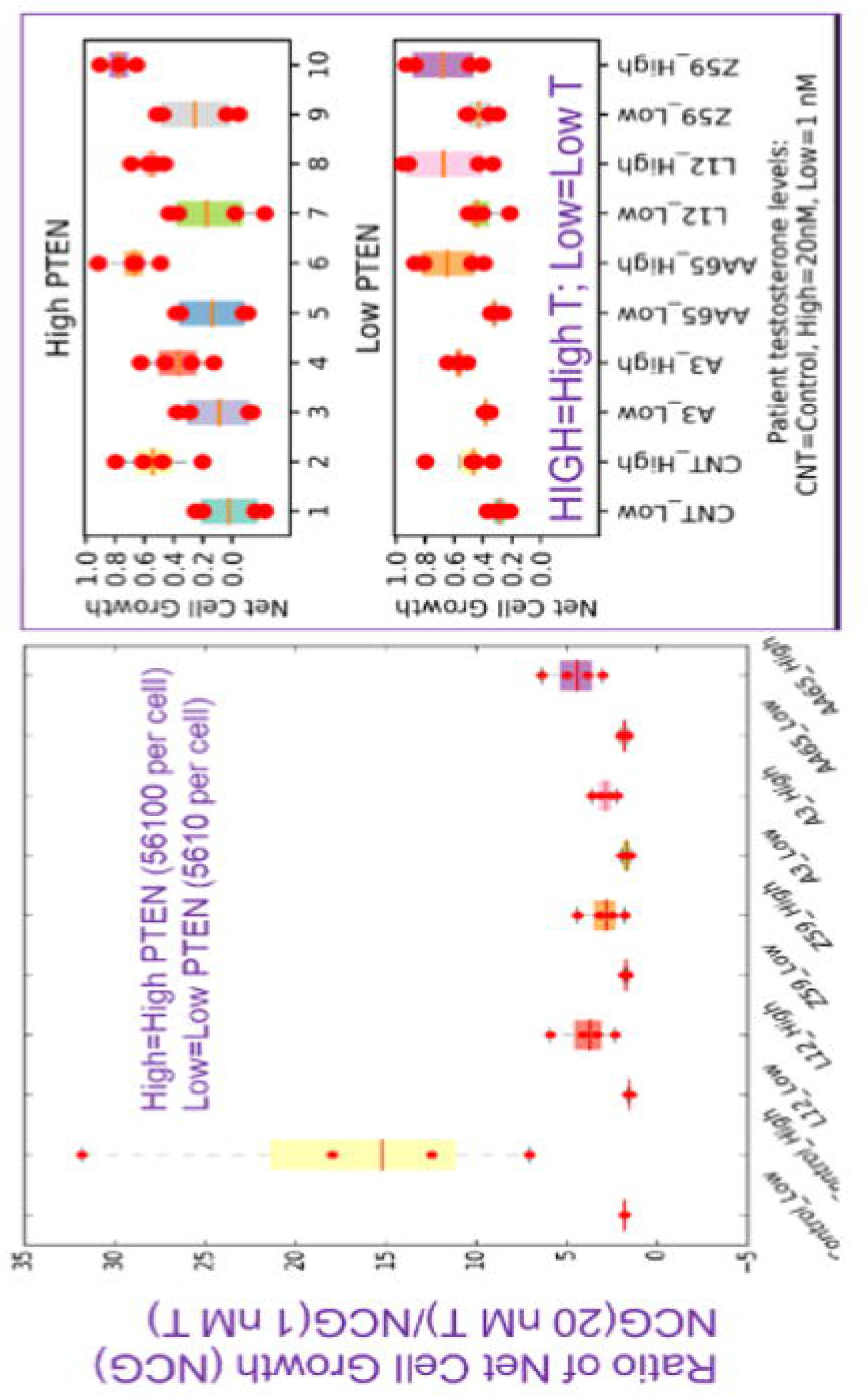
Patient-specific androgen sensitivity ratios stratify EUREKA1 patients by model-predicted androgen dependence. Patient-specific androgen sensitivity ratio (NCG at high testosterone / NCG at low testosterone) for S cells (top) and R cells (bottom) for 4 representative patients and the healthy control. A ratio near 1 indicates androgen independence; a high ratio indicates androgen dependence and expected sensitivity to ADT. DGEP-Y patients (those with differentially expressed genes) are highlighted.

Among DGEP-Y patients, the high-T/low-T NCG ratio was substantially more variable than among DGEP-N patients, whose ratios clustered near the control value. Critically, the R cell population (low PTEN) displayed consistently lower ratios than the corresponding S cell population (high PTEN) across all patients, confirming that PTEN deletion shifts the proliferative response function toward androgen independence regardless of the broader expression background, but that the magnitude of this shift is patient-specific. Patients with the lowest R-cell ratios (near-equal NCG under high and low androgen) represent the subgroup at highest risk of early CRPC onset under standard ADT. are the subgroup the model nominates as most androgen-independent, and therefore the candidate group for early CRPC onset under standard ADT.

### From Population Model to Personalized Prediction

Taken together, these results extend the framework from the cohort level to the individual patient. The same gene-expression-guided parameterization used to define the BR, TR, and control cohorts was applied to single patients, mapping each patient’s differentially expressed genes onto the corresponding species in the cellular model. This produced patient-resolved net cell growth profiles and, from the ratio of net growth at high versus low testosterone, a per-patient measure of androgen dependence. Sensitive populations showed two- to five-fold testosterone-dependent growth, while resistant populations showed little testosterone dependence, and this separation varied in magnitude across patients. While we did not repeat the intrinsic to extrinsic coupling to scale the simulation to agent-base simulations, the intrinsic heterogeneity resolved in this analysis would be a natural entry point for patient-specific tissue-scale simulations if relevant biomarkers are available to parametrize the tissue-scale model. The androgen-sensitivity ratio is therefore a model-derived stratifier of androgen dependence, not a validated predictor. It has not been tested against observed ADT outcomes in these patients, and prospective correlation with clinical response is required before predictive use.

## Discussion

This study was motivated by a central clinical question: why do prostate cancer patients with seemingly similar disease at the time of radical prostatectomy follow such divergent trajectories, from indolent biochemical recurrence to rapidly castration-resistant disease? The hypothesis that helped shape our modeling framework was that divergent clinical outcomes are not pre-determined by cell-intrinsic genetic differences or extrinsic microenvironmental pressures in isolation but emerge from their dynamic coupling. The results presented here support this hypothesis across three interconnected biological dimensions: the physical architecture of residual tumor tissue, the dynamics of androgen resource competition, and the spatial self-organization of genetically distinct clones. In each case, the same intrinsic genetic background produced qualitatively different tissue-level outcomes depending on the microenvironmental context, and the same environmental pressure produced divergent clonal selection outcomes depending on patient-cohort-specific intrinsic signaling properties. Neither genetic heterogeneity nor microenvironmental variation alone is sufficient to explain what we observe; it is their coupling that is explanatory.

Biochemical recurrence following radical prostatectomy occurs in approximately 20–30% of patients within a decade of surgery ^3^, yet the subsequent disease course varies enormously. Current clinical risk stratification relies heavily on serum PSA kinetics, which cannot distinguish mechanistically between indolent local recurrence and the early stages of systemic, treatment-resistant progression. This gap has real consequences: patients with occult aggressive disease risk undertreated progression, while others with indolent recurrence may be subjected to systemic therapies carrying significant morbidity. Developing frameworks that capture the mechanistic drivers of this divergence, rather than describing it statistically, is therefore a priority for precision oncology in prostate cancer. ^49^

### Cell-Intrinsic Heterogeneity Operates at Two Levels, namely Pathway Dosage and Genomic Background

Our cellular-scale results establish that PTEN status is a primary molecular determinant of recurrence potential, with higher PTEN deletion levels driving earlier biochemical recurrence and more aggressive PSA escalation under simulated androgen deprivation therapy. This dose-dependent relationship reflects the graded, constitutive activation of the PI3K/AKT pathway that PTEN loss confers, directly attenuating apoptotic sensitivity to androgen withdrawal. Importantly, this relationship held across BR and TR cohorts, confirming that PI3K/AKT pathway dosage is a conserved driver of recurrence potential. However, PTEN status alone does not capture the full picture. At equivalent high PTEN deletion levels, the TR cohort exhibited substantially more aggressive PSA kinetics than the BR cohort, demonstrating that the broader genomic background of each patient population amplifies or constrains the consequences of individual pathway perturbations. This is an important conceptual refinement: PTEN loss provides the molecular ignition for recurrence through PI3K/AKT-driven androgen independence, but the rate and severity of progression are conditioned on the background signaling state of the incorporated pathways, which differs systematically between clinically defined cohorts. Cell-intrinsic heterogeneity therefore operates at two levels, the dosage of individual pathway components, and the genome-specific signaling context in which those components operate.

### Convergent Identification of PTEN, MDM4, and AR as Clinically Actionable Molecular Drivers

To identify which molecular species most strongly determine cell fate across this complex signaling landscape, we applied a computational feature-ranking strategy to an efficient surrogate of our full mechanistic model. Across three independently trained classifier architectures, PTEN, MDM4, and AR emerged as the dominant intrinsic determinants of proliferative versus quiescent cell fate, a consensus that is robust to the specific modeling choices made for each classifier. The consistent identification of MDM4 is biologically notable. As a primary negative regulator of p53, MDM4 overexpression suppresses apoptotic signaling downstream of genotoxic and oncogenic stress; its upregulation is increasingly recognized as a driver of resistance in castration-resistant prostate cancer (CRPC). ^50,51^ Its emergence alongside PTEN and AR, the canonical genomic drivers of prostate cancer progression points to the p53-MDM4 axis ^52^ as a potentially underappreciated determinant of treatment resistance and an actionable therapeutic target.

The clinical relevance of these SHAP-identified drivers was directly validated by Kaplan-Meier survival analysis in cBioPortal prostate cancer patients. Patients with alterations in PTEN, MDM4, or AR exhibited significantly worse overall survival than those without such alterations (p < 0.05 by log-rank test), establishing a direct correspondence between model-predicted determinants of cell fate and patient prognosis. While this correspondence does not by itself establish causal directionality, the convergence of computational feature importance and clinical survival outcomes across an independent patient dataset substantially strengthens the case that these pathway nodes are capturing biologically meaningful disease biology, not modeling artifacts. Together, these findings establish PTEN, MDM4, and AR as a clinically actionable molecular signature of CRPC risk.

### The Post-Surgical Tissue Architecture Determines How Much of a Tumor’s Intrinsic Aggressive Potential Is Realized

Having established the intrinsic genetic determinants of recurrence potential, we asked how the physical environment of residual tumor tissue modulates the expression of that potential. PTEN-deleted prostate cancer cells possess constitutive resistance to androgen withdrawal-induced apoptosis. If such cells persist following prostatectomy, the question is not whether they have the molecular capacity to drive recurrence, but whether the physical context in which they find themselves allows that capacity to be realized.

Our tissue-scale simulations demonstrate that it is context-dependent. In physically unconstrained environments with low cellular packing density, the aggressive growth advantage of PTEN-resistant clones was amplified: high initial resistant-to-sensitive cell ratios translated directly into rapid tumor expansion. In contrast, densely packed environments imposed contact inhibition that substantially dampened this intrinsic advantage, even at identical resistant clone frequencies. This behavior was consistent across BR and TR cohorts. Intrinsic genotype sets the ceiling of aggressive potential; the physical permissiveness of the tissue microenvironment determines how much of that potential is realized. This finding carries a direct clinical implication. Following radical prostatectomy, the physical state of residual tissue, including surgical margin status, local fibrosis, and stromal remodeling, may be as important as the genetic composition of any surviving tumor cells in determining the pace of recurrence. Clinical evidence that tissue architecture independently stratifies recurrence risk beyond Gleason grading has been demonstrated through quantitative histological scoring of needle biopsies. ^53^ However, such approaches are descriptive and cannot explain why certain architectural configurations confer higher risk. The mechanistic framework presented here offers a potential biological basis for why tissue architecture features carry prognostic weight, linking packing density directly to the realized proliferative advantage of resistant clones. These results together suggest that tissue architecture constitutes a class of microenvironmental determinants of recurrence risk that merits prospective clinical investigation.

### Cohort-Specific Androgen Sensitivity Determines Whether Resource Competition Accelerates or Preserves Tumor Heterogeneity

Perhaps the most direct demonstration of intrinsic–extrinsic coupling emerges from the androgen resource competition results. Under androgen deprivation therapy, prostate tumors must survive in an environment of systemic androgen scarcity. One established mechanism of resistance is the upregulation of SLCO membrane transporters (SLCO2B1, SLCO1B3), which support active intracellular androgen uptake, maintaining proliferative signaling even at low systemic concentrations, and further depleting the androgen available to neighboring cells. ^17^ This self-induced resource depletion generates a shared environmental pressure on mixed tumors, but whether that pressure selectively eliminates androgen-sensitive clones depends on those clones’ intrinsic androgen sensitivity.

In the BR cohort, androgen-sensitive cells exhibited strong nonlinear dependence on testosterone availability. As resource sequestration reduced local androgen concentrations from physiological to intermediate levels, the proliferative capacity of PTEN sensitive cells in BR cohort declined sharply, falling well below that of resistant clones which maintained robust growth at intermediate androgen concentrations. This differential created a broad competitive window in which resistant clones progressively outcompeted their sensitive counterparts, converting a heterogeneous mixed tumor into a predominantly resistant one. Moderate-to-high androgen uptake therefore acted as a selective pressure that enriched the PTEN-resistant population in BR tumors. The TR cohort behaved in a fundamentally different way under identical environmental pressure. The PTEN-sensitive cells in TR cohort maintained robust proliferation at androgen concentrations well below the threshold that suppressed their BR counterparts, with their proliferative threshold shifted to extremely low concentrations. Consequently, even under conditions of high uptake-driven androgen depletion, TR-sensitive cells continued to proliferate alongside resistant clones, preventing the competitive dominance of the resistant population and maintaining intratumoral heterogeneity even under significant resource stress. This divergence offers a mechanistic perspective on why some patients with mixed-genotype tumors progress rapidly toward treatment-resistant disease while others, under comparable conditions, maintain tumor heterogeneity with slower overall progression. These simulations suggest the answer lies not in environmental pressure alone, nor in intrinsic signaling state alone, but in their coupling.

### EMT-Associated Adhesion Loss Drives Spatial Tumor Reorganization Through a Threshold-Dependent Mechanism

Beyond resource competition, the spatial organization of genetically distinct clones within the tumor mass is itself a clinically relevant feature of disease progression. The Epithelial-Mesenchymal Transition (EMT) is a key program through which prostate cancer cells shed adhesive constraints (primarily through loss of E-cadherin) and acquire invasive mesenchymal properties. Reduced E-cadherin expression is a hallmark of aggressive prostate cancer and is associated with poor prognosis. ^31^ Importantly, constitutive PI3K/AKT activation downstream of PTEN loss suppresses E-cadherin expression through transcriptional repressors including Snail ^32^, directly linking the genetic alterations captured in our cellular model to the adhesion-motility phenotype of cells in the tumor tissue. PTEN status therefore does not only determine a cell’s intrinsic proliferative potential; it also determines how that cell physically interacts with its neighbors and with the spatial architecture of the tumor.

Our tissue-scale simulations reveal that variations in cell–cell adhesion strength had negligible impact on overall tumor burden but drove dramatic divergences in tumor composition (redistributing clonal competitive advantage) and spatial organization. At low to moderate adhesion levels, motile forces dominated, and resistant cells dispersed freely throughout the tumor, producing a spatially mixed architecture. Beyond a critical adhesion strength, however, the balance shifted: adhesive forces drove the progressive self-organization of resistant cells into dense, spatially segregated clusters that excluded their sensitive neighbors. This transition from dispersed to clustered morphology occurring over a narrow range of adhesion values indicates that spatial reorganization of the tumor is a threshold-dependent biological phenomenon governed by the coupling of intrinsic adhesive state (set by PTEN-driven PI3K/AKT activity) and the mechanics of physical cell–cell contact. Extending this analysis across cohorts revealed that the adhesion-driven transition is not universal but patient cohort-specific. Control cells clustered most readily, requiring the least adhesion to trigger spatial segregation. BR cells required progressively higher adhesion, with this threshold rising further under low androgen conditions. TR cells were the most resistant to clustering, showing no clear transition at nominal androgen levels and only partial segregation at high adhesion. This ordering reflects a progressively motility-dominant phenotype across recurrence severity.

### Limitation

Multiscale mechanistic models of cancer progression inevitably require scope decisions. Capturing every known biological axis simultaneously leads to models that are computationally intractable and increasingly difficult to parameterize, validate, or interpret. The limitations described below define the scope of the results and clarify what biological information was not directly captured by the model.

First, the cellular model captures PTEN-driven PI3K/AKT dysregulation and AR signaling as the primary resistance mechanisms, but cancer cell-intrinsic heterogeneity in prostate cancer has several additional well-documented sources that are not directly represented. Prostate tumors are commonly multifocal, with multiple spatially separate lesions arising from independent transformation events. Up to 70% of multifocal cases show that some lesions carry the TMPRSS2-ERG fusion while others do not, confirming that different parts of the same gland can harbor cancer cell populations with fundamentally different genetic identities. ^54,55^ Within individual lesions, ongoing subclonal evolution generates further diversity: TP53 mutations arise as low-frequency subclonal events in primary tumors and subsequently drive selective expansion of subpopulations with greater metastatic potential and distinct therapy-response profiles. ^56^ Beyond genetic heterogeneity, epigenetic variation through differential DNA methylation at AR target loci creates phenotypically distinct cancer cell states among genetically identical cells, and these states are reversible and shift under therapeutic pressure. ^57,58^ AR splice variants such as AR-V7 further confer ligand-independent receptor activity, and intratumoral androgen synthesis via AKR1C3 upregulation maintains intracellular androgen levels despite systemic suppression. The practical consequence is that any patient’s tumor contains a mixture of cells with different AR dependencies, proliferative rates, and apoptotic thresholds. The current two-population PTEN-based framework captures one axis of this heterogeneity but does not account for the additional layers of genetic, subclonal, and epigenetic variation that collectively shape population-level tumor behavior in vivo. Two additional resistance axes warrant explicit mention as important mechanisms in disease progression: differential SLCO transporter expression and glucocorticoid receptor bypass after AR blockade. SLCO membrane transporters ^59^ actively import androgen precursors from the circulation into tumor cells, and their differential expression across cancer cell subpopulations creates heterogeneous intratumoral androgen gradients that our agent-based model is specifically designed to resolve spatially. When AR is pharmacologically blocked, GR becomes derepressed and drives a partially overlapping set of AR target genes, effectively bypassing androgen deprivation without any new AR mutation. ^60^ High GR expression is independently associated with advanced tumor stage, high Gleason grade, nodal metastases, and early biochemical recurrence in prostate cancer. ^61^ Unlike the fixed genetic events captured in the current framework, GR upregulation is a reversible adaptive response whose prevalence shifts dynamically with treatment intensity, making it a particularly interesting target for future agent-based extensions that model cell-state plasticity under therapeutic pressure.

Second, the tissue simulations are two-dimensional, which is sufficient to demonstrate coupling between intrinsic cell state and microenvironmental mechanics but does not fully recapitulate three-dimensional tumor architecture. Contributions from the stroma, vasculature, and immune infiltrate remain unrepresented at the tissue scale. The tumor stroma is not a passive scaffold but an active participant in prostate cancer progression. Cancer-associated fibroblasts (CAFs) are the dominant stromal cell type and exist in functionally distinct subtypes: inflammatory CAFs (iCAFs), which predominate in hormone-sensitive disease, and myofibroblastic CAFs (myCAFs), which are enriched in high-grade and castration-resistant disease and linked to poor prognosis. ^62^ CAFs remodel the extracellular matrix, secrete growth factors including TGF-β, HGF, and FGF, and establish paracrine signaling loops that directly stimulate cancer cell proliferation, invasion, and therapy resistance. Critically, ADT does not simply suppress cancer cells, it actively reprograms the stroma. Antiandrogen treatment unleashes TGF-β signaling that drives iCAF-to-myCAF phenotype switching, and the resulting SPP1+ myCAFs render cancer cells refractory to ADT, establishing the stroma as a direct driver of castration resistance rather than a bystander. ^63^ Loss of AR signaling in CAFs further promotes CCL2 and CXCL8 secretion that enhances cancer cell migration, meaning that ADT simultaneously removes a stromal restraint on invasion. ^64^ The physical consequence of CAF activation is progressive ECM stiffening through collagen deposition and crosslinking, which is sensed by cancer cells through integrin-focal adhesion kinase signaling that activates downstream PI3K/AKT and MAPK/ERK pathways, reinforcing the same proliferative and survival programs that PTEN loss engages intracellularly. ^65^ This creates a mechanical amplification loop where genetic vulnerability and physical microenvironmental pressure converge on the same oncogenic output. Beyond the stroma, the immune microenvironment represents an equally important and currently unrepresented axis. Prostate cancer is immunologically cold, characterized by sparse effector T cell infiltration and a microenvironment dominated by M2-polarized macrophages, MDSCs, and Tregs that collectively suppress CD8+ cytotoxic activity through IL-10, TGF-beta secretion, and immune checkpoint expression. ^66,67^ ADT transiently activates immune infiltration before a compensatory immunosuppressive state is re-established. ^68,69^ Existing prostate-specific computational models have begun addressing this: an ABM of M1/M2 macrophage polarization under ADT found that macrophage reprogramming can paradoxically enhance tumor survival ^70^, and a 3D hybrid multiscale model coupling CTL, Treg, and TAM dynamics with androgen signaling identified immunosuppression as a key CRPC driver. ^71^

Additionally, cohort parameterization relied on TCGA bulk tumor sequencing, which averages over spatially heterogeneous cell populations and may obscure subclonal dynamics. Single-cell and spatial RNA sequencing data ^72^, where available, could enable more granular parameterization at the level of individual cell populations. A further limitation concerns validation of the tissue-scale outputs. The ABM was verified for internal consistency against the two-population ODE submodel under spatially unconstrained conditions. (Fig S7.3.1) The ABM was validated only for internal consistency against the ODE submodel under spatially unconstrained conditions. Its spatial outputs, the clustering index and R/S tissue composition, were not compared against immunohistochemistry, spatial transcriptomics, or quantitative histology, and are therefore predictions rather than validated descriptions of tissue architecture. The predicted homotypic segregation of resistant clones is qualitatively consistent with documented subclonal spatial separation in multifocal prostate tumors ^54–56^, but direct validation would require spatially resolved data pairing clonal identity with cell position.

### Future Directions

Three directions follow directly from the limitations of the current framework. First, the cellular model should be extended to capture a broader landscape of intrinsic resistance mechanisms. The current formulation centers on PTEN-driven PI3K/AKT dysregulation as the primary driver of androgen independence. However, AR splice variants such as AR-V7 confer ligand-independent receptor activity that renders cells constitutively resistant regardless of androgen availability. Intratumoral androgen synthesis via CYP17A1 and AKR1C3 upregulation maintains intracellular androgen levels that sustain proliferation even under profound systemic suppression. Glucocorticoid receptor bypass represents a further escape mechanism in heavily pretreated disease.

Second, pharmacokinetic and pharmacodynamic models of second-generation antiandrogens can be coupled to the current framework. Agents such as enzalutamide, abiraterone, apalutamide, and darolutamide are now standard of care across multiple disease states, and PARP inhibitors such as olaparib and rucaparib are increasingly used in genomically selected patients. Emerging theranostic agents including 177Lu-PSMA-617 represent a further axis of treatment whose mechanistic integration into predictive models remains largely unexplored. ^73^ Coupling PK/PD representations of these agents to the MHS framework would enable in silico prediction of combination therapy responses, resistance emergence under sequential treatment, and patient-specific treatment sequencing strategies.

Third, the tissue-scale model should be extended to three dimensions with explicit representation of the stromal and immune compartments. Cancer-associated fibroblasts drive progression through ADT-induced iCAF-to-myCAF switching and ECM stiffening that converges on the same PI3K/AKT and MAPK/ERK programs engaged by PTEN loss. On the immune side, ADT-induced shifts in macrophage polarization and T cell infiltration create dynamic immunosuppressive feedback loops that warrant explicit representation in a three-dimensional spatial framework. Incorporating these stromal and immune interactions into a three-dimensional spatial framework would open the model to mechanistic investigation of immunotherapy and ADT combination strategies. As these modeling efforts mature, prospective validation using patient-derived organoids, which preserve patient-specific genomic context in a controlled experimental system, will be essential for directly testing the coupling hypotheses that emerge from each extended framework.

In summary, this study presents a mechanistic multiscale framework that translates static clinical genomic data into dynamic predictions of prostate cancer tumor evolution. By integrating cell-intrinsic signaling heterogeneity with the physical and metabolic constraints of the tumor microenvironment, the model provides a quantitative explanation for the divergent recurrence trajectories observed across Control, BR, and TR patient cohorts. The central finding is that recurrence is not a genetically predetermined outcome but an emergent property of how intrinsic clonal fitness interacts with tissue architecture, androgen resource competition, and the spatial mechanics of clonal self-organization. The microenvironment is an active participant in CRPC emergence, not merely a backdrop to it. Precision medicine for prostate cancer must therefore extend beyond genomic profiling to encompass the environmental context in which those genomic alterations operate. The framework developed here provides a platform for exploring how diverse microenvironmental pressures interact with patient-specific biology to determine whether a latent genetic risk remains dormant or progresses to aggressive disease.

## Methods

### Data Sources and Patient Cohort Definition

Tumor sequencing and clinical data for prostate cancer patients were obtained from The Cancer Genome Atlas (TCGA; http://cancergenome.nih.gov). Patients were characterized by biochemical recurrence status (occurrence and timing), adjuvant ADT receipt, PTEN deletion status, pathological Gleason score, surgical margin status, and tumor status assessed by biopsy and imaging. From an initial cohort of 501 patients, those who received adjuvant or neo-adjuvant radiotherapy were excluded, yielding a final analytical cohort of 250 patients for whom complete clinical information was available.

Patients were stratified in two stages. First, by PTEN deletion status (deleted vs. normal). Second, within each PTEN group, by ADT response: a Control cohort (CNT; no recurrence), a Biochemical Recurrence cohort (BR; PSA-defined recurrence), and a Tumor Recurrence cohort (TR; image-based determined recurrence). Fig 11 represents the flowchart of this cohort selection. Differentially expressed genes (DEGs) between BR or TR and CNT were identified using the R2 Genomics platform (http://r2.amc.nl) on the TCGA dataset. Initial concentrations of corresponding model species were scaled to reflect cohort-specific expression profiles, producing the genomically parameterized cohort instances used throughout the study. Mutation frequency data from cBioPortal informed the biological scope of the model, with genes mutated in >5% of patients prioritized as candidate pathway nodes.

**Figure 11.**
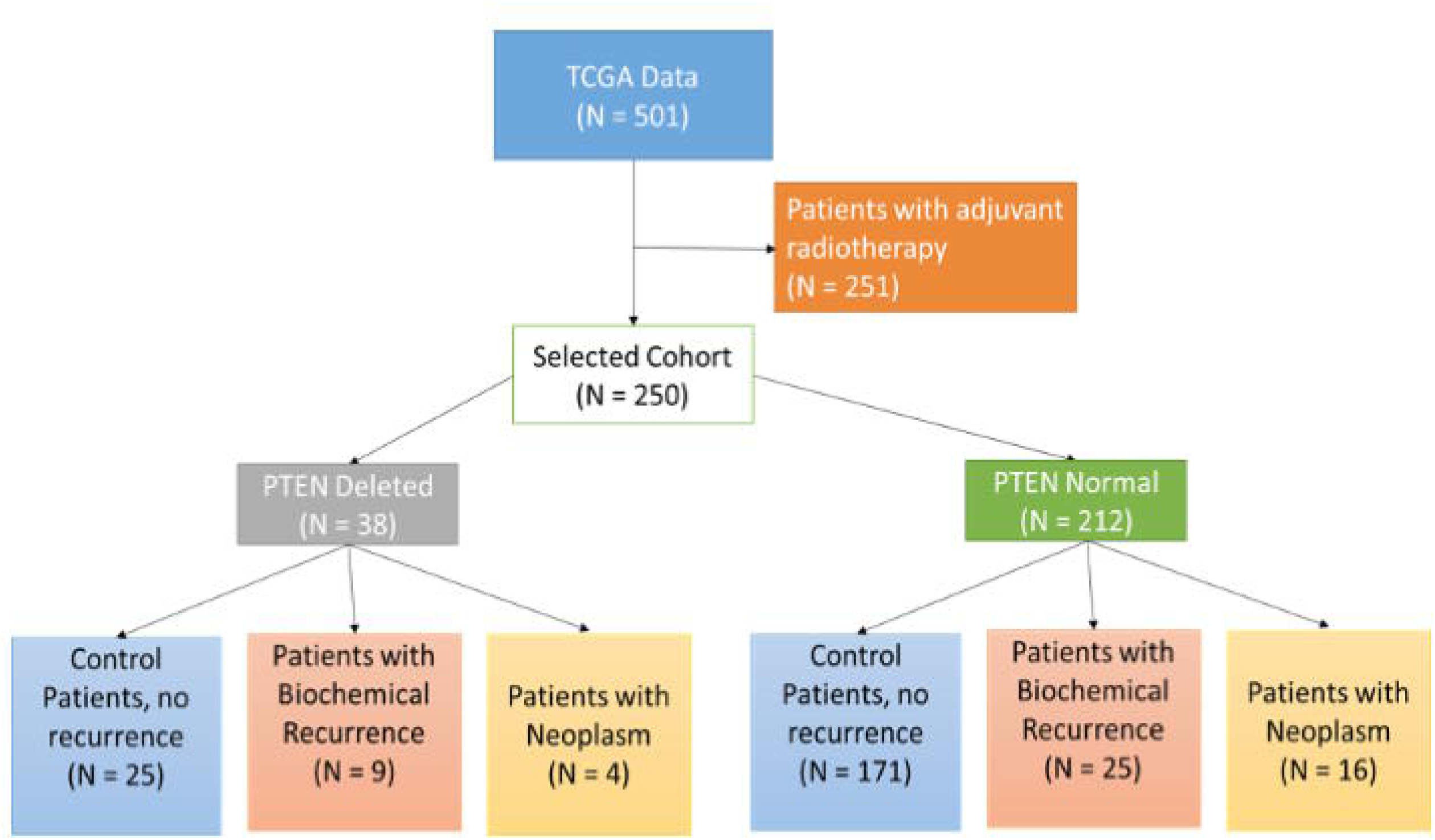
Flow chart of the cohort selection: stratification of TCGA-PRAD patients into Control, BR, and TR groups by PTEN deletion and recurrence status From the initial 501 patient samples, patients who underwent adjuvant radiotherapy were first excluded. This cohort of the patient was further classified into two groups based on whether or not PTEN deletion mutation was present. Each of these groups was further classified as control or patients with no recurrence, patients with biochemical recurrence (PSA values), and patients with tumor recurrence.

In addition to these publicly available datasets, we also attempted using this model on 14 patients in the EUREKA1 trial cohort. This data was provided by collaborators Caterina Guiot and ilaria Sturia at University of Torino, Italy. These patients have been anonymized and their genomic data has been used in this study.

### MHS Cellular Model

To investigate how cell-intrinsic genetic alterations determine recurrence potential and response to ADT, we developed the Prostate Cancer Multiscale Hybrid Systems (MHS) model. The model integrates five submodels representing key signaling pathways operating across distinct biological timescales (Fig 12): (1) AR pathway with EGFR-mediated Ras–MAPK signaling; (2) PI3K-AKT pathway incorporating PTEN status; (3) androgen biosynthesis and receptor activation; (4) a Boolean submodel of TP53-mediated cell cycle gene transcription; and (5) a two-population PSA dynamics submodel representing androgen-sensitive (S) and androgen-resistant (R) tumor cells. Cross-talk between the AR and PI3K/AKT pathways, and their shared interaction with p53, is a primary mechanism of ADT resistance and is explicitly encoded in the model architecture. ^15^

**Figure 12:**
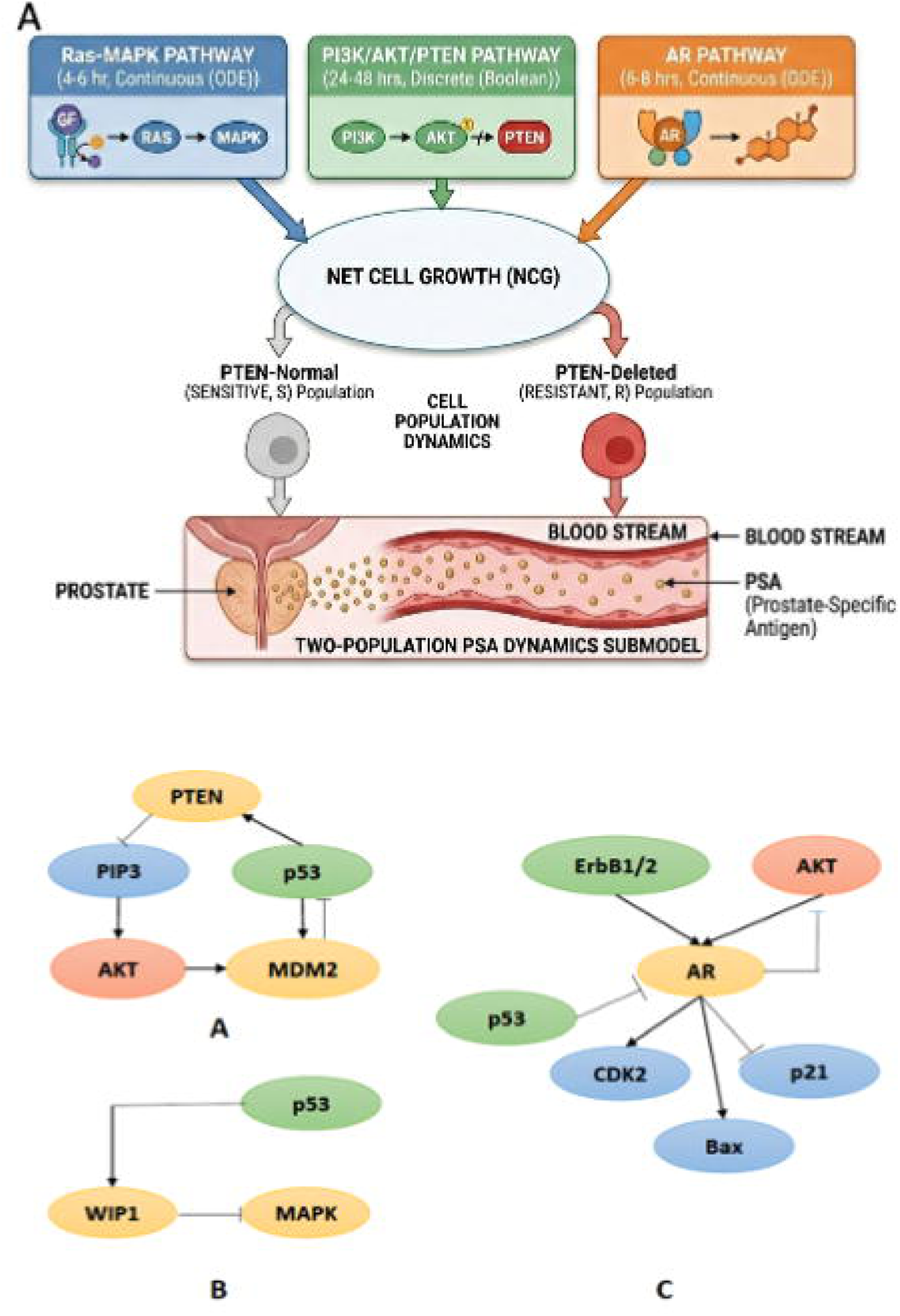
Overview of prostate cancer multiscale hybrid systems (MHS) model structure The MHS model integrates three submodels encoding key signaling pathways across distinct biological timescales to predict cell fate and PSA dynamics under ADT. (Model Architecture) Schematic showing how the growth factor (GF)-mediated Ras-MAPK, PI3K/AKT/PTEN, and AR pathway submodels converge to produce net cell growth (NCG) outputs for PTEN-normal (sensitive, S) and PTEN-deleted (resistant, R) cell populations, which in turn drive the two-population PSA dynamics submodel. (Pathway Interfaces) Module interfaces identified from literature defining the cross-talk encoded in the model architecture: (A) interface between the PI3K/AKT and p53 pathways, (B) interface between the Ras-MAPK and p53 pathways, and (C) interface between the AR pathway and the Ras-MAPK and p53 pathways. Abbreviations: AR, androgen receptor; GF, growth factor; NCG, net cell growth; PTEN, phosphatase and tensin homolog; PSA, prostate-specific antigen; ADT, androgen deprivation therapy; S, androgen-sensitive population; R, androgen-resistant population.

The molecular-level ODE submodels (1-3) operate on timescales of 4 - 8 hours; the Boolean p53 module operates on timescales of 24 - 48 hours. A hybrid simulator algorithm bridges these timescales by running faster ODE modules to steady state and passing discretized outputs to the Boolean module, which in turn returns bounded continuous values to the ODE layer. ADT was simulated by varying testosterone concentration from 1 to 20 nM, spanning the range of published pharmacokinetic data from PCa patients.^74^ The model was implemented and simulated in COPASI ^75^ and Python v2.7. Full equations, parameter values, and initial conditions for each separate sub model are provided in Supplementary Section S1.1, S1.2, S1.3. The methodology for integration of these sub models across biological time scales is described in Supplementary Section S2.

To represent cohort-level intrinsic heterogeneity, each cohort instance was simulated across two PTEN expression levels, five testosterone concentrations, and two intra-tumoral heterogeneity conditions (high and low GF levels), yielding 20 simulation instances per cohort. Net cell growth (NCG), which is the difference between simulated cell growth and kill probabilities, served as the primary cellular output and was parameterized as a Hill function of testosterone concentration separately for S and R populations (Supplementary Section S1.4 Two-population sub model). Serum PSA was computed as: PSA(*t*) = scoeff × S(*t*) + rcoeff × R(*t*), with per-cell secretion coefficients drawn from published values. ^76^

The two-population sub model was validated through two complementary approaches detailed in Supplementary Section S5: qualitative validation against the published pERK-pAKT response map of Chen and colleagues, confirming that the integrated sub model architecture reproduces the correct cell-fate boundaries between proliferating and quiescent states; and quantitative identification of the NCG classification threshold via three independently trained supervised classifiers (SVM, Decision Tree, and NN), which converged on NCG = 0.15 as the optimal boundary separating androgen-sensitive from androgen-resistant phenotypes across all cohorts. This threshold is used as the classification boundary throughout all subsequent analyses.

### Agent-Based Model

To investigate how intrinsic clonal fitness interacts with microenvironmental architecture to produce tissue-level outcomes, the MHS cellular model was embedded (via the two-population sub model) within a spatial agent-based modeling (ABM) framework implemented in PhysiCell (v1.10.4) ^77^, an open-source multi-cellular simulator that couples intracellular signaling with diffusive substrate transport via BioFVM. ^78^ The ABM operates in 2D and resolves individual cell dynamics - proliferation, migration, adhesion, and death - alongside continuous gradients of testosterone and mitogen substrates.

To ensure that tissue-scale ABM outputs could be interpreted as biologically meaningful spatial effects rather than numerical artifacts of the modeling framework, we verified consistency between the ABM and the two-population ODE sub model under equivalent, spatially unconstrained conditions, confirming that PSA outputs from five stochastic ABM replicates fell within one standard deviation of ODE-predicted values across all three cohorts at both low (1 ng/mL) and high (8 ng/mL) testosterone concentrations. (Supplementary Section S7)

Simulations were initialized with 150 cancer cells (S and R subtypes in defined R/S ratios) placed randomly in a circular arrangement under normoxia and physiological androgen concentrations. MHS-derived proliferation, migration, and adhesion propensities were assigned to each agent according to its genotype (S or R) and the local microenvironmental state, which in turn was updated by the agents’ substrate uptake and secretion. This bidirectional coupling (intrinsic genotype shaping environmental state, and environmental state modulating clonal competitive outcomes) is the structural implementation of the central hypothesis. Three microenvironmental axes were systematically varied: spatial packing density (low vs. high crowding), androgen uptake rate (as a proxy for SLCO transporter/AKR1C3 expression), and cell–cell adhesion strength (as a proxy for EMT state, with adhesion and motility varied reciprocally). Details about parameter choices and agent-based model setup are described in Supplementary Section S7. All simulations were run on the Bridges-2 cluster at the Pittsburgh Supercomputing Center; 10 stochastic replicates were conducted per condition and results are reported as mean ± standard deviation.

### Clustering Index for quantifying clustering in the prostate cancer tissue-level ABM simulations

To quantify emergent spatial organization of resistant clones, we defined a normalized clustering index C(t) based on the fraction of homotypic neighbors for each agent:

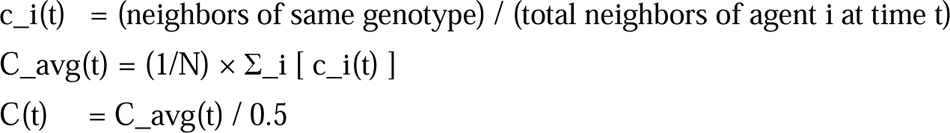

A value of C(t) > 1 indicates progressive spatial segregation of resistant clones relative to the initial random distribution; C(t) ≈ 1 indicates maintained mixing.

### Machine Learning Surrogate and SHAP Feature Ranking

The computational cost of global sensitivity analysis directly on the MHS model, which integrates ODEs and Boolean logic across disparate timescales, prohibits traditional variance-based methods. A machine learning surrogate was therefore trained to approximate MHS model behavior across the full parameter space, enabling efficient feature ranking.

Training data were generated from MHS simulations across all three cohorts, with NCG probability as the output. Cell fate was binarized as proliferative (NCG ≥ 0.15) or quiescent (NCG < 0.15); the threshold of 0.15 was selected by training classifiers across thresholds of 0 - 0.45 (increments of 0.05) and identifying the value that maximized balanced accuracy across all classifiers (Supplementary Section S5). Three classifier architectures were trained: Random Forest (RF), Support Vector Machine (SVM), and Neural Network (NN). Input features were z-score normalized; class imbalance was corrected using SMOTE; and hyperparameters were optimized by 5-fold cross-validation. Classifier performance was evaluated on held-out test sets by balanced accuracy.

Shapley Additive Explanations (SHAP) ^79^ were applied to each trained surrogate to rank model species by their marginal contribution to cell fate predictions. Features identified as top-ranked across all three classifier architectures were treated as robust consensus drivers, minimizing dependence on any single model’s inductive bias. Analyses were implemented in Python v3.8.5; computationally intensive steps (hyperparameter tuning, SHAP) were run on the Bridges-2 cluster.

### Kaplan - Meier Survival Analysis and Cox Proportional Hazards Regression

To evaluate the clinical relevance of the molecular features identified by SHAP analysis, Kaplan-Meier (KM) survival analysis was performed on prostate cancer patient data from cBioPortal (cbioportal.org) via the public REST API. Two independent analytical frameworks were applied: a combined molecular alteration analysis across seven OS-eligible cohorts, and a per-cohort mRNA expression analysis across four cohorts with available transcriptomic data. KM curves and log-rank tests were computed using the lifelines library (v0.30.3); Cox Proportional Hazards models were fit using lifelines CoxPHFitter with cohort code and age (where available) as covariates in the combined molecular analyses. A minimum of 20 patients per KM arm and 10 events per Cox model were required. Statistical significance was assessed at p < 0.05.

### Genomic Alteration-Based Stratification Approaches and Cohorts

Seven independent cohorts with OS data were downloaded and harmonized (Table 1). Overall survival (OS) was used as the primary endpoint throughout. A patient was classified as altered if they harbored any of the following event types, combined via logical OR: (1) somatic point mutations, restricted to truncating variants (nonsense, frameshift insertion or deletion, splice site, splice region, translation start site loss, or nonstop mutations) for tumor suppressors, and any recorded somatic mutation for oncogenes; (2) copy number alterations (CNA) derived from GISTIC scores, using a strict threshold of GISTIC score ≥ +2 for oncogene amplification and ≤ −2 for homozygous deletion of tumor suppressors; and (3) structural variants (SVs), including gene fusions, translocations, and rearrangements, where available in the source cohort. A patient was classified as wildtype only if confirmed as sequenced at that locus with no alteration detected across any class. Unsequenced patients were assigned NaN and excluded from analysis. Genes with fewer than 20 patients in the altered arm across the combined cohort were excluded from KM analysis. (Details in Supplementary Section S6).

**Table 1.** Patient Cohorts Used for Molecular KM Analysis.

| Cohort | Disease Type | cBioPortal N (# Patients) | N with OS Data | OS Events |
| --- | --- | --- | --- | --- |
| TCGA-PRAD | Primary | 494 | 494 | 10 |
| SU2C/PCF CRPC (2019) | Metastatic CRPC | 444 | 128 | 76 |
| MCTP Michigan | Metastatic | 119 | 48 | 48 |
| MSK Prostate (2025) | Mixed | 120 | 114 | 103 |
| MSK PIK3R1 (2021) | Mixed | 1,417 | 1,314 | 433 |
| MSK Clin Cancer Res (2024) | Mixed | 2,260 | 2,197 | 916 |
| Metastatic CSPC MSK | Metastatic CSPC | 424 | 423 | 102 |
| Combined |  | 5,278 | 4,718 | 1,688 |

### mRNA Expression-Based Stratification Approaches and Cohorts

For the expression-based analysis, four cohorts (Table 2) with available mRNA z-score profiles were analyzed on a per-cohort basis, with the survival endpoint matched to the events available in each cohort. Disease-free survival (DFS) was used for TCGA-PRAD and MSKCC, given insufficient OS events in localized disease. Overall survival (OS) was used for the metastatic SU2C and MCTP cohorts. Progression-free survival (PFS) was additionally analyzed for TCGA-PRAD, which had sufficient logged PFS events. MCTP contributed Cox hazard ratio estimates only and was excluded from Kaplan-Meier stratification due to small cohort size.

**Table 2.**
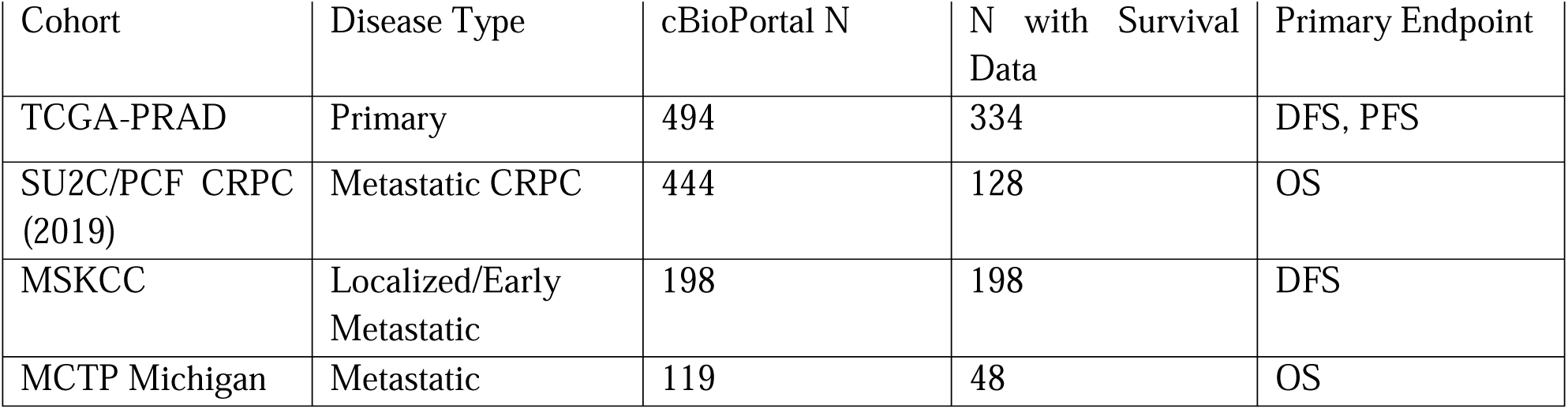
Patient Cohorts Used for mRNA Expression KM Analysis.

Continuous mRNA z-scores were dichotomized into High and Low groups using one of two strategies, selected per analysis. A z-score threshold (z > 1.0 versus the rest) was applied for the androgen uptake/synthesis and adhesion-motility axes. A median split (above versus below the cohort median) was applied for the crowding/mechanotransduction axis, with ANXA1 instead stratified by a top-versus-bottom quartile split for robustness. The stratification strategy used for each gene is indicated in the corresponding supplementary tables. (Details in Supplementary Section S6).

### Kaplan-Meier Survival Curves

Kaplan-Meier survival curves were constructed separately for the altered and wildtype groups for each gene. Survival probability was estimated at each event time using the product-limit estimator. Median survival was defined as the time at which the survival function crossed 0.50; where the curve did not reach 50% by the end of follow-up, the median was reported as not reached or derived from curve extrapolation. Statistical comparison between the two groups was performed using the log-rank (Mantel-Cox) test, with a two-sided significance threshold of p < 0.05.

### Multivariate Cox Proportional Hazards Regression

KM analysis, while informative, compares survival between groups without accounting for confounding variables. To determine whether gene-level alterations are independently associated with overall survival beyond the influence of patient age and study cohort of origin, multivariate Cox proportional hazards regression was performed for each gene whose alteration met the minimum statistical threshold of 10 events per arm. The Cox model estimates the hazard ratio (HR), which quantifies the instantaneous risk of death in the altered group relative to the unaltered group at any given time point, adjusted simultaneously for the effects of covariates. A HR greater than 1 indicates that patients with the alteration experience a higher rate of mortality, while a HR less than 1 indicates a protective association.

The binary alteration flag (altered vs. wildtype) was entered as the primary predictor. Covariates included study cohort (encoded as integer codes) and patient age at diagnosis (wherever available). Hazard ratios (HR) and 95% confidence intervals (CI) are reported from the multivariate model.

## Data and Code Availability

All TCGA clinical and genomic data used in this study are publicly available via the GDC Data Portal (https://portal.gdc.cancer.gov) and cBioPortal (https://www.cbioportal.org). Differential gene expression analysis was performed using the R2 Genomics platform (http://r2.amc.nl). The EUREKA-1 dataset is not directly available but we include DEG lists from anonymized patients with this manuscript. MHS model files (COPASI format), ABM PhysiCell configuration files, and Python analysis scripts are provided in the Supplementary Materials and will be deposited to https://github.com/Sharvari303/prostate-cancer-multiscale-model upon acceptance of the manuscript.

## Acknowledgements

We thank members of the Radhakrishnan and Janmey Labs for insightful discussions related to the manuscript. We also thank Georgios Stamatakos, Nobert Graf, Ilaria Stura, and Caterina Guiot for discussions on the early adaptations of the molecular model pursued under the Computational Horizons in Cancer (CHIC) EU funding initiative. This work has been supported in part by the National Institutes of Health under grants from NIGMS and NCI. Computational Resources were made available in part from the Advanced Cyberinfrastructure Coordination Ecosystem: Services & Support. (ACCESS) under grant MCB200101 and through the Penn Advanced Research Computing Center’s Betty cluster.

## Author contributions

S.K. and M.T. contributed equally to this work. S.K., M.T., A.G., and A.R. contributed to conceptualization, methodology, software development, formal analysis, investigation, data curation, and writing of the original draft. N.G. and G.S. contributed to conceptualization and writing (review and editing). I.S. and C.G. contributed to methodology development of the cellular signaling model, data curation and validation of the clinical datasets, and to writing (review and editing). R.R. (corresponding author) contributed to conceptualization, supervision, project administration, funding acquisition, and writing (review and editing). All authors reviewed and approved the final manuscript.

## References

1 Da, S., Me, O. N., Tb, R., Nf, D. & Hk, W. Prostate Cancer Incidence and Survival, by Stage and Race/Ethnicity - United States, 2001-2017 - PubMed. MMWR. Morbidity and mortality weekly report 69 (10/16/2020). 10.15585/mmwr.mm6941a1

2 TK, W. & OT, Z. Prostate Cancer: A Review of Genetics, Current Biomarkers and Personalised Treatments - PubMed. Cancer reports (Hoboken, N.J.) 7 (2024 Oct). 10.1002/cnr2.70016

3 T, B. & P, P. Prediction of biochemical recurrence after laparoscopic radical prostatectomy - PubMed. BMC urology 23 (11/13/2023). 10.1186/s12894-023-01350-2

4 Pound, C. R. et al. Natural History of Progression After PSA Elevation Following Radical Prostatectomy. JAMA 281 (1999/05/05). 10.1001/jama.281.17.1591

5 EM, S., et al. NCCN Guidelines® Insights: Prostate Cancer, Version 3.2024 - PubMed. Journal of the National Comprehensive Cancer Network: JNCCN 22 (2024 Apr). 10.6004/jnccn.2024.0019

6 Epstein, J. I. et al. The 2014 International Society of Urological Pathology (ISUP) Consensus Conference on Gleason Grading of Prostatic Carcinoma: Definition of Grading Patterns and Proposal for a New Grading System. The American Journal of Surgical Pathology 40 (February 2016). 10.1097/PAS.0000000000000530

7 DV, M., et al. Updated nomogram to predict pathologic stage of prostate cancer given prostate-specific antigen level, clinical stage, and biopsy Gleason score (Partin tables) based on cases from 2000 to 2005 - PubMed. Urology 69 (2007 Jun). 10.1016/j.urology.2007.03.042

8 MR, C., et al. The University of California, San Francisco Cancer of the Prostate Risk Assessment score: a straightforward and reliable preoperative predictor of disease recurrence after radical prostatectomy - PubMed. The Journal of urology 173 (2005 Jun). 10.1097/01.ju.0000158155.33890.e7

9 Sandeman, K. et al. Prostate MRI added to CAPRA, MSKCC and Partin cancer nomograms significantly enhances the prediction of adverse findings and biochemical recurrence after radical prostatectomy. PLOS ONE 15 (9 Jul 2020). 10.1371/journal.pone.0235779

10 M, S., et al. Androgen receptor mutations for precision medicine in prostate cancer - PubMed. Endocrine-related cancer 29 (08/17/2022). 10.1530/ERC-22-0140

11 ED, C., et al. Androgen-targeted therapy in men with prostate cancer: evolving practice and future considerations - PubMed. Prostate cancer and prostatic diseases 22 (2019 Mar). 10.1038/s41391-018-0079-0

12 Antonarakis, E. S. et al. AR-V7 and Resistance to Enzalutamide and Abiraterone in Prostate Cancer. New England Journal of Medicine 371 (2014-09-11). 10.1056/NEJMoa1315815

13 T, K., PG, C. & TC, T. Prostate cancer progression after androgen deprivation therapy: mechanisms of castrate resistance and novel therapeutic approaches - PubMed. Oncogene 32 (12/05/2013). 10.1038/onc.2013.206

14 Cairns, P. et al. Frequent inactivation of PTEN/MMAC1 in primary prostate cancer. Cancer research 57, 4997–5000 (1997).

15 BS, C., et al. Reciprocal feedback regulation of PI3K and androgen receptor signaling in PTEN-deficient prostate cancer - PubMed. Cancer cell 19 (05/17/2011). 10.1016/j.ccr.2011.04.008

16 Ogawara, Y. et al. Akt Enhances Mdm2-mediated Ubiquitination and Degradation of p53. Journal of Biological Chemistry 277 (2002/06/14). 10.1074/jbc.M109745200

17 Yang, M. et al. SLCO2B1 and SLCO1B3 May Determine Time to Progression for Patients Receiving Androgen Deprivation Therapy for Prostate Cancer. Journal of Clinical Oncology 29 (2011-6-20). 10.1200/JCO.2010.31.2405

18 C, C., et al. Intratumoral de novo steroid synthesis activates androgen receptor in castration-resistant prostate cancer and is upregulated by treatment with CYP17A1 inhibitors - PubMed. Cancer research 71 (10/15/2011). 10.1158/0008-5472.CAN-11-0532

19 Ea, M., et al. Contribution of Adrenal Glands to Intratumor Androgens and Growth of Castration-Resistant Prostate Cancer - PubMed. Clinical cancer research: an official journal of the American Association for Cancer Research 25 (01/01/2019). 10.1158/1078-0432.CCR-18-1431

20 Thiery, J. P., Acloque, H., Huang, R. Y. J. & Nieto, M. A. Epithelial-Mesenchymal Transitions in Development and Disease. Cell 139 (2009/11/25). 10.1016/j.cell.2009.11.007

21 Jolly, M. K. et al. Implications of the Hybrid Epithelial/Mesenchymal Phenotype in Metastasis. Frontiers in Oncology 5 (2015 Jul 20). 10.3389/fonc.2015.00155

22 NP, G., A, W., EW, T. & HJ, H. Mesenchymal-epithelial transition (MET) as a mechanism for metastatic colonisation in breast cancer - PubMed. Cancer metastasis reviews 31 (2012 Dec). 10.1007/s10555-012-9377-5

23 Kolokotroni, E. et al. A Multidisciplinary Hyper-Modeling Scheme in Personalized In Silico Oncology: Coupling Cell Kinetics with Metabolism, Signaling Networks, and Biomechanics as Plug-In Component Models of a Cancer Digital Twin. Journal of Personalized Medicine 14 (2024 Apr 29). 10.3390/jpm14050475

24 Hashemi, M. et al. Targeting PI3K/Akt signaling in prostate cancer therapy. Journal of Cell Communication and Signaling 17 (2022 Nov 11). 10.1007/s12079-022-00702-1

25 Lee, J. T. et al. Targeting prostate cancer based on signal transduction and cell cycle pathways. Cell cycle (Georgetown, Tex.) 7 (2008 Jun 16). 10.4161/cc.7.12.6166

26 B, W., et al. A spatial model predicts that dispersal and cell turnover limit intratumour heterogeneity - PubMed. Nature 525 (09/10/2015). 10.1038/nature14971

27 AR, A. & V, Q. Integrative mathematical oncology - PubMed. Nature reviews. Cancer 8 (2008 Mar). 10.1038/nrc2329

28. Data collection for models validation: Application to prostate cancer — Clinical aspects. IEEE-EMBS International Conference on Biomedical and Health Informatics (BHI) (2014). 10.1109/BHI.2014.6864290

29 CG, C. & SD, F. Multi-scale modeling of macrophage-T cell interactions within the tumor microenvironment - PubMed. PLoS computational biology 16 (12/23/2020). 10.1371/journal.pcbi.1008519

30 Alsinnawi, M. et al. Association of prostate cancer SLCO gene expression with Gleason grade and alterations following androgen deprivation therapy. Prostate Cancer and Prostatic Diseases 2019 22:4 22 (2019-03-19). 10.1038/s41391-019-0141-6

31 Umbas, R. et al. Decreased E-Cadherin Expression Is Associated with Poor Prognosis in Patients with Prostate Cancer. Cancer Research 54, 3929–3933 (1994).

32 Henderson, V. et al. Snail promotes cell migration through PI3K/AKT-dependent Rac1 activation as well as PI3K/AKT-independent pathways during prostate cancer progression. Cell Adhesion & Migration 9 (2015 Jul 24). 10.1080/19336918.2015.1013383

33 Voss, G. et al. Regulation of cell–cell adhesion in prostate cancer cells by microRNA-96 through upregulation of E-Cadherin and EpCAM. Carcinogenesis 41 (2019 Nov 18). 10.1093/carcin/bgz191

34 Poma, A. M. et al. Hippo pathway affects survival of cancer patients: extensive analysis of TCGA data and review of literature. Scientific Reports 2018 8:1 8 (2018-07-13). 10.1038/s41598-018-28928-3

35 H, X., Z, C. & C, L. The prognostic value of Piezo1 in breast cancer patients with various clinicopathological features - PubMed. Anti-cancer drugs 32 (04/01/2021). 10.1097/CAD.0000000000001049

36 Poli, A. et al. PIP4K2B is mechanoresponsive and controls heterochromatin-driven nuclear softening through UHRF1. Nature Communications 2023 14:1 14 (2023-03-14). 10.1038/s41467-023-37064-0

37 Qie, S. & Diehl, J. A. Cyclin D1, Cancer Progression and Opportunities in Cancer Treatment. *Journal of molecular medicine (Berlin*, Germany) 94 (2016 Oct 2). 10.1007/s00109-016-1475-3

38 Oaa, M., et al. The role of hypoxia on prostate cancer progression and metastasis - PubMed. Molecular biology reports 50 (2023 Apr). 10.1007/s11033-023-08251-5

39. Nukpezah, J. Unlocking Cancer Insights: A Data-Driven Approach to Unravel Genomic Variations in Kinome and Genomic and Proteome Remodeling by Mechano-Chemical Signals Doctor of Philosophy thesis, University of Pennsylvania, (2025).

40 B, Z., et al. Inactivation of YAP oncoprotein by the Hippo pathway is involved in cell contact inhibition and tissue growth control - PubMed. Genes & development 21 (11/01/2007). 10.1101/gad.1602907

41 S, D. Regulation of YAP/TAZ Activity by Mechanical Cues: An Experimental Overview - PubMed. Methods in molecular biology (Clifton, N.J.) 1893 (2019). 10.1007/978-1-4939-8910-2_15

42 Ju, X. et al. Identification of a Cyclin D1 Network in Prostate Cancer That Antagonizes Epithelial–Mesenchymal Restraint. Cancer Research 74 (2014/01/15). 10.1158/0008-5472.CAN-13-1313

43 H Zhong, F. A., A A Baccala, E Laughner, N Rioseco-Camacho, W B Isaacs, J W Simons, G L Semenza. Increased expression of hypoxia inducible factor-1alpha in rat and human prostate cancer. Cancer Res 58(**23**):5280-4. (1998).

44 Miyazaki, Y. et al. Consecutive Prostate Cancer Specimens Revealed Increased Aldo–Keto Reductase Family 1 Member C3 Expression with Progression to Castration-Resistant Prostate Cancer. Journal of Clinical Medicine 8 (2019 May 1). 10.3390/jcm8050601

45 K, G., OJ, H., SA, H. & LA, A. A switch from E-cadherin to N-cadherin expression indicates epithelial to mesenchymal transition and is of strong and independent importance for the progress of prostate cancer - PubMed. Clinical cancer research: an official journal of the American Association for Cancer Research 13 (12/01/2007). 10.1158/1078-0432.CCR-07-1263

46 Wen, Y.-C. et al. Snail as a potential marker for predicting the recurrence of prostate cancer in patients at stage T2 after radical prostatectomy. Clinica Chimica Acta 431 (2014/04/20). 10.1016/j.cca.2014.01.036

47 Barber, A. G. et al. PI3K/AKT pathway regulates E cadherin and Desmoglein 2 in aggressive prostate cancer. Cancer Medicine 4 (2015/08/01). 10.1002/cam4.463

48 Schmidt, L. J. et al. RhoA as a Mediator of Clinically Relevant Androgen Action in Prostate Cancer Cells. Molecular Endocrinology 26 (2012 Mar 28). 10.1210/me.2011-1130

49 Phan, T. et al. Review: Mathematical Modeling of Prostate Cancer and Clinical Application. Applied Sciences 2020, Vol. 10, Page 2721 **10** (2020-04-15). 10.3390/app10082721

50 Mejía-Hernández, J. O. et al. Targeting MDM4 as a Novel Therapeutic Approach in Prostate Cancer Independent of p53 Status. Cancers 14 (2022 Aug 16). 10.3390/cancers14163947

51 Liu, J. et al. MDM4 was associated with poor prognosis and tumor-immune infiltration of cancers. European Journal of Medical Research 29 (2024 Jan 27). 10.1186/s40001-024-01684-z

52 Gerhart, S. V. et al. Activation of the p53-MDM4 regulatory axis defines the anti-tumour response to PRMT5 inhibition through its role in regulating cellular splicing. Scientific Reports 2018 8:1 8 (2018-06-26). 10.1038/s41598-018-28002-y

53 M, P., S, K., M, K., M, G. & M, V. Multi-scale tissue architecture analysis of favorable-risk prostate cancer: Correlation with biochemical recurrence - PubMed. Investigative and clinical urology 61 (2020 Sep). 10.4111/icu.20200018

54 M, B., S, P., F, D. & MA, R. TMPRSS2-ERG fusion heterogeneity in multifocal prostate cancer: clinical and biologic implications - PubMed. Urology 70 (2007 Oct). 10.1016/j.urology.2007.08.032

55 MC, H., et al. Genomic and phenotypic heterogeneity in prostate cancer - PubMed. Nature reviews. Urology 18 (2021 Feb). 10.1038/s41585-020-00400-w

56 Hong, M. K. H. et al. Tracking the origins and drivers of subclonal metastatic expansion in prostate cancer. Nature Communications 2015 6:1 6 (2015-04-01). 10.1038/ncomms7605

57 D, B., et al. Intratumor DNA methylation heterogeneity reflects clonal evolution in aggressive prostate cancer - PubMed. Cell reports 8 (08/07/2014). 10.1016/j.celrep.2014.06.053

58 G, C., K, G. & N, K. Epigenetic mechanisms underlying subtype heterogeneity and tumor recurrence in prostate cancer - PubMed. Nature communications 14 (02/02/2023). 10.1038/s41467-023-36253-1

59 A, H., et al. Effect of SLCO1B3 haplotype on testosterone transport and clinical outcome in caucasian patients with androgen-independent prostatic cancer - PubMed. Clinical cancer research: an official journal of the American Association for Cancer Research 14 (06/01/2008). 10.1158/1078-0432.CCR-07-4118

60 VK, A., et al. Glucocorticoid receptor confers resistance to antiandrogens by bypassing androgen receptor blockade - PubMed. Cell 155 (12/05/2013). 10.1016/j.cell.2013.11.012

61 Heckmann, N. et al. High level expression of glucocorticoid receptor (GR) is linked to aggressive tumor features, early biochemical recurrence, and genetic instability in prostate cancer. Prostate Cancer and Prostatic Diseases 2025 29:1 29 (2025-11-05). 10.1038/s41391-025-01046-8

62 Pan, J. et al. Frontiers | Identification of cancer-associated fibroblasts subtypes in prostate cancer. Frontiers in Immunology 14 (2023/03/24). 10.3389/fimmu.2023.1133160

63 H, W., et al. Antiandrogen treatment induces stromal cell reprogramming to promote castration resistance in prostate cancer - PubMed. Cancer cell 41 (07/10/2023). 10.1016/j.ccell.2023.05.016

64 B, C., et al. Loss of androgen receptor signaling in prostate cancer-associated fibroblasts (CAFs) promotes CCL2- and CXCL8-mediated cancer cell migration - PubMed. Molecular oncology 12 (2018 Aug). 10.1002/1878-0261.12327

65 C, L., T, H., DP, L. & F, B. The Extracellular Matrix Stiffening: A Trigger of Prostate Cancer Progression and Castration Resistance? - PubMed. Cancers 14 (06/11/2022). 10.3390/cancers14122887

66 Brea, L. & Yu, J. Tumor-intrinsic regulators of the immune-cold microenvironment of prostate cancer. Trends in Endocrinology & Metabolism 36 (2025/09/01). 10.1016/j.tem.2024.12.003

67 Anton, A. et al. An immune suppressive tumor microenvironment in primary prostate cancer promotes tumor immune escape. PLOS ONE 19 (27 Nov 2024). 10.1371/journal.pone.0301943

68 MC, D., et al. Androgen Deprivation Therapy Drives a Distinct Immune Phenotype in Localized Prostate Cancer - PubMed. Clinical cancer research: an official journal of the American Association for Cancer Research 30 (11/15/2024). 10.1158/1078-0432.CCR-24-0060

69 A, J., et al. The temporal dynamics of the immune response to neoadjuvant androgen deprivation therapy suggests a window-of-opportunity for checkpoint inhibitor therapy in prostate cancer - PubMed. medRxiv: the preprint server for health sciences (01/13/2026). 10.64898/2026.01.10.26343859

70 van Genderen, M. N. G. et al. Agent-based modeling of the prostate tumor microenvironment uncovers spatial tumor growth constraints and immunomodulatory properties. npj Systems Biology and Applications 2024 10:1 10 (2024-02-21). 10.1038/s41540-024-00344-6

71 Ji, Z., Zhao, W., Lin, H.-K. & Zhou, X. Systematically understanding the immunity leading to CRPC progression. PLOS Computational Biology 15 (10 Sept 2019). 10.1371/journal.pcbi.1007344

72 Ali, A. et al. Single-cell and spatial RNA sequencing in prostate cancer. Nature Reviews Urology 2026 (2026-05-28). 10.1038/s41585-026-01149-4

73 He, Y. et al. Targeting signaling pathways in prostate cancer: mechanisms and clinical trials. Signal Transduction and Targeted Therapy 2022 7:1 7 (2022-06-24). 10.1038/s41392-022-01042-7

74 C, H. & CV, H. Studies on prostatic cancer. I. The effect of castration, of estrogen and androgen injection on serum phosphatases in metastatic carcinoma of the prostate - PubMed. CA: a cancer journal for clinicians 22 (1972 Jul-Aug). 10.3322/canjclin.22.4.232

75 Hoops, S. et al. COPASI—a COmplex PAthway SImulator. Bioinformatics 22 (2006/12/15). 10.1093/bioinformatics/btl485

76 JD, M., A, P., RA, E., JD, N. & Y, K. Mechanisms of resistance to intermittent androgen deprivation in patients with prostate cancer identified by a novel computational method - PubMed. Cancer research 74 (07/15/2014). 10.1158/0008-5472.CAN-13-3162

77 A, G., R, H., SH, F., SM, M. & P, M. PhysiCell: An open source physics-based cell simulator for 3-D multicellular systems - PubMed. PLoS computational biology 14 (02/23/2018). 10.1371/journal.pcbi.1005991

78 Ghaffarizadeh, A., Friedman, S. H. & Macklin, P. BioFVM: an efficient, parallelized diffusive transport solver for 3-D biological simulations. Bioinformatics 32 (2015 Dec 12). 10.1093/bioinformatics/btv730

79 Salih, A. et al. A Perspective on Explainable Artificial Intelligence Methods: SHAP and LIME. (2023/05/03). 10.48550/arXiv.2305.02012

