## Supplementary material for "Coupled Cell-Intrinsic and Microenvironmental Heterogeneity Drives Divergent Trajectories in Castration-Resistant Prostate Cancer": SI Text and Figures

**Supplementary Information**: **MHS Model Description and Validation**

### S1. Molecular Submodel Architecture

The Multiscale Hybrid Signaling (MHS) model integrates three molecular sub models operating at distinct biological timescales: androgen receptor (AR) signaling, growth factor (GF)-mediated Ras/MAPK and PI3K/AKT signaling, and p53-mediated cell cycle.^1^ Together, these sub models encode the intracellular logic that determines whether a cell proliferates, undergoes apoptosis, or enters growth arrest - the cell-fate outcomes whose cohort-specific distributions drive the divergent recurrence trajectories described in the main text.

#### S1.1 AR Signaling Sub model

The AR signaling sub model captures testosterone-driven transcriptional activation and PSA production across three compartments: extracellular medium, cytoplasm, and nucleus. (Figure S1.1.1) Extracellular testosterone is maintained at a buffered physiological level and enters the cell at a fixed rate. In the cytoplasm, testosterone is converted to dihydrotestosterone (DHT) by 5α-reductase; both ligands bind the androgen receptor, with DHT binding at substantially higher affinity.^2^ Ligand binding triggers release of heat-shock proteins, AR dimerization, and nuclear translocation. Nuclear AR dimers bind DNA and regulate transcription of target genes including PSA, modeled as a catalytic reaction with AR dimer as the activator. Separate reactions govern AR synthesis and degradation of both AR and PSA. The complete reaction set is given in Table S1.1.1.

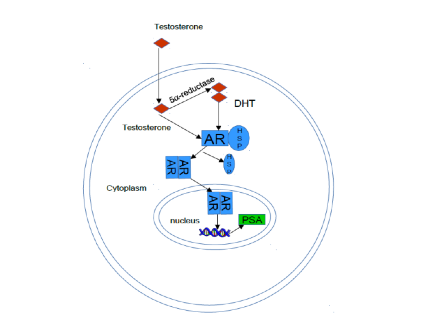

Figure S1.1.1 Schematic diagram of AR activation by testosterone/DHT and its translocation to the nucleus and subsequent transcription and release of PSA into the serum.

AR can also be activated by non-androgenic signals, in particular GF mediated (e.g., ErbB family receptors, enabling continued AR pathway activity at low androgen concentrations and providing a key mechanism for ligand-independent CRPC progression. ^3^ Additionally, in cell cycle regulation, p53 suppresses AR synthesis, such that p53 inactivation (common in aggressive disease) elevates AR expression and promotes cell cycle entry ^4-6^. AR in turn upregulates cell cycle promoters (CDK4) and downregulates inhibitors (p21) ^7^.

**Table S1.1.1.** *Reactions in the AR signaling submodel.*

| **No.** | **Reaction** | **Equation** |
| --- | --- | --- |
| 1 | AR dimerization (testosterone) | AR(cyt) + AR(cyt) → AR₂; T |
| 2 | AR dimerization (DHT) | AR(cyt) + AR(cyt) → AR₂(cyt); D |
| 3 | AR nuclear transport | AR₂(cyt) → AR₂(nuc) |
| 4 | AR synthesis | → AR(cyt) |
| 5 | AR degradation | AR(cyt) → |
| 6 | PSA synthesis (AR-catalyzed) | AR₂(nuc) → AR₂(nuc) + PSA |
| 7 | PSA synthesis (basal) | → PSA |
| 8 | PSA degradation | PSA → |
| 9 | Non-androgenic AR activation | AR(cyt) + AR(cyt) → AR₂; N |
| 10 | Testosterone reduction | T → D; 5α-reductase |

#### S1.2 GF-Mediated Ras/MAPK and PI3K/AKT Signaling Submodel

The Ras/MAPK and PI3K/AKT submodel captures mitogenic and survival signaling downstream of a mitogenic growth factor receptor. Cross-talk between these two pathways was adopted from published work by Chen and colleagues ^8^. PTEN negatively regulates the PI3K/AKT arm; PTEN-deleted cells exhibit constitutive AKT activation and reciprocal feedback between PI3K/AKT and AR signaling, such that pharmacological inhibition of one arm activates the other - a well-documented mechanism underlying therapeutic resistance in CRPC. Steady-state outputs of this sub model, including phospho-AKT and phospho-ERK, are passed to the AR signaling sub model as described in Supplementary section S2.

#### S1.3 Cell cycle and p53-Mediated DNA Damage Response Submodel

The p53 module was adopted from a published discrete Boolean model ^9^, in which each species takes a binary state (0 or 1) and the network generates three distinct cell fate outcomes: proliferation, apoptosis, and growth arrest (senescence). Because p53-governed processes operate on timescales substantially longer than AR or GF signaling ^9^, the module receives discretized inputs from the AR sub model and returns binary state values for downstream species (including CDK4, p21) that in turn influence AR-driven transcription. The interfacing logic between modules is described in Supplementary section S2.

#### S1.4 Two-Population Tumor Growth Sub model

Two mechanisms have been proposed to explain CRPC onset: an adaptation model, in which androgen resistance arises through acquired genetic or epigenetic events; and a selection model, in which resistance reflects the preferential survival and expansion of pre-existing resistant clones ^10^. Clinical evidence from PTEN immunostaining supports the co-existence of two distinct cell populations - homogeneous and heterogeneous PTEN loss - within the same CRPC tumor. ^11^ The MHS model adopts the selection framework, initializing tumors with a mixed population of androgen-sensitive (S) and androgen-resistant (R) cells whose intrinsic growth potentials are defined by the upstream molecular sub models.

Net cell growth (NCG) - the difference between simulated cell growth and cell kill probabilities - was estimated by systematically varying three key inputs: endogenous GF concentration (0.1 nM and 100 nM ^12^, testosterone concentration (0.05–20 nM, spanning the range of published PCa patient PK data ^13^), and PTEN status (normal, excess, deletion). The resulting NCG dynamics were parameterized as Hill functions of testosterone concentration (Figure S1.1.4), independently for S and R cell populations:

*rcond × ( 1 + mcond × Tncond / ( pcondncond + Tncond* ) )

where cond denotes cell type (S or R); r, m, p, and n are parameters fitted to in vitro cell growth data; and T is testosterone concentration. This formulation captures the nonlinear, saturable androgen-dependence of each population and is the basis for all cohort-specific proliferative response functions used in the main text.

Under ADT (LHRH agonist, with or without combined AR blockade), S and R population dynamics are governed by:

dS/dt = ( rs × k + as × (K − R) / K ) × S

dR/dt = ( rr × k + ar × (K − S) / K ) × R

where r denotes the Hill-function growth rate, k is a mapping factor from molecular to population scale, a is an additional growth contribution under castrate conditions, and K is the carrying capacity. Total serum PSA is then:

PSA(t) = scoeff × S(t) + rcoeff × R(t)

with per-cell PSA production coefficients drawn from published values for sensitive (0–0.29 ag/mL/cell/day) and resistant (0.015 - 0.35 ag/mL/cell/day) populations ^14^.

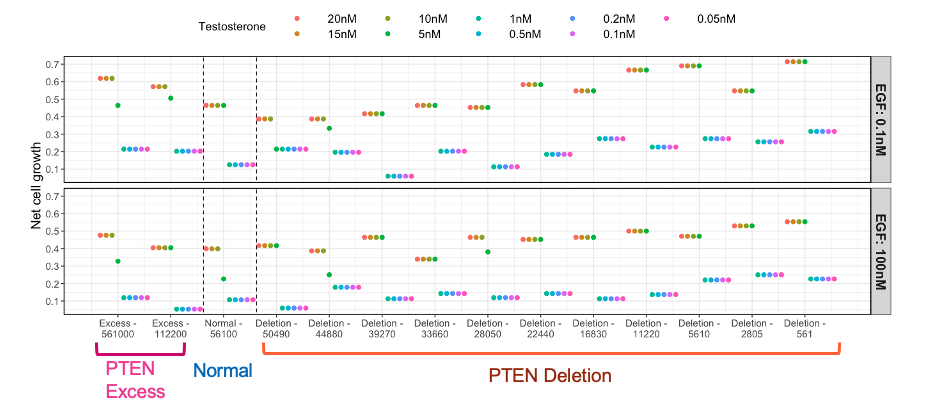

Figure S1.4.1 Tumor growth rate was based on the NCG predictions with varied GF concentration in prostate cells, PTEN level, and testosterone concentration.

### S2. Integration of Sub models Across Biological Timescales

The three sub models operate on distinct biological timescales; this hierarchy determines both the sequence of information exchange and the form of the interface between modules (Figure S2).

#### S2.1 AR signaling and GF-mediated Ras/MAPK and PI3K/AKT signaling

Both modules are formulated as systems of ordinary differential equations. Because GF-mediated signaling operates on a faster timescale than AR-driven transcription, the Ras/MAPK and PI3K/AKT module is run to steady state first, and the resulting species concentrations, including phospho-AKT (an activator of AR–testosterone binding) and phospho-ERK, are passed as fixed inputs to the AR module. This timescale separation avoids numerical stiffness while faithfully reflecting the biological order of signal propagation from membrane receptor to nuclear transcription factor.

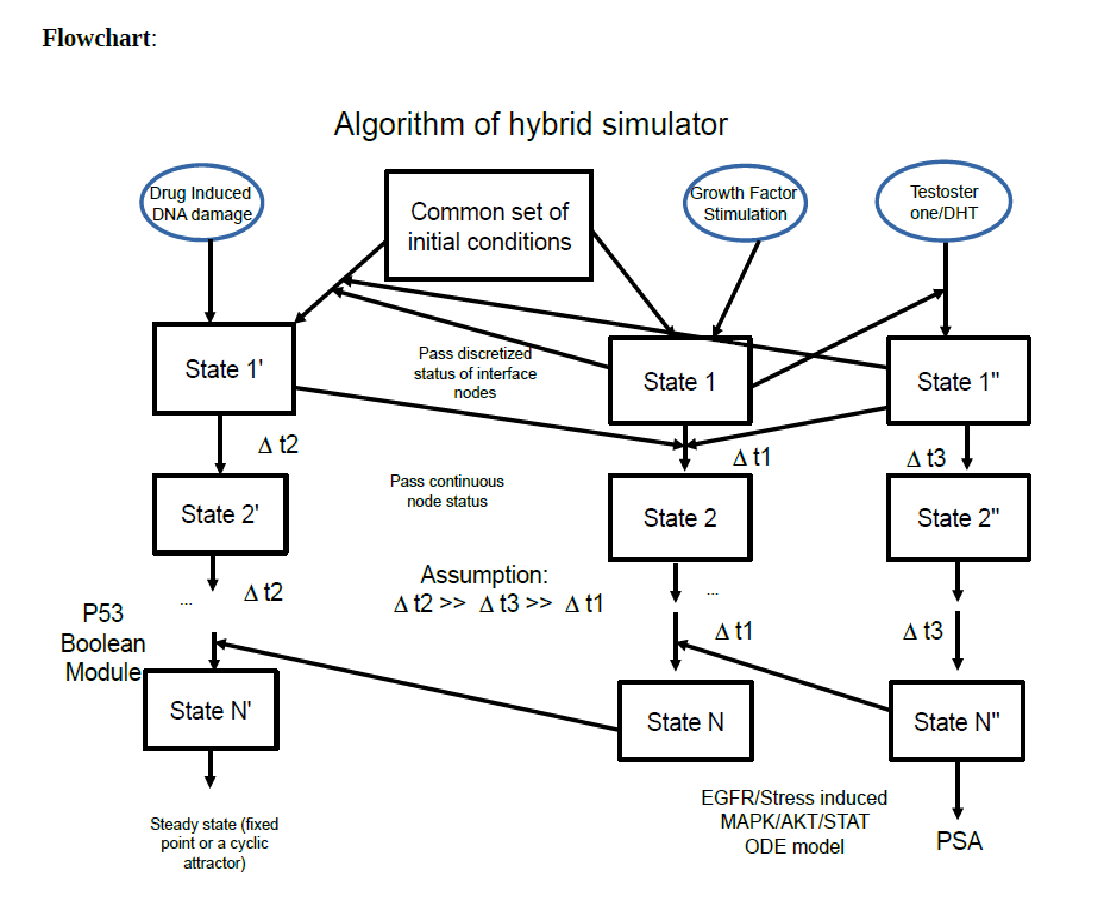

Figure S2 Three sub models with different time scale and time descriptions is integrated with the multiscale hybrid algorithm.

#### S2.2 AR signaling and p53-mediated DNA damage response

The p53 module is a discrete Boolean network in which each species takes a value of 0 or 1. Because p53-level events operate on timescales much longer than AR dynamics, continuous AR species concentrations are converted to discrete on/off signals via defined thresholds before being passed to the p53 module. Reciprocally, Boolean output states are converted to bounded continuous values for use by the AR module. Species crossing this interface include AR (influencing CDK4, p21, and Bax state transitions in the p53 module) and p53 (inhibiting AR synthesis in the ODE module). As a representative example, the net rate of AR production is:

d[AR]/dt = f( [T], [DHT], AKTact, HER2act, p53bool )

where AKT activator terms are supplied by the Ras/MAPK and PI3K/AKT module, and p53 is the Boolean inhibitor term supplied by the DNA damage module. The full three-module integration architecture, including the direction and form of all inter-module connections, is shown in Figure S2.

### S3. Cohort Parameterization from TCGA Patient Genomic Data

To translate patient genomic data into model initial conditions, TCGA prostate cancer patients were stratified into three cohorts: Control (no recurrence), Biochemical Recurrence (BR), and Tumor Recurrence (TR), based on PSA kinetics and biopsy outcomes, within both PTEN-deleted and PTEN-normal subgroups. Differentially expressed genes (DEGs) in BR and TR cohorts relative to CNT were identified using the R2 Genomics analysis platform on the TCGA dataset. The analysis method used multiple correlation testing using False Discovery Rate correction (p value 0.01). The genes that are most differentially expressed between BR vs Control in TCGA prostate adenocarcinoma dataset were CDK1, MDM4, CDKN1A, TP53, MAP4K4, EGF. The genes that are most differentially expressed between TR vs Control were CDK1, CCNE1, CDK2, RAF1, PTEN, RASGRP1. The initial concentrations of corresponding model species were adjusted to reflect these expression differences, producing cohort-specific model instances that encode the genome-wide signaling background of each patient group.

For each cohort, simulations were run across two PTEN expression levels, five testosterone concentrations (1 - 20 nM), and two intra-tumoral heterogeneity conditions representing combinations of high and low GF concentration. This yields 20 simulation instances per cohort. Conditions are summarized in Table S3.1. The resulting NCG distributions form the basis for all cohort-specific proliferative response functions used in the main text analyses.

| **(A) PTEN Expression × Testosterone Levels** | | **(B) GF Concentration** | |
| --- | --- | --- | --- |
| **PTEN Status** | **Testosterone (nM)** | **GF Condition** | **Concentration** |
| **Normal (PTEN #)**  56000 | 1 nM | **Low GF**  **High GF** | 0.1 nM  100 nM |
|  | 5 nM |  |  |
|  | 10 nM |  |  |
|  | 15 nM |  |  |
|  | 20 nM |  |  |
| **Deleted (PTEN #)**  5600 | 1 nM |  |  |
|  | 5 nM |  |  |
|  | 10 nM |  |  |
|  | 15 nM |  |  |
|  | 20 nM |  |  |

Table S3.1: Table showing the main simulation conditions. There were two main classification schemes, initial PTEN expression and Testosterone levels. We selected 5 different testosterone levels from 1nM to 20nM, two different PTEN expressions (A), and 2 GF conditions (high and low) (B).

### S4. MHS Model Sensitivity Analysis

Sensitivity analyses were performed in two stages, local followed by global, to assess model robustness and identify the molecular parameters that most strongly determine the two key model outputs: net cell growth rate (NCG) and serum PSA concentration.

#### S4.1 Local Sensitivity Analysis: Parameter Influence on NCG

Local sensitivity analysis was performed across all parameters in the three molecular sub models using Latin Hypercube Sampling (LHS). Each parameter was varied ± 20% from its nominal value and the normalized sensitivity index with respect to simulated NCG was computed. Analysis was conducted across combinations of PTEN deletion level and testosterone concentration to span the clinically relevant input space.

Results show that parameter sensitivity is broadly comparable across PTEN deletion levels at a given testosterone concentration, while testosterone level is the primary modulator of NCG sensitivity across the tested range (Figure S4.1.1). This is consistent with the biological expectation that androgen availability is the dominant environmental driver of growth rate variability in the pre-CRPC setting. The overall robustness of the model to ± 20% parameter perturbations supports the validity of the nominal parameterization for downstream cohort analyses.

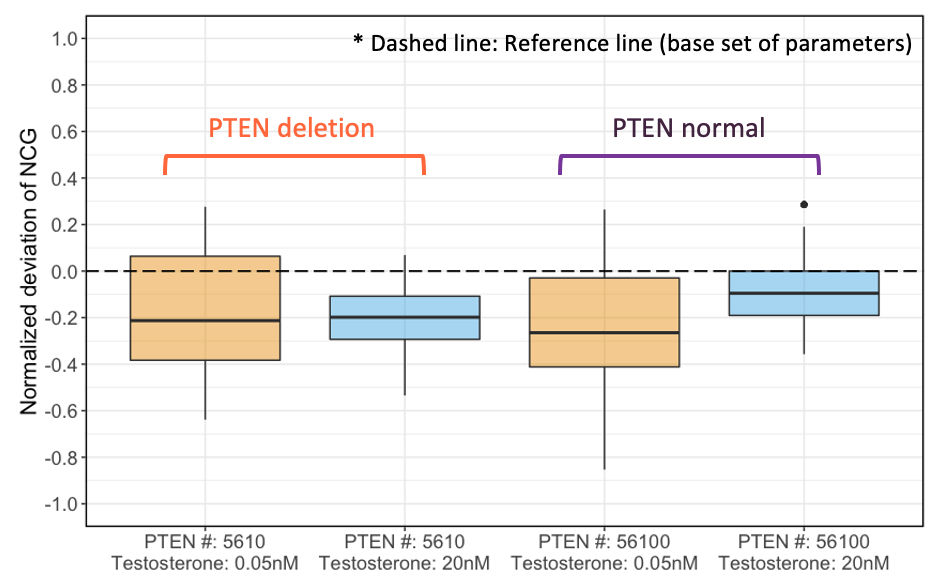

Figure S4.1.1 LHS results on sensitivity index for different tested scenarios.

#### S4.2 Global Sensitivity Analysis: Parameter Influence on PSA

Global sensitivity analysis was performed on the two-population growth sub model using the extended Fourier Amplitude Sensitivity Test (eFAST) ^15^, with simulated serum PSA as the output of interest. All parameters were simultaneously perturbed using sinusoidal sampling within biologically plausible ranges. First-order (Si) and total-order (Sti) sensitivity indices were computed for each parameter, and a two-sample t-test was applied to identify parameters with statistically significant effects on PSA dynamics.

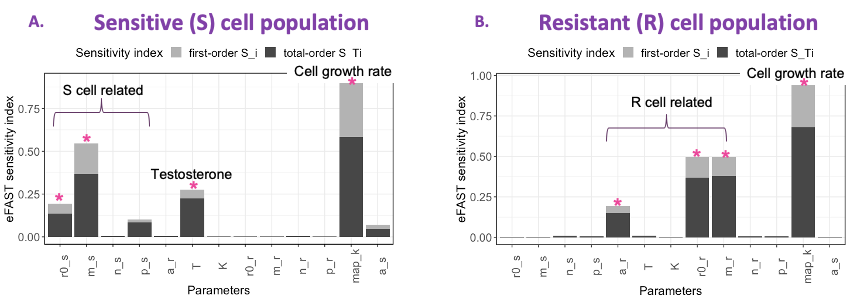

Figure S4.2.1 Sensitivity index from eFAST for BR cohort with PTEN deletion with both PSA concentration from androgen-sensitive cell population (A) and androgen-resistant cell population (B).

Analysis was carried out independently for all three cohorts (CNT, BR, TR); results for the BR cohort with PTEN deletion are shown as a representative example (Figure S4.2.1). In both the androgen-sensitive and androgen-resistant cell populations, sensitivity is concentrated within parameters intrinsic to each population's own dynamics, with limited cross-population parameter influence. This result confirms two key properties of the model: (i) the two-population sub model is structurally sufficient to describe each clone type's behavior and PSA dynamics independently; and (ii) the outputs are not dominated by parameter interactions between populations, supporting the model's ability to attribute differential PSA kinetics to biologically distinct clonal behaviors rather than to modeling crosstalk.

### S5. Validation of the Two-Population Cell Model

A foundational assumption of the MHS model is the co-existence of two functionally distinct cell populations (androgen-sensitive and androgen-resistant) with different proliferative responses to androgen availability. This assumption was validated in two complementary ways: qualitative comparison against published experimental data, and quantitative threshold identification using supervised machine learning.

#### S5.1 Qualitative Validation Against the pERK–pAKT Response Map

Chen and colleagues characterized a two-dimensional phospho-ERK (pERK) / phosphor-AKT (pAKT) response map in prostate cancer cells, in which the direction and amplitude of the ERK-AKT activity vector predicts cell fate: high pERK with low pAKT drives differentiation, low pERK with high pAKT drives proliferation, and a quiescent state occupies the low-pERK/low-pAKT region. ^16^ To test whether the MHS model reproduces this landscape, we systematically varied GF concentration and PTEN deletion level across all three patient cohorts (CNT, BR, TR) and plotted the resulting steady-state pERK and pAKT outputs. The simulated response map qualitatively reproduced the curved boundary between differentiating and proliferating states observed experimentally (Figure S5.1.1), providing a first-principles validation that the integrated sub model architecture captures the correct qualitative cell-fate logic.

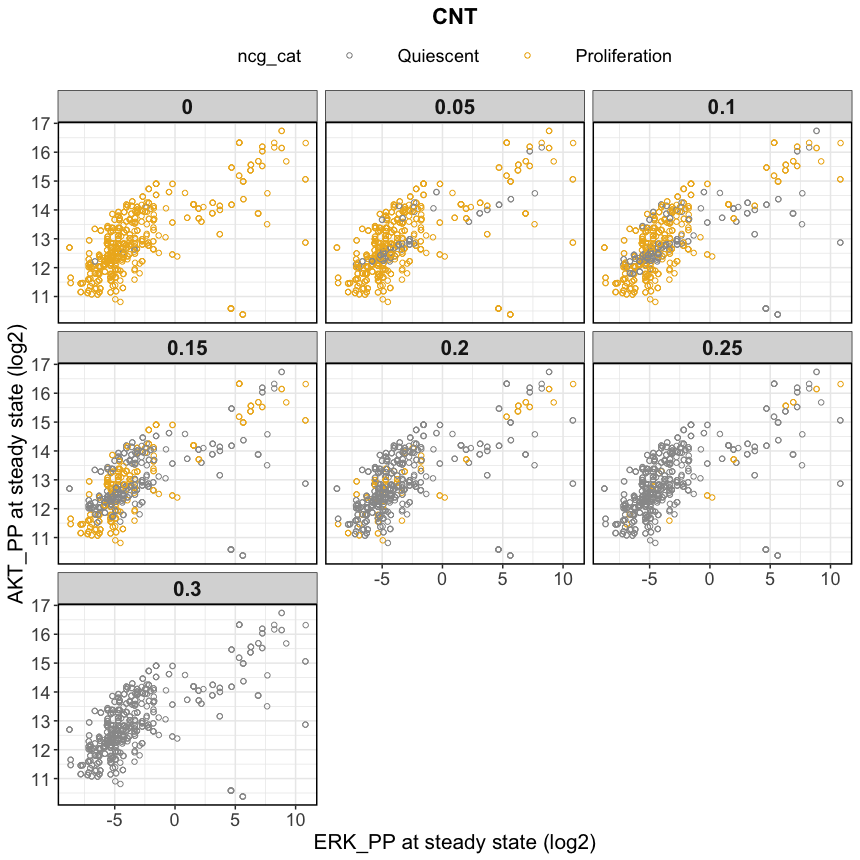

Figure S5.1.1 The simulated response map of pERK and pAKT at steady state with different NCG as threshold to assess cell behaviors.

#### S5.2 Quantitative Threshold Identification by Supervised Machine Learning

To identify the NCG value that most robustly separates the two cell populations, simulation outputs were classified at NCG thresholds ranging from 0 to 0.45 (in increments of 0.05), using three independently trained supervised classifiers: Support Vector Machine (SVM), Decision Tree, and K-Nearest Neighbors (KNN). Input features were steady-state pERK, steady-state pAKT, and simulated NCG. Datasets were split into training and held-out test sets; each classifier was trained at each threshold and evaluated on the test set to avoid overfitting. Prediction accuracy across classifiers and thresholds is shown in Figure S5.2.1.

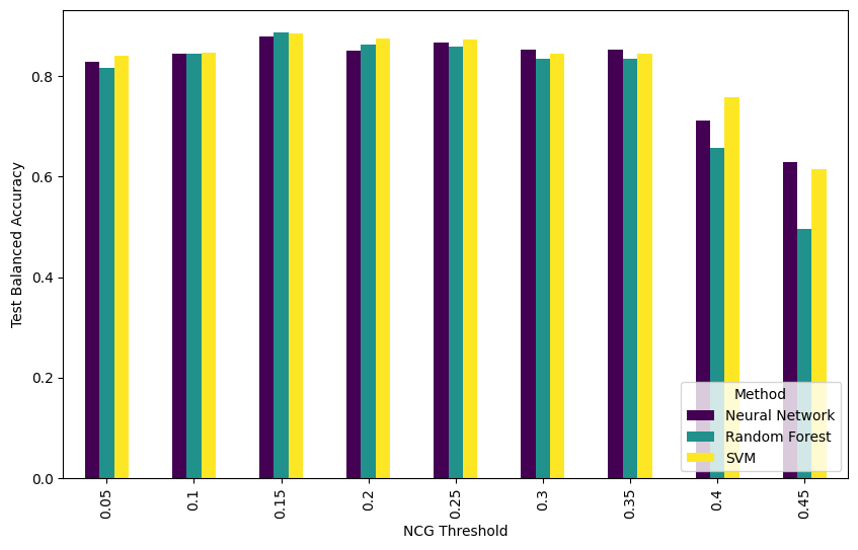

Figure S5.2.1 Trained supervised machine learning algorithm prediction accuracy with different tested threshold. Balanced accuracy on the held-out test set (30% of data) was evaluated across NCG probability thresholds (0–0.4) for all three classifiers - Neural Network (NN), Random Forest (RF), and Support Vector Machine (SVM). We identified 0.15 as the optimal threshold for binary classification of cell fate as proliferative (NCG ≥ 0.15) or quiescent (NCG < 0.15).

All three classifiers achieved peak prediction accuracy at NCG = 0.15, independently and consistently. This convergence across algorithms with different inductive biases provides robust quantitative evidence that NCG = 0.15 is the optimal threshold for distinguishing androgen-sensitive from androgen-resistant cell phenotypes in the MHS model. This value is used as the classification boundary throughout the main text analyses and in the machine learning surrogate training described in the main Methods.

### S6. Extended results from Survival Analysis

#### S6.1 KM analysis on cellular model surrogate driven Shapley ranking features

Table S6.1.1. KM Analysis on Cellular Model Surrogate-Driven Shapley Ranking Features 7-cohort combined molecular pool (OS endpoint).

| Gene | Alteration Type | Direction | N Altered | Median OS Altered (mo) | Median OS Wildtype (mo) | Log-rank p | Multivariate HR (95% CI) | p (cox) |
| --- | --- | --- | --- | --- | --- | --- | --- | --- |
| AR | Oncogene | Gain | 514 | 19.7 | 86.1 | 2.87E-131 | 4.07 [3.62-4.58] | 2.58E-122 |
| PTEN | Suppressor | Loss | 973 | 54.0 | 76.9 | 5.85E-12 | 1.51 [1.36-1.69] | 1.58E-13 |
| MDM4 | Oncogene | Gain | 73 | 32.9 | 74.6 | 1.86E-6 | 2.25 [1.66-3.05] | 1.70E-7 |
| MDM2 | Oncogene | Gain | 56 | 30.0 | 74.2 | 9.77E-6 | 2.20 [1.57-3.09] | 5.42E-6 |
| AKT1 | Oncogene | Gain | 99 | 52.4 | 74.6 | 6.64E-4 | 1.58 [1.18-2.11] | 2.38E-3 |
| CDKN1A | Suppressor | Loss | <20 † | - | - | - | - |  |
| ATM | Suppressor | Loss | 124 | 83.8 | 73.7 | 0.677 | 1.08 [0.81-1.44] | 0.619 |
| RAF1 | Oncogene | Gain | <20 † | - | - | - | - |  |
| BCL2 | Oncogene | Gain | <20 † | - | - | - | - |  |
| CASP9 | Suppressor | Loss | <20 † | - | - | - | - |  |
| BAX | Suppressor | Loss | <20 † | - | - | - | - |  |

**Footnote:** 1) Genes with N altered < 20 across the combined cohort were excluded from KM analysis; Cox HR is reported where ≥ 10 events were available. 2) † KM curve not generated due to insufficient altered arm size (n < 20). Cox HR estimated from pooled model.

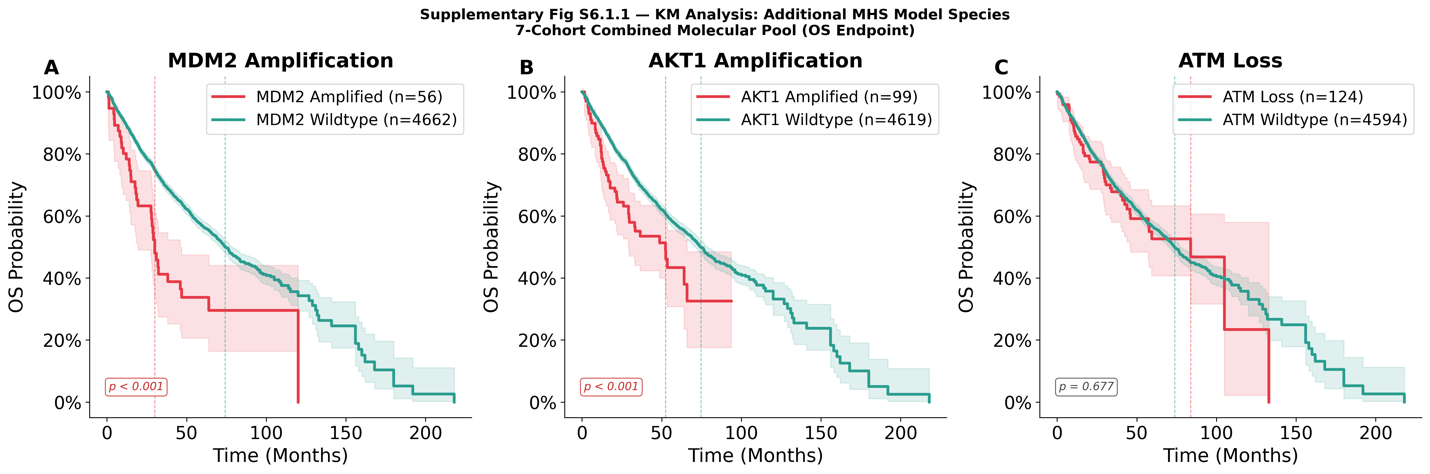

Figure S6.1.1**.** Kaplan-Meier overall survival curves for genes in the 7-cohort molecular pool not shown in the main manuscript. Patients are stratified by alteration status (altered vs. wildtype; strict CNA + truncating mutation thresholds). MDM2 and AKT1 reached statistical significance (log-rank p < 0.05); ATM did not. Significance threshold: p < 0.05*.*

#### S6.2 KM analysis on tissue level model: Androgen uptake and synthesis axis

The androgen uptake and synthesis axis was evaluated using two complementary approaches: per-cohort mRNA z-score analysis across four expression-eligible cohorts (TCGA-PRAD [primary], MSKCC [localized/early metastatic], SU2C [metastatic CRPC], MCTP [metastatic]) and genomic alteration analysis (CNA and somatic mutation combined via OR logic) across the 7-cohort molecular pool. Three genes spanning active androgen uptake (SLCO2B1, SLCO1B3) and intratumoral de novo androgen synthesis (AKR1C3) were assessed. mRNA stratification used a z-score threshold (z > 1.0 vs. rest), with cohort-specific endpoints: DFS for TCGA-PRAD and MSKCC; OS for SU2C. MCTP was excluded from KM analysis due to small cohort size.

Significant results for the three genes are discussed in the main text. Full results across all genes, cohorts, and z-score stratification strategy are reported in Table S6.2.1. Figure S6.2.1 contains KM curves (which did not reach significance) not added to the main text. Genomic alteration analysis for all three genes was not feasible due to insufficient altered arm sizes across the 7-cohort molecular pool (for SLCO2B1, SLCO1B3, AKR1C3), precluding reliable KM or Cox inference. For multivariate HR in single cohort analysis (mRNA expression), age was used as a covariate. When running multivariate HR for combined cohorts (genomic alteration analysis), the covariates were age and cohort names.

Table S6.2.1. mRNA Expression and genomic alteration KM Results: Androgen Uptake and Synthesis Axis

| **Gene** | **Analysis Type** | **Cohort** | **Endpoint** | **Stratification** | **N High / N Low** | **Log-rank p** | **Multivariate HR (95% CI)** | **p (Cox)** | **Direction** |
| --- | --- | --- | --- | --- | --- | --- | --- | --- | --- |
| AKR1C3 | mRNA Expression | TCGA-PRAD [Primary] | DFS | continuous z-score | — | — | 1.51 [1.25–1.83] | <0.001 | High is bad |
| AKR1C3 | mRNA Expression | TCGA-PRAD [Primary] | PFS | z-score threshold (z > 1) | 21 / 472 | 0.038 | 2.19 [1.01–4.75] | 0.046 | High is bad |
| AKR1C3 | mRNA Expression | MSKCC [Localized/Early Met.] | DFS | z-score threshold (z > 1) | 27 / 113 | — | 6.37 [2.17–18.73] | <0.001 | High is bad |
| AKR1C3 | mRNA Expression | SU2C [Metastatic CRPC] | OS | z-score threshold (z > 1) | — | — | 5.89 [1.25–27.72] | 0.025 | High is bad |
| AKR1C3 | mRNA Expression | MCTP [Metastatic] | OS | z-score threshold (z > 1) | — | — | 6.66 [1.65–26.97] | 0.008 | High is bad |
| SLCO2B1 | mRNA Expression | TCGA-PRAD [Primary] | DFS | z-score threshold (z > 1) | 33 / 300 | 0.467 | — | — | — |
| SLCO2B1 | mRNA Expression | TCGA-PRAD [Primary] | PFS | z-score threshold (z > 1) | 60 / 433 | 0.050 | 1.66 [0.97–2.85] | 0.066 | — |
| SLCO2B1 | mRNA Expression | MSKCC [Localized/Early Met.] | DFS | z-score threshold (z > 1) | 27 / 113 | 0.718 | — | — | — |
| SLCO1B3 | mRNA Expression | TCGA-PRAD [Primary] | PFS | z-score threshold (z > 1) | 30 / 463 | 0.433 | — | — | — |
| SLCO1B3 | mRNA Expression | MSKCC [Localized/Early Met.] | DFS | z-score threshold (z > 1) | 62 / 78 | 0.991 | — | — | — |
| SLCO2B1 | Genomic Alteration (CNA/Mut) | 7-Cohort Pool | OS | OR logic | Insufficient N (n=10) | — | — | — | — |
| SLCO1B3 | Genomic Alteration (CNA/Mut) | 7-Cohort Pool | OS | OR logic | Insufficient N (n=8) | — | — | — | — |
| AKR1C3 | Genomic Alteration (CNA/Mut) | 7-Cohort Pool | OS | OR logic | Insufficient N (n=5) | — | — | — | — |

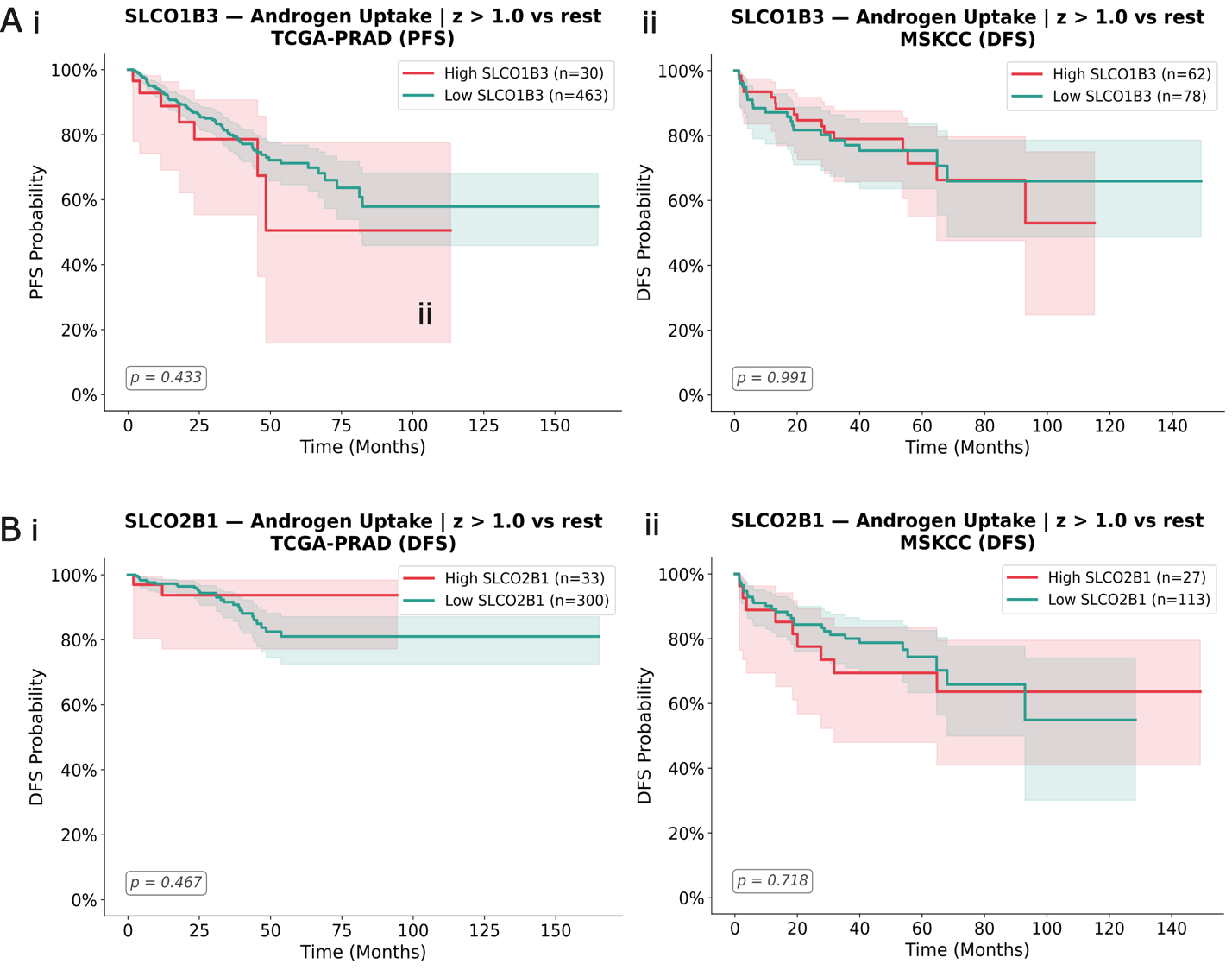

Figure S6.2.1 Kaplan-Meier survival curves for androgen uptake axis genes with non-significant mRNA expression associations. Patients were stratified using a z-score threshold (z > 1 vs. rest). All results were non-significant (log-rank p > 0.05). (A) SLCO1B3: (i) TCGA-PRAD progression-free survival (PFS; p = 0.433), (ii) MSKCC disease-free survival (DFS; p = 0.991). (B) SLCO2B1: (i) TCGA-PRAD disease-free survival (DFS; p = 0.467), (ii) MSKCC disease-free survival (DFS; p = 0.718). Dashed vertical lines indicate median survival where estimable.

#### S6.3 KM analysis on tissue level model: Adhesion/motility/cytoskeletal axis

The adhesion-motility axis was evaluated using two complementary approaches: per-cohort mRNA z-score analysis across four expression-eligible cohorts (TCGA-PRAD [primary], MSKCC [localized/early metastatic], SU2C [metastatic CRPC], MCTP [metastatic]) for ten genes spanning epithelial retention markers (CDH1, EPCAM), EMT drivers (VIM, CDH2, SNAI1), ECM remodeling and invasion (MMP9, FN1), cytoskeletal motility regulators (RHOA, ITGB1), and pro-invasive signaling (CXCL8); and genomic alteration analysis (CNA and somatic mutation combined via OR logic) across the 7-cohort molecular pool using OS as the primary endpoint. z-score threshold (z > 1.0 vs. rest) mRNA stratification strategy was applied. Endpoint was cohort-specific: DFS for TCGA-PRAD and MSKCC; OS for SU2C and MCTP. HR was calculated with adjustment for age as a covariate.

mRNA z-score threshold (z > 1 vs. rest) stratification was applied per cohort independently across TCGA-PRAD, MSKCC, and SU2C; MCTP was excluded from all analyses due to insufficient cohort size (N = 31). KM stratification was not feasible where the z > 1 group fell below the minimum arm size of 20 patients; Cox regression results are reported for these cases but should be interpreted cautiously in the absence of a supporting KM curve. MSKCC HR values reflect univariate Cox models as no covariates were available in that cohort. Full results across all ten genes and cohorts are reported in Table S6.3.1. Fig S6.3.1 gives KM curves for insignificant results not reported in the main text.

Table S6.3.1. mRNA Expression and genomic alteration KM Results: Adhesion and Motility Axis

| **Gene** | **Subgroup** | **Analysis Type** | **Cohort** | **Endpoint** | **Stratification** | **N High / N Low** | **Log-rank p** | **HR (95% CI)** | **p (Cox)** | **Direction** |
| --- | --- | --- | --- | --- | --- | --- | --- | --- | --- | --- |
| CDH1 | Epithelial | mRNA Expression | TCGA-PRAD [Primary] | DFS | z-score threshold (z > 1) | 287 / 46 | 0.892 | — | — | — |
| CDH1 | Epithelial | mRNA Expression | TCGA-PRAD [Primary] | PFS | z-score threshold (z > 1) | 435 / 58 | 0.048 | 2.34 [0.95–5.77] | 0.065 | Low is bad |
| CDH1 | Epithelial | mRNA Expression | MSKCC [Localized/Early Met.] | DFS | z-score threshold (z > 1) | 108 / 32 | 0.770 | — | — | — |
| CDH1 | Epithelial | mRNA Expression | SU2C [Metastatic CRPC] | OS | z-score threshold (z > 1) | — | — | 0.81 [0.34–1.91] | 0.631 | — |
| EPCAM | Epithelial | mRNA Expression | TCGA-PRAD [Primary] | DFS | z-score threshold (z > 1) | 290 / 43 | 0.667 | — | — | — |
| EPCAM | Epithelial | mRNA Expression | TCGA-PRAD [Primary] | PFS | z-score threshold (z > 1) | 425 / 68 | 0.666 | — | — | — |
| EPCAM | Epithelial | mRNA Expression | MSKCC [Localized/Early Met.] | DFS | z-score threshold (z > 1) | 36 / 104 | 0.887 | — | — | — |
| EPCAM | Epithelial | mRNA Expression | SU2C [Metastatic CRPC] | OS | z-score threshold (z > 1) | — | — | 0.55 [0.24–1.25] | 0.153 | — |
| VIM | Mesenchymal | mRNA Expression | TCGA-PRAD [Primary] | DFS | z-score threshold (z > 1) | 35 / 298 | 0.758 | — | — | — |
| VIM | Mesenchymal | mRNA Expression | TCGA-PRAD [Primary] | PFS | z-score threshold (z > 1) | 67 / 426 | 0.180 | — | — | — |
| VIM | Mesenchymal | mRNA Expression | SU2C [Metastatic CRPC] | OS | z-score threshold (z > 1) | — | — | 1.22 [0.50–2.98] | 0.657 | — |
| CDH2 | Mesenchymal | mRNA Expression | TCGA-PRAD [Primary] | DFS | z-score threshold (z > 1) | 25 / 308 | 0.485 | — | — | — |
| CDH2 | Mesenchymal | mRNA Expression | TCGA-PRAD [Primary] | PFS | z-score threshold (z > 1) | 32 / 461 | 0.660 | — | — | — |
| MMP9 | Mesenchymal | mRNA Expression | TCGA-PRAD [Primary] | PFS | z-score threshold (z > 1) | 34 / 459 | 0.455 | — | — | — |
| MMP9 | Mesenchymal | mRNA Expression | MSKCC [Localized/Early Met.] | DFS | z-score threshold (z > 1) | 61 / 79 | 0.692 | — | — | — |
| MMP9 | Mesenchymal | mRNA Expression | SU2C [Metastatic CRPC] | OS | z-score threshold (z > 1) | — | — | 1.76 [0.72–4.31] | 0.213 | — |
| MMP9 | Mesenchymal | mRNA Expression | MCTP [Metastatic] | OS | z-score threshold (z > 1) | — | — | 4.81 [1.24–18.71]† | 0.023 | High is bad |
| CXCL8 | Mesenchymal | mRNA Expression | TCGA-PRAD [Primary] | PFS | z-score threshold (z > 1) | 36 / 457 | 0.371 | — | — | — |
| SNAI1 | Mesenchymal | mRNA Expression | TCGA-PRAD [Primary] | DFS | z-score threshold (z > 1) | 32 / 301 | 0.498 | — | — | — |
| SNAI1 | Mesenchymal | mRNA Expression | TCGA-PRAD [Primary] | PFS | z-score threshold (z > 1) | 59 / 434 | 0.393 | — | — | — |
| SNAI1 | Mesenchymal | mRNA Expression | MSKCC [Localized/Early Met.] | DFS | z-score threshold (z > 1) | 28 / 112 | 0.615 | — | — | — |
| RHOA | Mesenchymal | mRNA Expression | TCGA-PRAD [Primary] | DFS | z-score threshold (z > 1) | 42 / 291 | 0.866 | — | — | — |
| RHOA | Mesenchymal | mRNA Expression | TCGA-PRAD [Primary] | PFS | z-score threshold (z > 1) | 84 / 409 | 0.017 | 1.80 [1.11–2.90] | 0.016 | High is bad |
| RHOA | Mesenchymal | mRNA Expression | MSKCC [Localized/Early Met.] | DFS | z-score threshold (z > 1) | — | — | 3.44 [1.32–8.93]† | 0.011 | High is bad |
| RHOA | Mesenchymal | mRNA Expression | SU2C [Metastatic CRPC] | OS | z-score threshold (z > 1) | — | — | 0.66 [0.20–2.16] | 0.492 | — |
| ITGB1 | Mesenchymal | mRNA Expression | TCGA-PRAD [Primary] | DFS | z-score threshold (z > 1) | 34 / 299 | 0.720 | — | — | — |
| ITGB1 | Mesenchymal | mRNA Expression | TCGA-PRAD [Primary] | PFS | z-score threshold (z > 1) | 54 / 439 | 0.305 | — | — | — |
| FN1 | Mesenchymal | mRNA Expression | TCGA-PRAD [Primary] | DFS | z-score threshold (z > 1) | — | — | 4.59 [0.62–34.0] | 0.135 | — |
| FN1 | Mesenchymal | mRNA Expression | TCGA-PRAD [Primary] | PFS | z-score threshold (z > 1) | — | — | 2.65 [0.83–8.40] | 0.099 | — |
| FN1 | Mesenchymal | mRNA Expression | MSKCC [Localized/Early Met.] | DFS | z-score threshold (z > 1) | — | — | 3.97 [1.80–8.76]† | 0.001 | High is bad |
| FN1 | Mesenchymal | mRNA Expression | SU2C [Metastatic CRPC] | OS | z-score threshold (z > 1) | — | — | 0.78 [0.19–3.25] | 0.730 | — |
| CDH1 | Epithelial | Genomic Alteration (CNA/Mut) | 7-Cohort Pool | OS | CNA (GISTIC ≤ −2 or ≥ +2) OR truncating mutation OR SV | N altered = 71 | 0.108 | — | — | — |
| EPCAM | Epithelial | Genomic Alteration (CNA/Mut) | 7-Cohort Pool | OS | CNA (GISTIC ≤ −2 or ≥ +2) OR truncating mutation OR SV | N altered = 27 | 0.088 | — | — | — |
| Other |  | Genomic Alteration (CNA/Mut) | 7-Cohort Pool | OS | CNA (GISTIC ≤ −2 or ≥ +2) OR truncating mutation OR SV | Insufficient N (<20 | — | — | — | — |

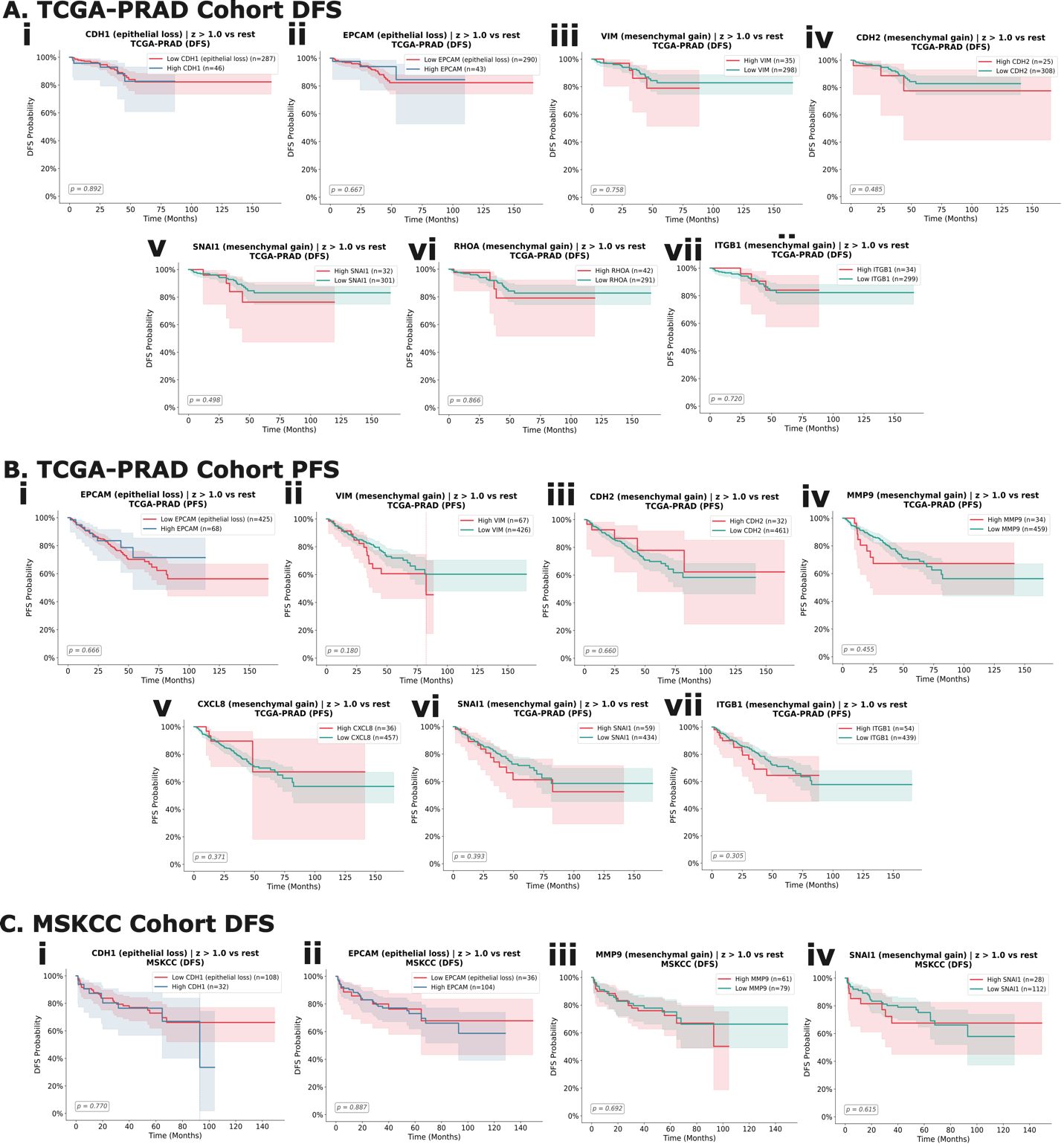

Fig S6.3.1 Kaplan-Meier Survival Analysis for Adhesion-Motility Axis: Per-Cohort mRNA Expression

Kaplan-Meier survival analysis of adhesion-motility axis mRNA expression in TCGA-PRAD and MSKCC cohorts. Patients were stratified using a z-score threshold (z > 1 vs. rest). All results were non-significant (log-rank p > 0.05). (A) TCGA-PRAD disease-free survival (DFS): (i) CDH1, (ii) EPCAM, (iii) VIM, (iv) CDH2, (v) SNAI1, (vi) RHOA, (vii) ITGB1. (B) TCGA­-PRA­­­D progression-free survival (PFS): (i) EPCAM, (ii) VIM, (iii) CDH2, (iv) MMP9, (v) CXCL8, (vi) SNAI1, (vii) ITGB1. (C) MSKCC disease-free survival (DFS): (i) CDH1, (ii) EPCAM, (iii) MMP9, (iv) SNAI1. Dashed vertical lines indicate median survival where estimable. Log-rank p-values are annotated on each panel. FN1 and RHOA in MSKCC, and MMP9 in MCTP, are not shown as KM stratification was not feasible due to insufficient high-expression group sizes.

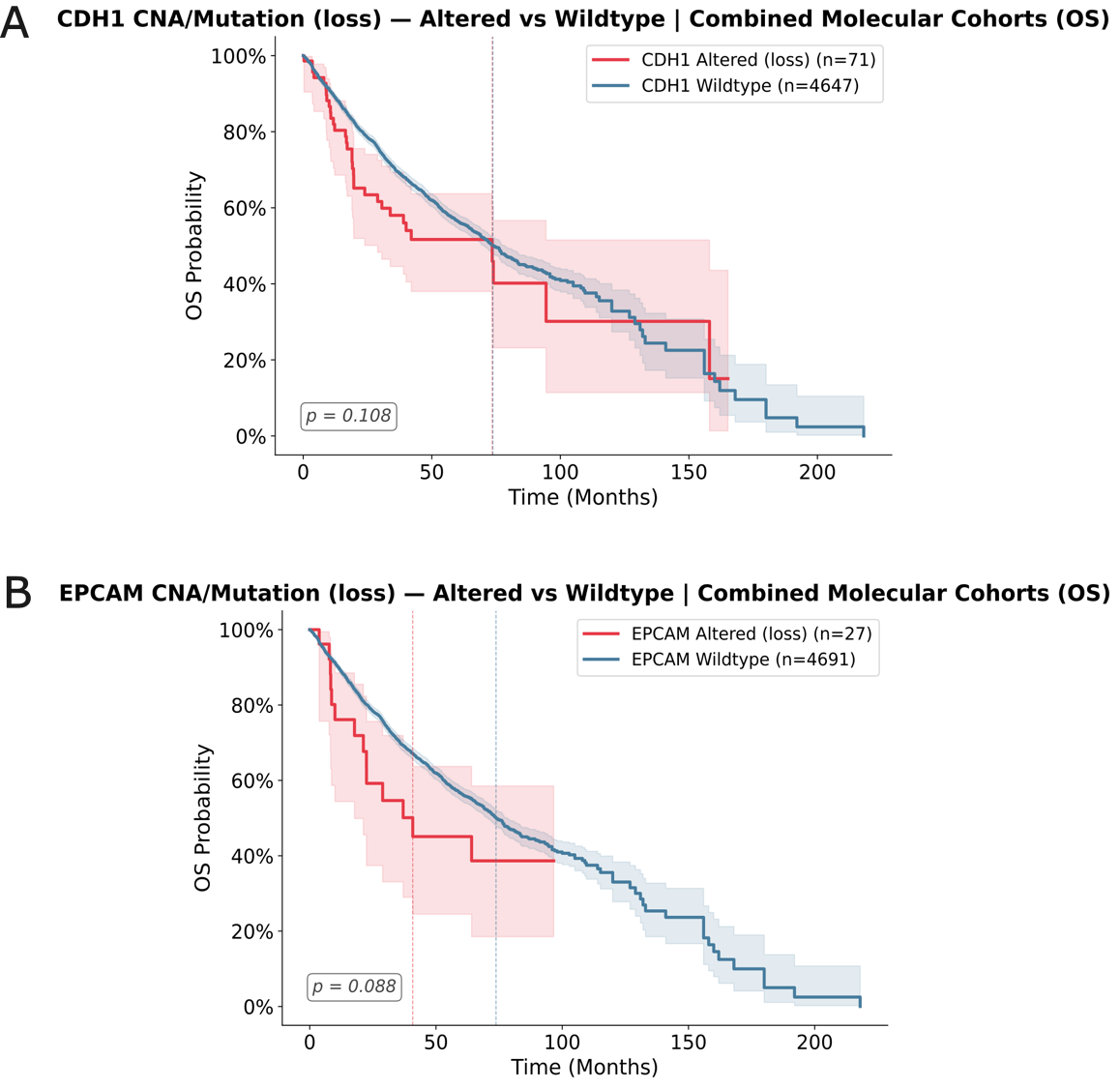

Figure S6.3.2 Genomic alteration analysis of adhesion-motility axis genes across the 7-cohort molecular pool. Kaplan-Meier overall survival curves for patients with vs. without genomic alterations (copy number alteration [GISTIC ≤ −2 or ≥ +2], truncating somatic mutation, or structural variant; combined via OR logic). (A) CDH1 (epithelial loss; N altered = 71; log-rank p = 0.108). (B) EPCAM (epithelial loss; N altered = 27; log-rank p = 0.088). Neither CDH1 nor EPCAM reached statistical significance. Mesenchymal genes (VIM, CDH2, MMP9, CXCL8, SNAI1, RHOA, ITGB1, FN1) are not shown as altered group sizes were insufficient for KM inference (N < 20) across the 7-cohort pool.

#### S6.4 KM analysis on tissue level model: Crowding / Mechanotransduction axis

The crowding / mechanotransduction axis was evaluated by per-cohort mRNA z-score analysis in TCGA-PRAD (n = 494, primary disease) (from cBioportal), the cohort with paired expression and recurrence endpoints, for 27 genes spanning six pathways that normally enforce crowding-induced growth arrest: (1) Hippo/YAP-TAZ contact inhibition (YAP1, WWTR1, LATS1, LATS2); (2) mechanosensitive ion channels(PIEZO1, TRPV4); (3) phosphoinositide kinases (PIP4K2B, PIP5K1C, PIK3CA); (4) cell-cycle arrest regulators (CDKN1A/p21, CDKN1B/p27, CCND1, MKI67); (5) hypoxia mediators (HIF1A, EPAS1, VEGFA); and (6) a PIP2-trafficking / cytoskeletal mechanosensing signature derived from prostate cancer cell-line proteomics ^17^ (ANXA1, KPNA4, RAE1, PLS1, ZYX, ARF1, CDC42, EZR, LAMP2, HMOX1, FLNC). Patients were stratified by median expression (above vs. below the cohort median) for all genes except ANXA1, which used a more robust top-vs-bottom quartile split. Cohort-specific endpoints were disease-free survival (DFS) and progression-free survival (PFS). Overall survival (OS) was not analyzed because TCGA-PRAD is effectively OS-event-free (10/494 events, 2%).

Each gene was additionally tested for a gene × PTEN-deletion interaction (PTEN deep deletion, GISTIC = −2; 85 of 494 tumors, PTEN-Resistant; the remainder PTEN-Sensitive), asking whether a gene's prognostic effect differs between PTEN-deleted and PTEN-intact tumors. Full per-gene KM results are reported in Table S6.4.1 and the interaction results in Table S6.4.2. Figure S6.4.1summarizes the significant interactions; KM curves not shown in the main text are in Figure S6.4.2.

**Table S6.4.1. mRNA Expression KM Results: Crowding / Mechanotransduction Axis**

| **Gene** | **Axis** | **Endpoint** | **Stratification** | **N High / N Low** | **Log-rank p** | **HR (95% CI)** | **p (Cox)** | **Direction** |
| --- | --- | --- | --- | --- | --- | --- | --- | --- |
| YAP1 | Hippo/YAP-TAZ | DFS | Median split | 170 / 163 | 0.037 | 0.62 [0.43–0.90] | 0.011 | High is bad |
| YAP1 | Hippo/YAP-TAZ | PFS | Median split | 246 / 247 | 0.180 | 0.80 [0.66–0.98] | 0.028 | High is bad |
| WWTR1 | Hippo/YAP-TAZ | DFS | Median split | 168 / 165 | 0.078 | 0.47 [0.28–0.79] | 0.004 | High is bad |
| WWTR1 | Hippo/YAP-TAZ | PFS | Median split | 246 / 247 | 0.228 | 0.71 [0.55–0.92] | 0.009 | High is bad |
| LATS1 | Hippo/YAP-TAZ | DFS | Median split | 171 / 162 | 0.090 | 0.58 [0.33–1.02] | 0.056 | Low is bad |
| LATS1 | Hippo/YAP-TAZ | PFS | Median split | 246 / 247 | 0.330 | 0.73 [0.53–0.99] | 0.042 | Low is bad |
| LATS2 | Hippo/YAP-TAZ | DFS | Median split | 166 / 167 | 0.413 | 0.68 [0.45–1.02] | 0.063 | Low is bad |
| LATS2 | Hippo/YAP-TAZ | PFS | Median split | 246 / 247 | 0.387 | 0.83 [0.68–1.03] | 0.088 | Low is bad |
| PIEZO1 | Mechanosensitive ion channel | DFS | Median split | 177 / 156 | 0.275 | 1.33 [0.96–1.85] | 0.087 | High is bad |
| PIEZO1 | Mechanosensitive ion channel | PFS | Median split | 246 / 247 | 0.666 | 1.14 [0.93–1.39] | 0.201 | High is bad |
| TRPV4 | Mechanosensitive ion channel | DFS | Median split | 173 / 160 | 0.977 | 0.90 [0.59–1.35] | 0.602 | High is bad |
| TRPV4 | Mechanosensitive ion channel | PFS | Median split | 246 / 247 | 0.922 | 1.00 [0.80–1.23] | 0.968 | High is bad |
| PIP4K2B | Phosphoinositide kinase | DFS | Median split | 173 / 160 | 0.147 | 1.27 [0.92–1.76] | 0.149 | High is bad |
| PIP4K2B | Phosphoinositide kinase | PFS | Median split | 246 / 247 | 0.233 | 1.12 [0.94–1.32] | 0.198 | High is bad |
| PIP5K1C | Phosphoinositide kinase | DFS | Median split | 175 / 158 | 0.165 | 1.08 [0.83–1.41] | 0.563 | High is bad |
| PIP5K1C | Phosphoinositide kinase | PFS | Median split | 246 / 247 | 0.920 | 0.95 [0.79–1.14] | 0.566 | High is bad |
| PIK3CA | Phosphoinositide kinase | DFS | Median split | 165 / 168 | 0.022 | 0.62 [0.38–0.98] | 0.043 | High is bad |
| PIK3CA | Phosphoinositide kinase | PFS | Median split | 246 / 247 | 0.308 | 0.82 [0.64–1.04] | 0.098 | High is bad |
| CDKN1A | Cell-cycle arrest | DFS | Median split | 162 / 171 | 0.457 | 1.08 [0.81–1.46] | 0.597 | Low is bad |
| CDKN1A | Cell-cycle arrest | PFS | Median split | 246 / 247 | 0.206 | 0.92 [0.73–1.14] | 0.435 | Low is bad |
| CDKN1B | Cell-cycle arrest | DFS | Median split | 157 / 176 | 0.119 | 0.79 [0.55–1.14] | 0.210 | Low is bad |
| CDKN1B | Cell-cycle arrest | PFS | Median split | 246 / 247 | 0.829 | 1.09 [0.90–1.32] | 0.405 | Low is bad |
| CCND1 | Cell-cycle arrest | DFS | Median split | 164 / 169 | 0.033 | 1.08 [0.90–1.29] | 0.414 | High is bad |
| CCND1 | Cell-cycle arrest | PFS | Median split | 246 / 247 | 0.049 | 1.01 [0.84–1.21] | 0.955 | High is bad |
| MKI67 | Cell-cycle arrest | DFS | Median split | 150 / 183 | 0.104 | 1.22 [0.93–1.60] | 0.151 | High is bad |
| MKI67 | Cell-cycle arrest | PFS | Median split | 246 / 247 | 0.001 | 1.38 [1.21–1.59] | < 0.001 | High is bad |
| HIF1A | Hypoxia | DFS | Median split | 166 / 167 | 0.005 | 0.47 [0.27–0.82] | 0.008 | High is bad |
| HIF1A | Hypoxia | PFS | Median split | 246 / 247 | 0.057 | 0.88 [0.69–1.11] | 0.270 | High is bad |
| EPAS1 | Hypoxia | DFS | Median split | 175 / 158 | 0.360 | 0.97 [0.60–1.58] | 0.901 | High is bad |
| EPAS1 | Hypoxia | PFS | Median split | 246 / 247 | 0.118 | 0.91 [0.67–1.23] | 0.532 | High is bad |
| VEGFA | Hypoxia | DFS | Median split | 178 / 155 | 0.972 | 1.39 [1.03–1.87] | 0.030 | High is bad |
| VEGFA | Hypoxia | PFS | Median split | 246 / 247 | 0.761 | 1.16 [0.95–1.42] | 0.135 | High is bad |
| ANXA1 | PIP2/cytoskeletal | DFS | Top vs bottom quartile | 89 / 83 | 0.494 | 0.83 [0.55–1.24] | 0.355 | High is bad |
| ANXA1 | PIP2/cytoskeletal | PFS | Top vs bottom quartile | 124 / 124 | 0.318 | 0.87 [0.69–1.09] | 0.215 | High is bad |
| KPNA4 | PIP2/cytoskeletal | DFS | Median split | 158 / 175 | 0.145 | 0.85 [0.63–1.16] | 0.314 | High is bad |
| KPNA4 | PIP2/cytoskeletal | PFS | Median split | 246 / 247 | 0.170 | 0.94 [0.79–1.10] | 0.428 | High is bad |
| RAE1 | PIP2/cytoskeletal | DFS | Median split | 150 / 183 | 0.588 | 1.23 [0.95–1.61] | 0.122 | High is bad |
| RAE1 | PIP2/cytoskeletal | PFS | Median split | 246 / 247 | 0.210 | 1.36 [1.19–1.56] | < 0.001 | High is bad |
| PLS1 | PIP2/cytoskeletal | DFS | Median split | 186 / 147 | 0.025 | 0.66 [0.43–1.01] | 0.055 | High is bad † |
| PLS1 | PIP2/cytoskeletal | PFS | Median split | 246 / 247 | 0.002 | 0.77 [0.61–0.96] | 0.018 | High is bad † |
| ZYX | PIP2/cytoskeletal | DFS | Median split | 158 / 175 | 0.926 | 1.04 [0.76–1.44] | 0.792 | High is bad |
| ZYX | PIP2/cytoskeletal | PFS | Median split | 246 / 247 | 0.906 | 1.07 [0.89–1.28] | 0.470 | High is bad |
| ARF1 | PIP2/cytoskeletal | DFS | Median split | 168 / 165 | 0.820 | 1.08 [0.82–1.43] | 0.588 | High is bad |
| ARF1 | PIP2/cytoskeletal | PFS | Median split | 246 / 247 | 0.571 | 0.91 [0.76–1.08] | 0.293 | High is bad |
| CDC42 | PIP2/cytoskeletal | DFS | Median split | 160 / 173 | 0.371 | 1.00 [0.73–1.39] | 0.989 | High is bad |
| CDC42 | PIP2/cytoskeletal | PFS | Median split | 246 / 247 | 0.256 | 1.04 [0.86–1.25] | 0.716 | High is bad |
| EZR | PIP2/cytoskeletal | DFS | Median split | 166 / 167 | 0.018 | 1.13 [0.81–1.58] | 0.458 | High is bad |
| EZR | PIP2/cytoskeletal | PFS | Median split | 246 / 247 | 0.388 | 0.98 [0.80–1.21] | 0.859 | High is bad |
| LAMP2 | PIP2/cytoskeletal | DFS | Median split | 180 / 153 | 0.483 | 0.76 [0.52–1.11] | 0.159 | High is bad |
| LAMP2 | PIP2/cytoskeletal | PFS | Median split | 246 / 247 | 0.054 | 0.70 [0.56–0.88] | 0.002 | High is bad |
| HMOX1 | PIP2/cytoskeletal | DFS | Median split | 149 / 184 | 0.786 | 1.19 [0.98–1.45] | 0.081 | High is bad |
| HMOX1 | PIP2/cytoskeletal | PFS | Median split | 246 / 247 | 0.062 | 1.18 [1.05–1.33] | 0.006 | High is bad |
| FLNC | PIP2/cytoskeletal | DFS | Median split | 172 / 161 | 0.095 | 0.69 [0.42–1.15] | 0.155 | High is bad |
| FLNC | PIP2/cytoskeletal | PFS | Median split | 246 / 247 | 0.035 | 0.71 [0.54–0.95] | 0.021 | High is bad |

*Footnote: HR is the univariate Cox hazard ratio per +1 standard-deviation increase in mRNA z-score (continuous), reported alongside the median/quartile log-rank test. "Direction" is the prior expectation from the source pathway/proteomic signature.

† PLS1 was annotated "high is bad" from the cross-cancer proteomic signature, but in TCGA-PRAD higher PLS1 expression was associated with better outcome (protective HR); its prostate-specific direction is unconfirmed.

Table S6.4.2. Gene × PTEN-deletion interaction (Cox): Crowding / Mechanotransduction Axis

Interaction term from a Cox model gene + PTEN-deletion + (gene × PTEN-deletion), TCGA-PRAD. Only terms reaching nominal significance (interaction p < 0.05) are listed. An interaction HR > 1 indicates the gene's hazard is amplified in PTEN-deleted tumors.

| **Gene** | **Axis (A#)** | **Endpoint** | **N PTEN-deleted** | **Events (total)** | **Interaction HR** | **Interaction p** |
| --- | --- | --- | --- | --- | --- | --- |
| CCND1 | A4 Cell-cycle arrest | DFS | 55 | 29 | 5.40 | 1.6 × 10⁻⁴ |
| CCND1 | A4 Cell-cycle arrest | PFS | 85 | 91 | 2.78 | 0.015 |
| ANXA1 | A6 PIP2/cytoskeletal | PFS | 85 | 91 | 0.48 | 0.021 |
| YAP1 | A1 Hippo/YAP-TAZ | PFS | 85 | 91 | 0.53 | 0.021 |
| CDKN1A | A4 Cell-cycle arrest | PFS | 85 | 91 | 0.24 | 0.022 |
| EPAS1 | A5 Hypoxia | PFS | 85 | 91 | 0.29 | 0.023 |
| WWTR1 | A1 Hippo/YAP-TAZ | PFS | 85 | 91 | 0.44 | 0.024 |
| LATS2 | A1 Hippo/YAP-TAZ | PFS | 85 | 91 | 0.55 | 0.031 |
| PIK3CA | A 3 Phosphoinositide kinase | PFS | 85 | 91 | 0.47 | 0.035 |
| PLS1 | A6 PIP2/cytoskeletal | DFS | 55 | 29 | 0.33 | 0.048 |

*Footnote: events are total recurrence events in the model (PTEN-deleted + sensitive). Interaction estimates on DFS rest on 29 events (8 in the PTEN-deleted arm), this should be interpreted only as a preliminary exploratory analysis.

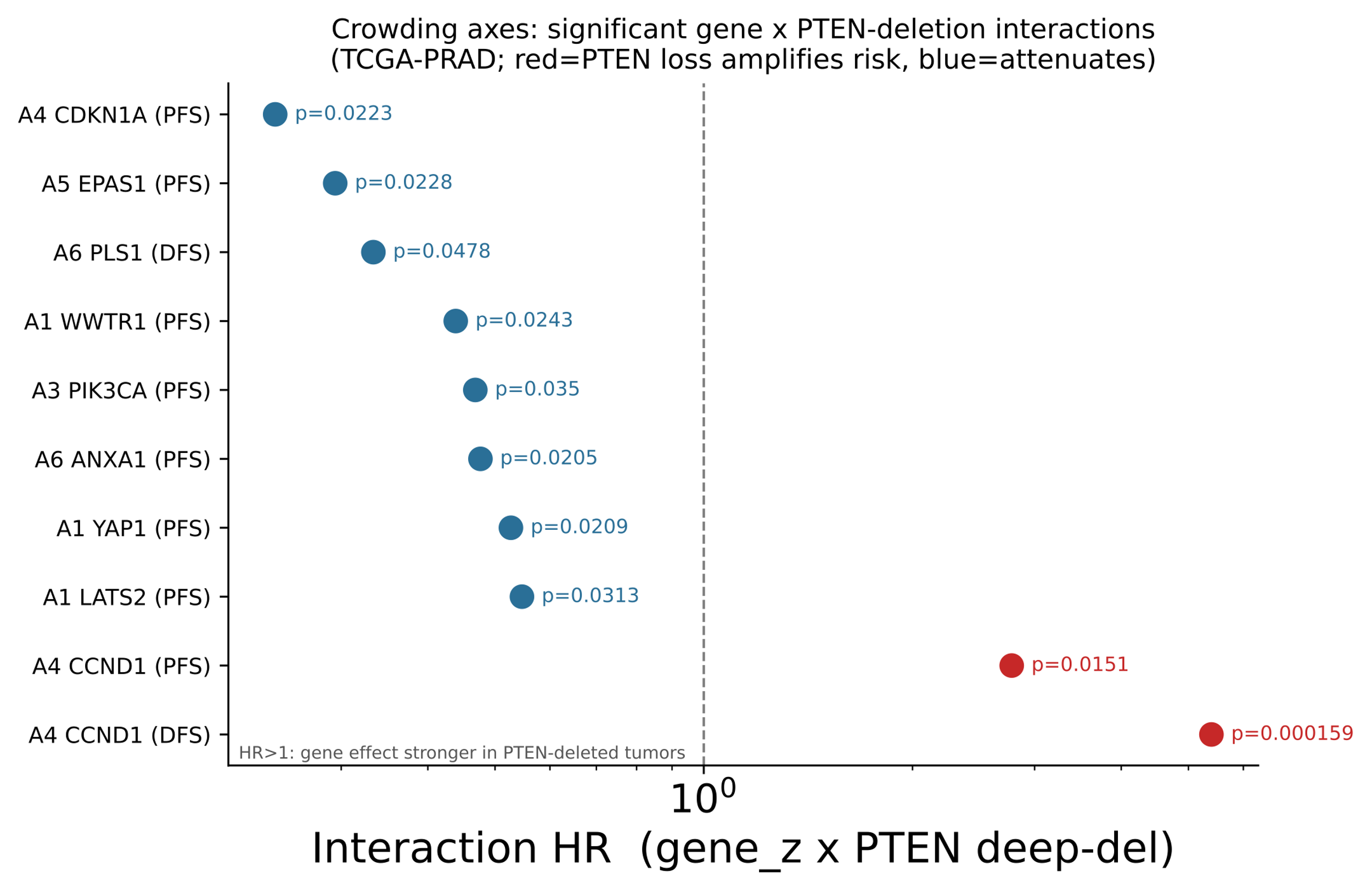

Figure S6.4.1

Forest plot of the gene × PTEN-deletion interaction terms reaching nominal significance (p < 0.05) across the six crowding axes in TCGA-PRAD. Points are interaction hazard ratios on a log scale; an HR > 1 (red) indicates the gene's prognostic effect is amplified in PTEN-deleted tumors, HR < 1 (blue) attenuated. CCND1 on DFS is the strongest interaction (HR 5.40, p = 1.6 × 10⁻⁴). A# associated with the genes are listed in the Table S6.4.2.

### S7. Agent-Based Model Implementation, Parameters, and Multi-scale modeling Simulation Design

#### S7.1. Model Framework

*Domain:*

The tissue-scale agent-based model was implemented in PhysiCell (v1.10.4) ^18^, an open-source physics-based multicellular simulator that couples discrete cell - level mechanics and phenotypic rules with spatiotemporal diffusion of microenvironmental substrates via BioFVM. ^19^ The ABM operates in 2D on a rectilinear domain of 500 × 500 × 20 µm with a uniform voxel resolution of 20 µm. All simulations were run for 15 months (648,000 simulation minutes), with outputs saved every 6,480 minutes (approximately 4.5 days), yielding 101 time points per run.

*Microenvironment:*

Two diffusible substrates were modeled: oxygen and testosterone. Oxygen was assigned a diffusion coefficient of 100 µm²/min with Dirichlet boundary conditions of 38 mmHg on all domain boundaries, representing well-oxygenated peri-vascular tissue, and a cellular uptake rate of 0.1 min⁻¹ for both cell types. ^19,20^ Necrosis was triggered below a local pO₂ of 5 mmHg, with a linear ramp from 0 to 1 over the 5-7 mmHg range governing the oxygen-dependent proliferation multiplier. Testosterone was assigned a diffusion coefficient of 1 µm²/min (estimated by scaling the oxygen diffusion coefficient by molecular weight via the Stokes-Einstein relation, D_T ≈ D_O₂·(MW_O₂/MW_T)^(1/3), and applying an additional dampening factor to account for hindered steroid transport through the extracellular matrix), boundary concentrations of 8 ng/mL under physiological androgen conditions and 1 ng/mL to represent androgen deprivation therapy (ADT). Both substrates had zero decay and were initialized at their respective boundary values. Microenvironmental gradients were governed by the reaction-diffusion equation:

∂c/∂t = ∇²(D∇c) − U(x,y,t) + S(x,y,t)

where D is the diffusion coefficient, U(x,y,t) is the cellular uptake rate field (nonzero only in voxels occupied by agents and dependent on agent genotype), and S(x,y,t) represents source and sink terms.

*Agent Initialization and Cell Types:*

Two cell populations were modeled, representing the clonal heterogeneity observed at biochemical recurrence: androgen-sensitive cells (PTEN-normal, S) and androgen-resistant cells (PTEN-deleted, R). Unless otherwise specified in individual parameter sweeps, simulations were initialized with 150 cancer cells placed randomly in a circular arrangement under normoxia and physiological androgen concentrations.

*Cell Proliferation rules:*

The net growth rate for each agent was governed by a testosterone- and oxygen-dependent Hill function derived from MHS model outputs:

λ = λ_O₂ · r[1 + (m/r) · Tⁿ / (pⁿ + Tⁿ)] · κ / (30 × 24 × 60)

where T is local testosterone concentration, r is the basal growth rate, m is the maximum testosterone-stimulated growth rate, n is the Hill exponent governing androgen response sharpness, p is the half-saturation concentration, and κ is a cohort-specific scaling factor. This represents the two-population model. Each cohort (BR, TR and Control) had its own set of params fitted. Details on this in Supplementary Section S.1.4

***Cell Mechanics and Motility:***

Cell–cell interactions and parameters followed PhysiCell's spring-adhesion model.^18^ Cells within 1.25× the cell diameter exert attractive forces governed by a cell–cell adhesion strength parameter (baseline 0.4 µm/min), while overlapping cells generate repulsive forces (cell–cell repulsion strength = 10.0 µm/min). Cell-basement membrane adhesion and repulsion strengths were set to 4.0 and 10.0 µm/min respectively. Random motility was implemented as Brownian-like displacement with a baseline migration speed of 0.4 µm/min, a persistence time of 1 min, and a migration bias of 0.5. Chemotaxis was disabled. Both cell types shared identical baseline mechanics and motility parameters, with deviations applied only in the parameter sweep conditions.

*Cellular Testosterone uptake:*

Cellular testosterone uptake was modeled as a first-order process governed by a per-cell uptake rate parameter (baseline 1 × 10⁻⁵ min⁻¹) ^21,22^, applied uniformly to both PTEN-normal and PTEN-deleted cell types under nominal conditions. Both cell types shared identical baseline uptake parameters, with deviations applied in the parameter sweep conditions.

#### S7.2. Coupling MHS model to Tissue-level ABM

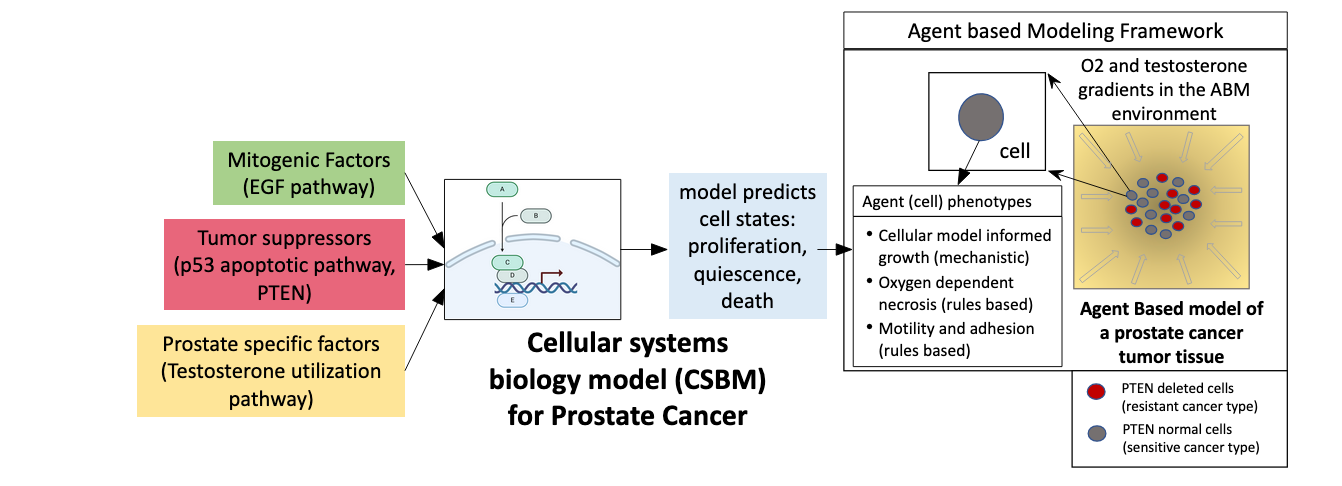

Fig S7.2.1 Multiscale modeling framework architecture.

The prostate cancer ABM integrates patient multi-omics data and serves as a conduit between cellular-scale signaling and tissue-scale environmental dynamics. Cohort-specific multi-omics data, derived from The Cancer Genome Atlas (TCGA) transcriptomic profiles, are mapped onto nodes of the MHS cellular model, propagating patient-specific signaling states through the agent-based simulation to generate cohort-differentiated tissue-scale progression dynamics. The MHS model serves as the intracellular logic engine of the PhysiCell ABM, coupling cohort-specific genetic states to local microenvironmental gradients of testosterone, mitogen, and physical crowding. Intrinsic clonal fitness determines how individual agents respond to and reshape their microenvironment, while extrinsic microenvironmental signals modulate the competitive outcomes of PTEN-resistant and PTEN-sensitive clones.

#### S7.3. ABM - ODE Consistency Check

The main text results combine two modeling layers: the two-population ODE sub-model (MHS cellular model) and the spatial agent-based model (PhysiCell ABM). A prerequisite for interpreting differences between ODE and ABM outputs as biologically meaningful spatial effects, rather than numerical artifacts of the modeling framework, is that the two layers produce consistent results under equivalent, spatially unconstrained conditions. This section reports that consistency check.

ABM simulations were initialized under low-density conditions (150 cells in a circular arrangement with no contact inhibition enforced), which approximate the spatially unconstrained regime in which the ODE model operates. PSA outputs were generated for all three cohorts (CNT, BR, TR) at low (1 ng/mL) and high (8 ng/mL) testosterone concentrations. Five stochastic ABM replicates were run per condition; ODE solutions were computed at matched testosterone values using the MATLAB implementation of the two-population sub model.

ODE-predicted PSA values fell within one standard deviation of the ABM simulation distributions across all cohorts and testosterone concentrations (Figure S7.3.1). This agreement confirms that the two model layers are built on a consistent mathematical and biological foundation: differences in spatial organization, clustering dynamics, and tissue-level outcomes reported in the main text ABM analyses reflect emergent spatial effects rather than inter-model inconsistencies. It further validates the use of ODE-derived Hill function parameters as the governing proliferative response functions for ABM agent phenotypes across cohorts.

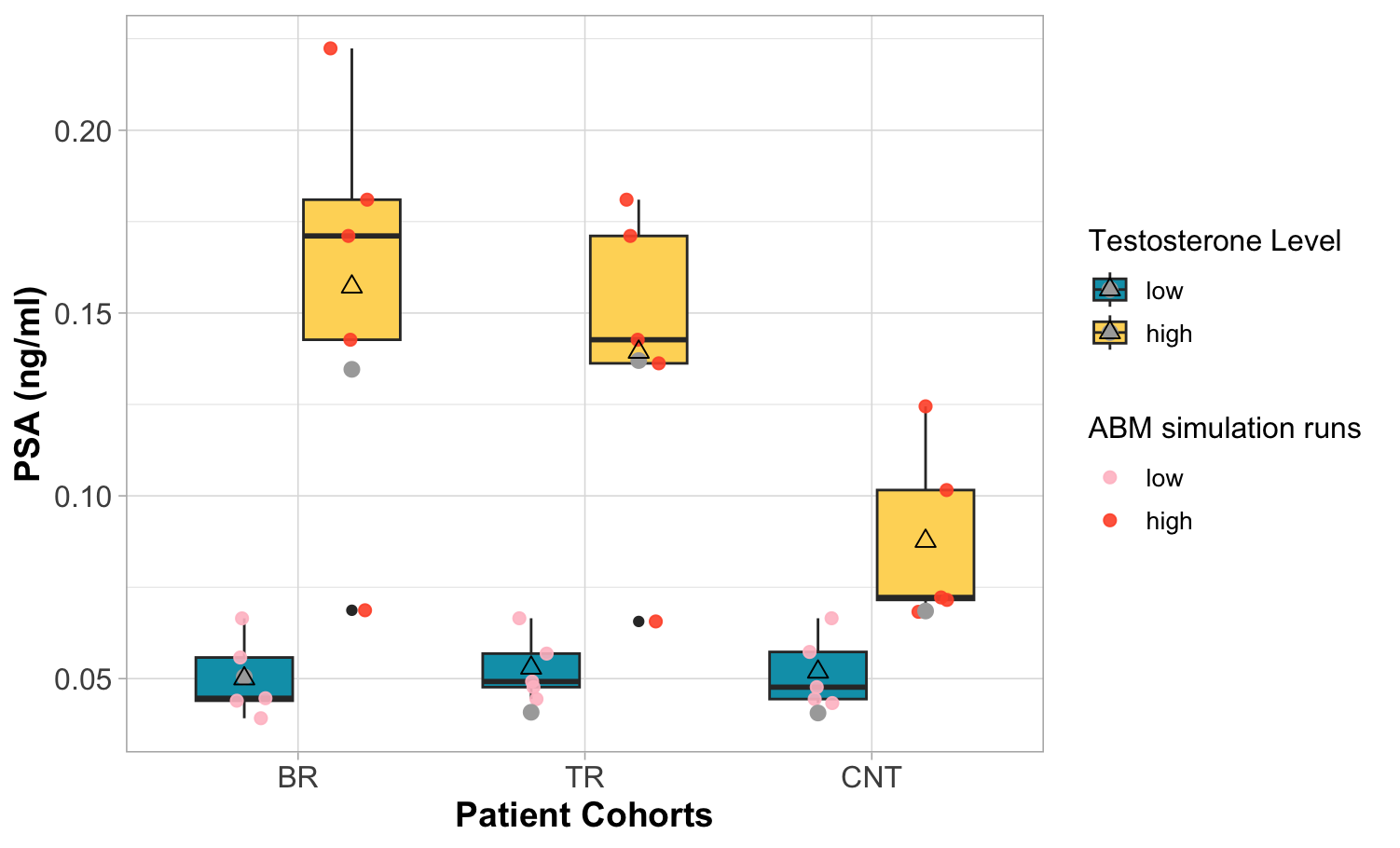

**Figure S7.3.1** *ABM–ODE consistency across cohorts and testosterone concentrations. Box plots show ABM PSA distributions (5 stochastic replicates per condition; mean indicated by hollow triangle). Blue: low testosterone (1 ng/mL); yellow: high testosterone (8 ng/mL). Individual ABM replicates shown as points (pink: low T; orange: high T). ODE solutions shown as filled grey circles. Results are shown for CNT, BR, and TR cohorts.*

#### S7.4. Comparing 2D vs 3D ABM simulations

Across both 2D and 3D geometries, ADT reduced total tumor cell count by approximately 54% in 2D (176 ± 14 vs. 381 ± 19 cells) and 60% in 3D (171 ± 8 vs. 425 ± 54 cells) relative to normal androgen conditions (p < 0.001, Mann-Whitney U for both PTEN-normal and PTEN-deleted cell counts), consistent with expected growth suppression of the androgen-sensitive population. (Fig S7.4.1)

The R/S ratio was lower under ADT than under normal androgen in both geometries. The two ADT conditions converged to nearly identical total cell counts regardless of geometry (171 ± 8 in 3D vs. 176 ± 14 in 2D, p = 0.173, Mann-Whitney U).

A notable geometric difference was observed in the R/S ratio specifically. Under 3D geometry, ADT produced a statistically significant reduction in R/S ratio (0.585 ± 0.135 vs. 0.445 ± 0.077, p = 0.021, Mann-Whitney U), whereas in 2D this difference did not reach significance (0.500 ± 0.131 vs. 0.456 ± 0.114, p = 0.345). This suggests that 3D spatial structure amplifies ADT's selective pressure on clonal dynamics in a way that 2D geometry does not capture.

Furthermore, normal androgen conditions in 3D produced substantially greater outcome variability across seeds than in 2D, whereas ADT collapsed 3D variability to below even 2D normal androgen levels (SD = 0.077 under 3D ADT). This indicates that testosterone-driven proliferation is a primary driver of outcome heterogeneity amplification in three dimensions, and that ADT suppresses not only mean tumor size but also the stochastic spread of clonal outcomes.

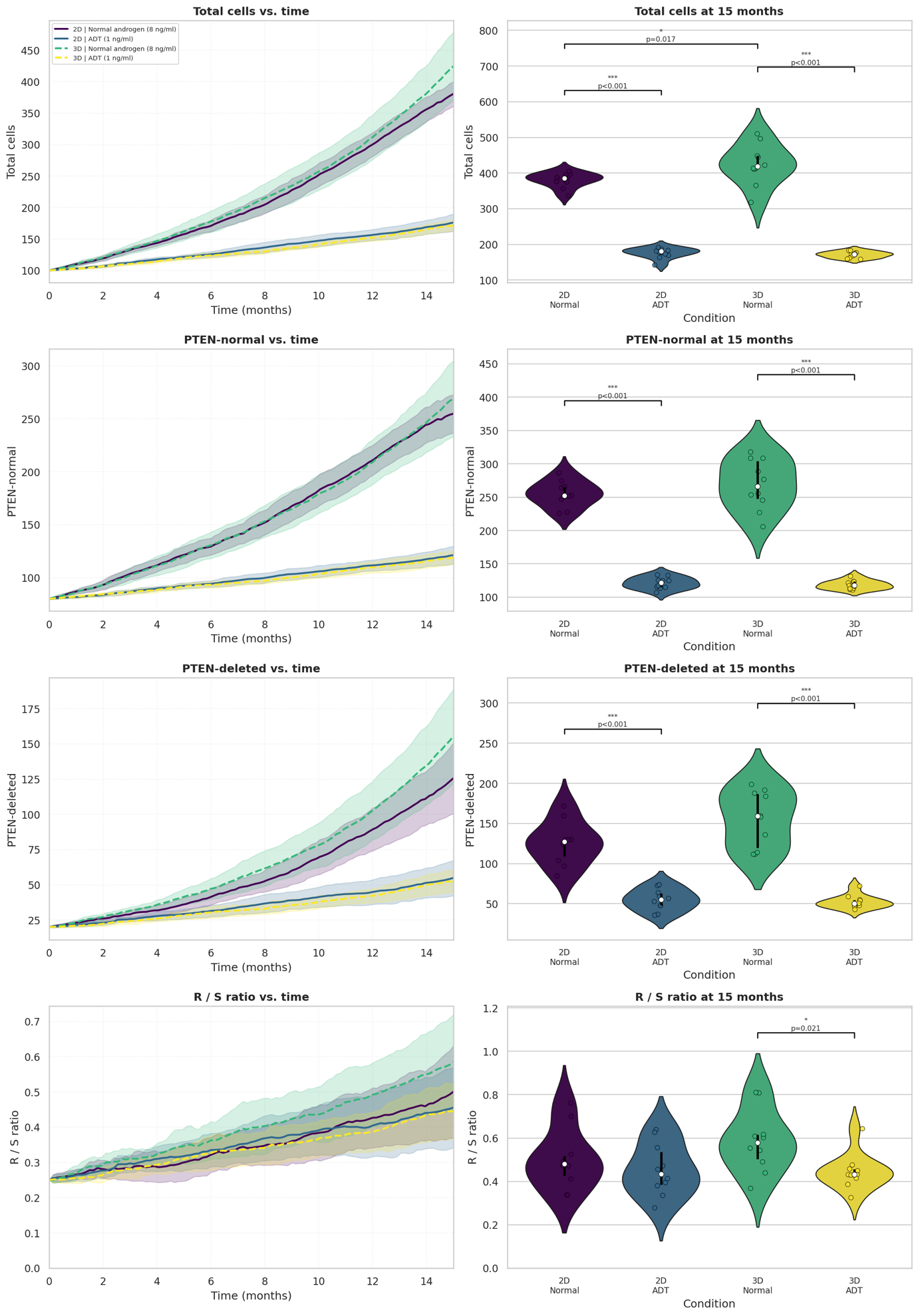

Fig S7.4.1 Baseline BR cohort simulations. Each row shows one metric (total cells, PTEN-normal, PTEN-deleted, R/S ratio). Left: temporal evolution (mean $\pm$ 1 SD) for all four conditions; blue = normal androgen (8 ng/ml), orange = ADT (1 ng/ml), solid = 2D, dashed = 3D. Right: distributions at $t=15$ months across all conditions. Violin plots show distributions over 10 random seeds; brackets indicate Mann-Whitney U test results for adjacent condition pairs (${}^{*}p<0.05$, ${}^{**}p<0.01$, ${}^{***}p<0.001$, ns = not significant).

### S8. PSA Trajectory Validation Against Individual Patient Data

Sections S1 - S5 establish the MHS model structure, cohort parameterization, and internal validation of the two-population cell fate logic. Here we assess whether the model produces PSA trajectories that are consistent with longitudinal clinical data from individual prostate cancer patients treated with ADT, a necessary condition for the population-level cohort comparisons in the main text. The patient-specific androgen-sensitivity ratios derived in Section S8.2 are model outputs characterizing androgen dependence; they are not validated against these patients' ADT outcomes, which were not available.

#### S8.1 Patient Cohort and Simulation Designs

Clinical data were drawn from the EUREKA1 prospective registry ^23^, which enrolled high-risk prostatectomized patients defined by Gleason score ≥ 8 and positive surgical margins. From 84 patients who underwent radical prostatectomy (RP) and received adjuvant hormone therapy for ≥ 6 months with no radiotherapy or neo-adjuvant therapy, five experienced biochemical recurrence during adjuvant therapy. Of these, two had an initial post-surgical PSA below the clinical threshold of 0.2 ng/mL and were therefore selected as recurrence cases for validation. A non-recurrent patient from the same registry served as the control as described in the main text. Since longitudinal PSA values were available for these 3 cases, we used them for clinical validation of model predicted PSA trajectories. (Detailed in S8.3)

Additionally, there were 14 prostatectomized patients in the trial with gene expression data available. These patients were classified according to their differential gene expression profiles and their model predicted NCGs were analyzed across different simulation conditions. (Detailed in S8.2).

#### S8.2 Individual Gene Expression Profiles Stratify Patients into Biologically Distinct Response Groups

Gene expression data were available for 14 prostatectomized patients. For each patient, the top 14 differentially expressed genes relative to a healthy control subject were identified, and the initial concentrations of the corresponding MHS model species were adjusted to reflect that patient's expression profile. Patients for whom no genes were differentially expressed relative to control (DGEP-N) and those with at least one differentially expressed gene (DGEP-Y) were retained as two naturally defined patient groups for comparison. Representative expression profiles for the four DGEP-Y patients are shown in Table 2; color coding indicates that patients with similar differential expression patterns cluster together, suggesting that patient-specific expression data could, in principle, support data-driven cohort stratification beyond the binary PTEN-based grouping.

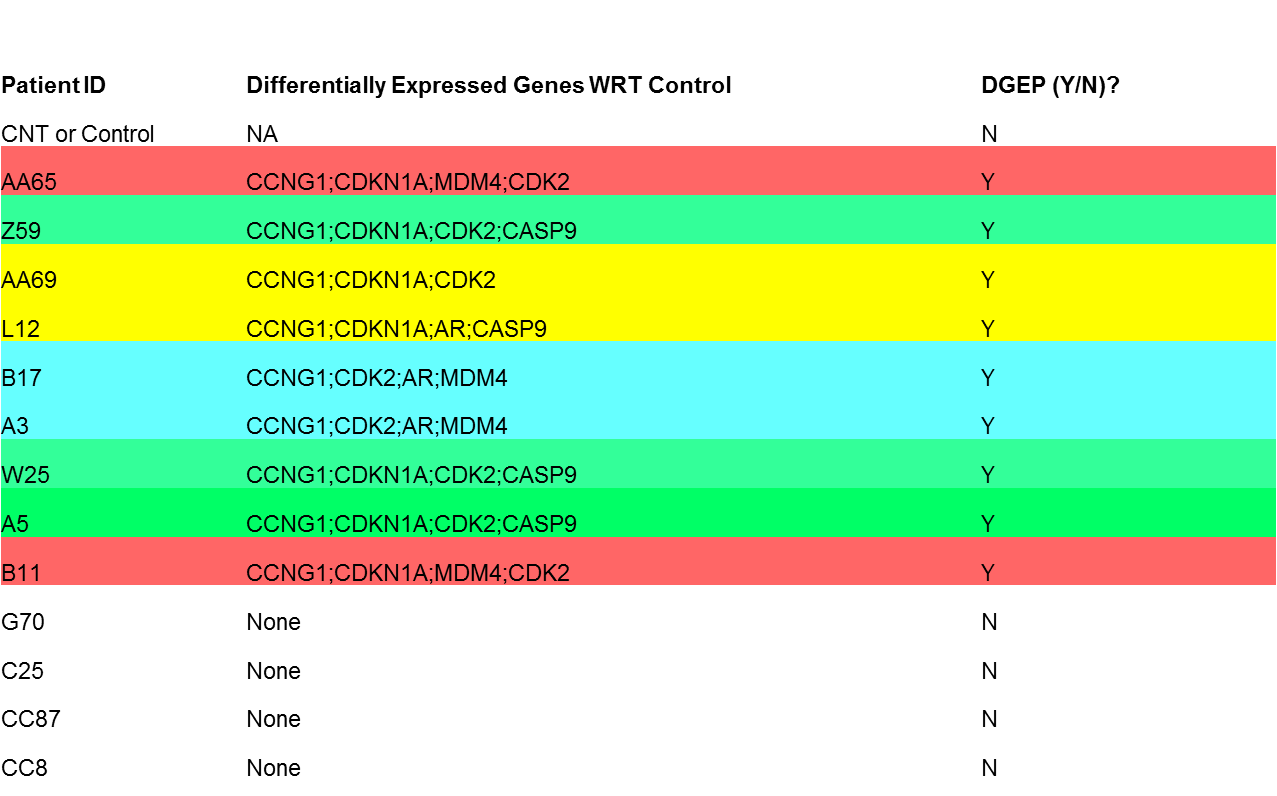

**Table S8.2.1** Patient gene expression profiles showing top differentially expressed genes compared to a control (healthy) subject without cancer. The colors represent patients with similar patterns of differentially expressed genes. Patients shown in same color can in principle be stratified into a cohort. For the purposes of this study, we pool all patients with differentially expressed genes into one cohort. These are indicated DGEP=Y in the last column. Thus, two patient cohorts we consider here are represented by DGEP=Y and DEGP=N.

For each patient and the healthy control, the MHS molecular sub models were run across 20 simulation conditions (two PTEN expression levels × five testosterone concentrations × two intra-tumoral heterogeneity states; see Methods and Table S3.1), yielding individualized distributions of cell growth rate, cell kill rate, and NCG for both S and R cell populations (Figure S8.2.1). DGEP-N patients produced NCG profiles closely matching the healthy control, consistent with the absence of expression-level perturbations to the molecular network. DGEP-Y patients produced divergent NCG profiles that differed both from the control and from each other, reflecting the distinct consequences of their individual expression perturbations on the AR - PI3K/AKT - p53 signaling architecture. (Figure S8.2.1).

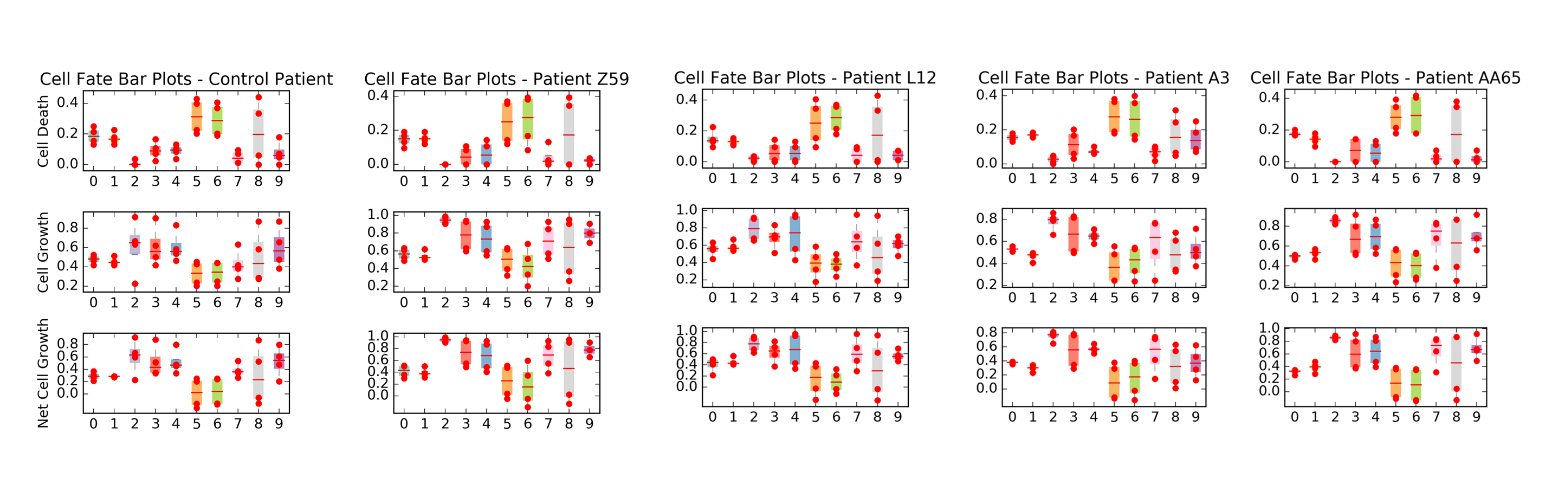

Figure S8.2.1. Patient-specific predictions of cell kill rate, growth rate, and net cell growth (NCG) for the healthy control and four representative DGEP-Y patients. Conditions 0 - 9 correspond to the ten combinations of PTEN level and testosterone concentration (Table S3.1 A); the individual data points per condition correspond to intra-tumoral heterogeneity states (Table S3.1 B). Means across heterogeneity states are shown for each condition. Color coding corresponds to the patient groups identified in Supplementary Table S8.2.1.

#### S8.3 PSA Trajectory Validation Against Longitudinal Clinical Data: Results for Patients of EUREKA trial

For each patient, three simulation scenarios were run: (1) untreated tumor growth (no ADT); (2) ADT in the presence of a pre-existing resistant clone (R cells present at therapy initiation); and (3) ADT with no pre-existing resistant clone (S cells only at therapy initiation). Serum testosterone dynamics were supplied by an external ICCS pharmacokinetic model.^1^ Because individual patient-level molecular data regarding tumor composition was not available, simulations were run to account for presence or absence of resistant subtype. This generated a family of curves for each of the scenarios account for simulations runs over different conditions to mimic tumor heterogeneity.

The two recurrent patients (Figure S8.3.1 a, b) were best described by the scenario in which a substantial resistant clone was present prior to ADT initiation (scenario 2), consistent with the selection model of CRPC emergence adopted by the MHS framework. The non-recurrent patient (Figure S8.3.1c) was best described by the scenario in which no pre-existing resistant clone was present (scenario 3), with PSA declining monotonically under sustained ADT. Simulated PSA envelopes encompassed the observed longitudinal PSA values for all three patients (Figure S8.3.1). These results confirm that the two-population MHS model produces clinically plausible PSA dynamics under ADT and that the distinction between pre-existing resistant and sensitive-only initial conditions is the key determinant of recurrence trajectory, consistent with the central hypothesis of the main text.

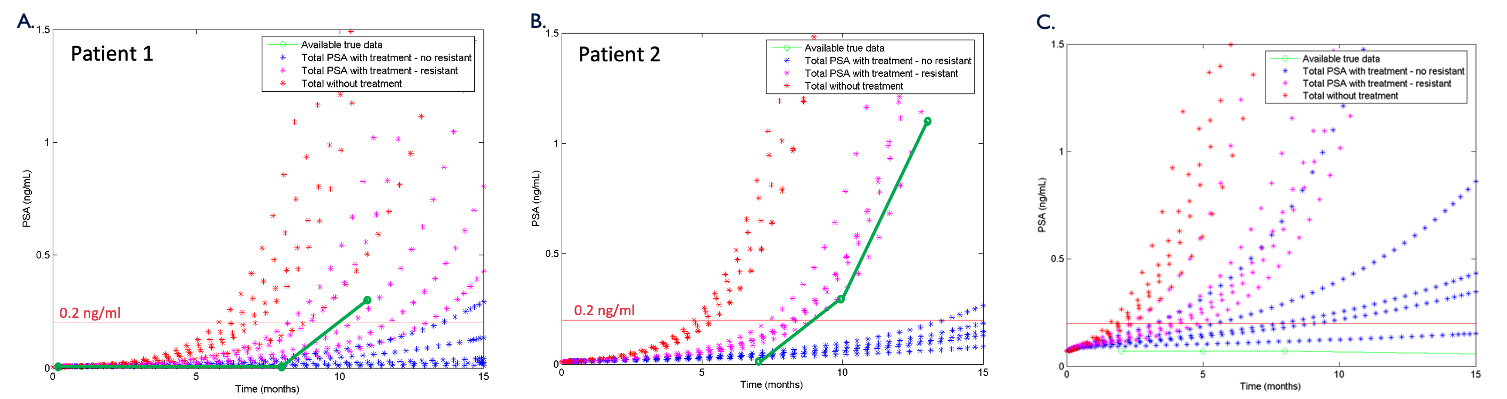

**Figure S8.3.1.** Simulated individual patient PSA trajectories under three treatment scenarios for (a) recurrent patient 1, (b) recurrent patient 2, and (c) non-recurrent control. Red: untreated growth; magenta: ADT with pre-existing resistant clone; blue: ADT with sensitive-only initial condition. Green dots: observed EUREKA1 patient data.

### References

1	Ghosh, A. *A Heterogeneous And Multiscale Modeling Framework To Develop Patient-Specific Pharmacodynamic Systems Models In Cancer* Doctor of Philosophy thesis, University of Pennsylvania, (2019).

2	AA, S., AE, Y. & NL, W. Androgen receptors in hormone-dependent and castration-resistant prostate cancer - PubMed. *Pharmacology & therapeutics* **140** (2013 Dec). <https://doi.org/10.1016/j.pharmthera.2013.07.003>

3	Lonergan, P. E. & Tindall, D. J. Androgen receptor signaling in prostate cancer development and progression. *Journal of Carcinogenesis* **10** (2011 Aug 23). <https://doi.org/10.4103/1477-3163.83937>

4	F, A. *et al.* Expression of androgen receptor is negatively regulated by p53 - PubMed. *Neoplasia (New York, N.Y.)* **9** (2007 Dec). <https://doi.org/10.1593/neo.07769>

5	(Open Access) A comprehensive analysis of coregulator recruitment, androgen receptor function and gene expression in prostate cancer. (2017) | Song Liu | 61 Citations. *eLife* **6** (2017-08-18). <https://doi.org/10.7554/ELIFE.28482>

6	Cronauer, M. V. *et al.* Inhibition of p53 function diminishes androgen receptor-mediated signaling in prostate cancer cell lines. *Oncogene 2003 23:20* **23** (2004-04-29). <https://doi.org/10.1038/sj.onc.1207346>

7	Kase, A. M., III, J. A. C. & Tan, W. Novel Therapeutic Strategies for CDK4/6 Inhibitors in Metastatic Castrate-Resistant Prostate Cancer. *OncoTargets and therapy* **13** (2020 Oct 15). <https://doi.org/10.2147/OTT.S266085>

8	WW, C. *et al.* Input-output behavior of ErbB signaling pathways as revealed by a mass action model trained against dynamic data - PubMed. *Molecular systems biology* **5** (2009). <https://doi.org/10.1038/msb.2008.74>

9	M, C., J, S., SH, J., X, C. & KH, C. Attractor landscape analysis reveals feedback loops in the p53 network that control the cellular response to DNA damage - PubMed. *Science signaling* **5** (11/20/2012). <https://doi.org/10.1126/scisignal.2003363>

10	Y, Z. & AS, G. Adaptation or selection--mechanisms of castration-resistant prostate cancer - PubMed. *Nature reviews. Urology* **10** (2013 Feb). <https://doi.org/10.1038/nrurol.2012.237>

11	Lotan, T. L. *et al.* PTEN Protein Loss by Immunostaining: Analytic Validation and Prognostic Indicator for a High Risk Surgical Cohort of Prostate Cancer Patients. *Clinical Cancer Research* **17** (2011/10/15). <https://doi.org/10.1158/1078-0432.CCR-11-1244>

12	Kholodenko, B. N., Demin, O. V., Moehren, G. & Hoek, J. B. Quantification of Short Term Signaling by the Epidermal Growth Factor Receptor *. *Journal of Biological Chemistry* **274** (1999/10/15). <https://doi.org/10.1074/jbc.274.42.30169>

13	C, H. & CV, H. Studies on prostatic cancer. I. The effect of castration, of estrogen and androgen injection on serum phosphatases in metastatic carcinoma of the prostate - PubMed. *CA: a cancer journal for clinicians* **22** (1972 Jul-Aug). <https://doi.org/10.3322/canjclin.22.4.232>

14	JD, M., A, P., RA, E., JD, N. & Y, K. Mechanisms of resistance to intermittent androgen deprivation in patients with prostate cancer identified by a novel computational method - PubMed. *Cancer research* **74** (07/15/2014). <https://doi.org/10.1158/0008-5472.CAN-13-3162>

15	Marino, S., Hogue, I. B., Ray, C. & Kirschner, D. A methodology for performing global uncertainty and sensitivity analysis in systems biology. *Journal of Theoretical Biology* **254** (2008). <https://doi.org/10.1016/j.jtbi.2008.04.011>

16	JY, C., JR, L., KA, C. & T, M. A two-dimensional ERK-AKT signaling code for an NGF-triggered cell-fate decision - PubMed. *Molecular cell* **45** (01/27/2012). <https://doi.org/10.1016/j.molcel.2011.11.023>

17	Nukpezah, J. *Unlocking Cancer Insights: A Data-Driven Approach to Unravel Genomic Variations in Kinome and Genomic and Proteome Remodeling by Mechano-Chemical Signals* Doctor of Philosophy thesis, Unviersity of Pennsylvania, (2025).

18	A, G., R, H., SH, F., SM, M. & P, M. PhysiCell: An open source physics-based cell simulator for 3-D multicellular systems - PubMed. *PLoS computational biology* **14** (02/23/2018). <https://doi.org/10.1371/journal.pcbi.1005991>

19	Ghaffarizadeh, A., Friedman, S. H. & Macklin, P. BioFVM: an efficient, parallelized diffusive transport solver for 3-D biological simulations. *Bioinformatics* **32** (2015 Dec 12). <https://doi.org/10.1093/bioinformatics/btv730>

20	DR, G., C, K., K, B. & M, P. A method for estimating the oxygen consumption rate in multicellular tumour spheroids - PubMed. *Journal of the Royal Society, Interface* **11** (01/15/2014). <https://doi.org/10.1098/rsif.2013.1124>

21	TM, S. *et al.* Differential Expression of OATP1B3 Mediates Unconjugated Testosterone Influx - PubMed. *Molecular cancer research : MCR* **15** (2017 Aug). <https://doi.org/10.1158/1541-7786.MCR-16-0477>

22	Larhed, A. W., Artursson, P., Gråsjö, J. & Björk, E. Diffusion of drugs in native and purified gastrointestinal mucus. *Journal of Pharmaceutical Sciences* **86** (1997/06/01). <https://doi.org/10.1021/js960503>

23	Gabriele, D. *et al.* Eureka-1 database: an epidemiogical analysis. *Minerva Urol. Nefrol.* **1**, 9-15 (2015).
